# Structure-activity mapping of the peptide- and force-dependent landscape of T-cell activation

**DOI:** 10.1101/2021.04.24.441194

**Authors:** Yinnian Feng, Xiang Zhao, Adam K. White, K. Christopher Garcia, Polly M. Fordyce

## Abstract

Adaptive immunity relies on T lymphocytes that use αβ T-cell receptors (TCRs) to discriminate amongst peptides presented by MHC molecules (pMHCs). An enhanced ability to screen for pMHCs capable of inducing robust T-cell responses could have broad applications in diagnosing and treating immune diseases. T cell activation relies on biomechanical forces to initiate triggering of the TCR. Yet, most *in vitro* screening technologies for antigenic peptides test potential pMHCs for T cell binding without force and thus are often not predictive of activating peptides. Here, we present a technology that uses biomechanical force to initiate T cell triggering in high throughput. BATTLES (<u>B</u>iomechanically-<u>A</u>ssisted <u>T</u>-cell <u>T</u>riggering for <u>L</u>arge-scale <u>E</u>xogenous-pMHC <u>S</u>creening) displays candidate pMHCs on spectrally encoded ‘smart beads’ capable of applying physiological loads to T cells, facilitating exploration of the force- and sequence-dependent landscape of T-cell responses. BATTLES can be used to explore basic T-cell mechanobiology and T cell-based immunotherapies.

## INTRODUCTION

The adaptive immune response relies on the ability of T cells to sensitively discriminate non-self from self, allowing mammals to detect pathogen infection or malignant transformations. At a molecular level, this process is mediated by αβ T cell receptors (TCRs) that recognize specific peptides presented by major histocompatibility complex (MHC) molecules expressed on antigen presenting cells (APCs). T cell recognition of a pMHC molecule via its TCR triggers downstream signaling events (*e.g.* Ca^2+^ flux) that ultimately regulate the activation state of the T cells. The parameters that determine whether a given pMHC antigen will bind and trigger T cell activation have been extensively studied, but important questions remain about the inter-relationship between binding and TCR triggering. An enhanced understanding of these molecular mechanisms, as well as peptide screening methods that simultaneously probe both of these parameters, would have broad applications for immune and infectious disease diagnosis and treatment.

Currently, it is not possible to predict which peptide sequences will engage and activate a particular TCR. *In vivo*, T cell detection of antigenic peptides is exquisitely sensitive, capable of identifying <10 non-self/altered peptides (Huang et al., 2013; Purbhoo et al., 2004) on infected or transformed cells (Keskin et al., 2015). This detection is also highly specific, effectively distinguishing peptides that differ by only a single amino acid (Sasada et al., 2000). By contrast, measured *in vitro* 3D affinities for known stimulatory peptides are surprisingly weak (K_d_ ~ μM) and often do not differ significantly from non-stimulatory peptides (K_d_, _non-self_ <10 K_d_, _self_) (Huang et al., 2010). In addition, measured *in vitro* affinities often do not accurately correlate with T-cell activity (Huang et al., 2010; Lyons et al., 1996; Stone et al., 2009). Many high-affinity interactions occur with high frequency in the human T cell repertoire yet do not stimulate a T cell response (Sibener et al., 2018), while many of the most potent stimulatory peptides bind with only weak or moderate affinities (Corse et al., 2010; Ueno et al., 2004). Together, these paradoxical results establish that equilibrium binding is not sufficient to explain T cell recognition *in vivo*.

Recent evidence suggests that the pN to nN biomechanical forces generated at TCR-pMHC interfaces during T-cell immunosurveillance and synapse formation may be critical for sensitive and specific recognition (Feng et al., 2018). For stimulatory peptides, single TCR-pMHC complex bond lifetimes increase with loads up to an optimal force (characteristic for ‘catch bonds’); for non-stimulatory peptides, bond lifetimes shorten as force increases (characteristic for ‘slip bonds’) (Das et al., 2015; Liu et al., 2014; Sibener et al., 2018). Distinguishing agonist peptides (‘catch’) from non-agonist peptides (‘slip’) requires an *in vitro* assay in the presence of applied loads that quantifies not just binding but downstream activation.

A variety of *in vitro* assays have been designed to identify peptides with potent stimulatory potential. Affinity-based screening approaches (*e.g.* DNA-barcoded pMHC multimer screening (Bentzen et al., 2016; Bentzen et al., 2018; Zhang et al., 2018) and pMHC yeast display (Birnbaum et al., 2014)) assay 10^3^-10^9^ different sequences, facilitating deep exploration of the vast potential sequence space. However, these assays quantify binding alone (and not activation), take place in the absence of force, and present peptides at artificially high local concentrations, reducing their physiological relevance. Synthetic biology approaches that co-culture engineered T cell lines with peptide-pulsed APCs directly assess cellular responses (Joglekar et al., 2019; Kula et al., 2019; Li et al., 2019), yet peptides remain displayed at artificially high densities. By contrast, low-throughput mechanobiology techniques (*e.g.* traction force microscopy (Bashour et al., 2014; Basu et al., 2016; Dustin and Groves, 2012; Hui et al., 2015; Liu et al., 2016a), optical trapping (Feng et al., 2017; Wei et al., 1999), atomic force microscopy (Hu and Butte, 2016), biomembrane force probes (Husson et al., 2011; Li et al., 2010; Liu et al., 2014), and optomechanical actuator nanoparticles (Liu et al., 2016b)) can exert external well-calibrated shear loads on T cells interacting with specific pMHCs displayed at low densities. However, these techniques typically require extensive costly equipment and are labor intensive, restricting measurements to one or a limited number of peptide sequences.

Here, we present a novel technology (BATTLES, for <u>B</u>iomechanically-<u>A</u>ssisted <u>T</u>-cell <u>T</u>riggering for <u>L</u>arge-scale <u>E</u>xogenous-pMHC <u>S</u>creening) that profiles T cell signaling responses for thousands of cells interacting with different pMHCs at low densities and in the presence of physiological shear loads in a single experiment. To test and validate the BATTLES technology, we profiled responses of two T cell lines (TCR589 and TCR55) previously shown to bind an HIV pol-derived peptide (IPLTEEAEL, HIVpol). BATTLES measurements for >11,000 TCR589 and TCR55 cells interacting with 21 peptides (Sibener et al., 2018) correctly recapitulated known responses for TCR589 and identified an alternate synthetic peptide sequence (VPLTEDALL) capable of stimulating TCR55 cells with as few as three TCR-pMHC interactions at the interface. By using BATTLES to systematically vary the magnitude of the applied force and the density of presented peptides, we further establish that TCR55 cells interacting with this alternate synthetic peptide exhibit catch bond-like behavior with the strongest response at 20-30 pN/s loads, and that observed specificities decrease with increasing pMHC densities. Finally, we apply BATTLES to profile responses of a third clinically relevant TCR (DMF5) previously shown to interact with two different peptide classes (Riley et al., 2018). These model systems establish BATTLES as a new technology suitable for obtaining deeper insight into the structure-activity relationships of TCR triggering and discovering novel pMHC antigens under conditions of physiological force.

## RESULTS

To apply well-calibrated shear forces, BATTLES deposits T cells onto the surface of hydrogel ‘smart beads’ bearing pMHCs that swell upon small changes in temperature (**Figure 1A**). To monitor downstream signaling responses via high-throughput single-cell microscopy, T cells and ‘smart beads’ are loaded into microwell arrays in the presence of a Ca^2+^-sensitive dye (**Figure 1B**). By linking pMHCs to spectrally encoded ‘smart beads’ with a 1:1 relationship between the presented peptide sequence and embedded spectral code, each BATTLES experiment can profile responses of ~1000-3000 cells interacting with > 20 peptide sequences.

**Figure 1.**
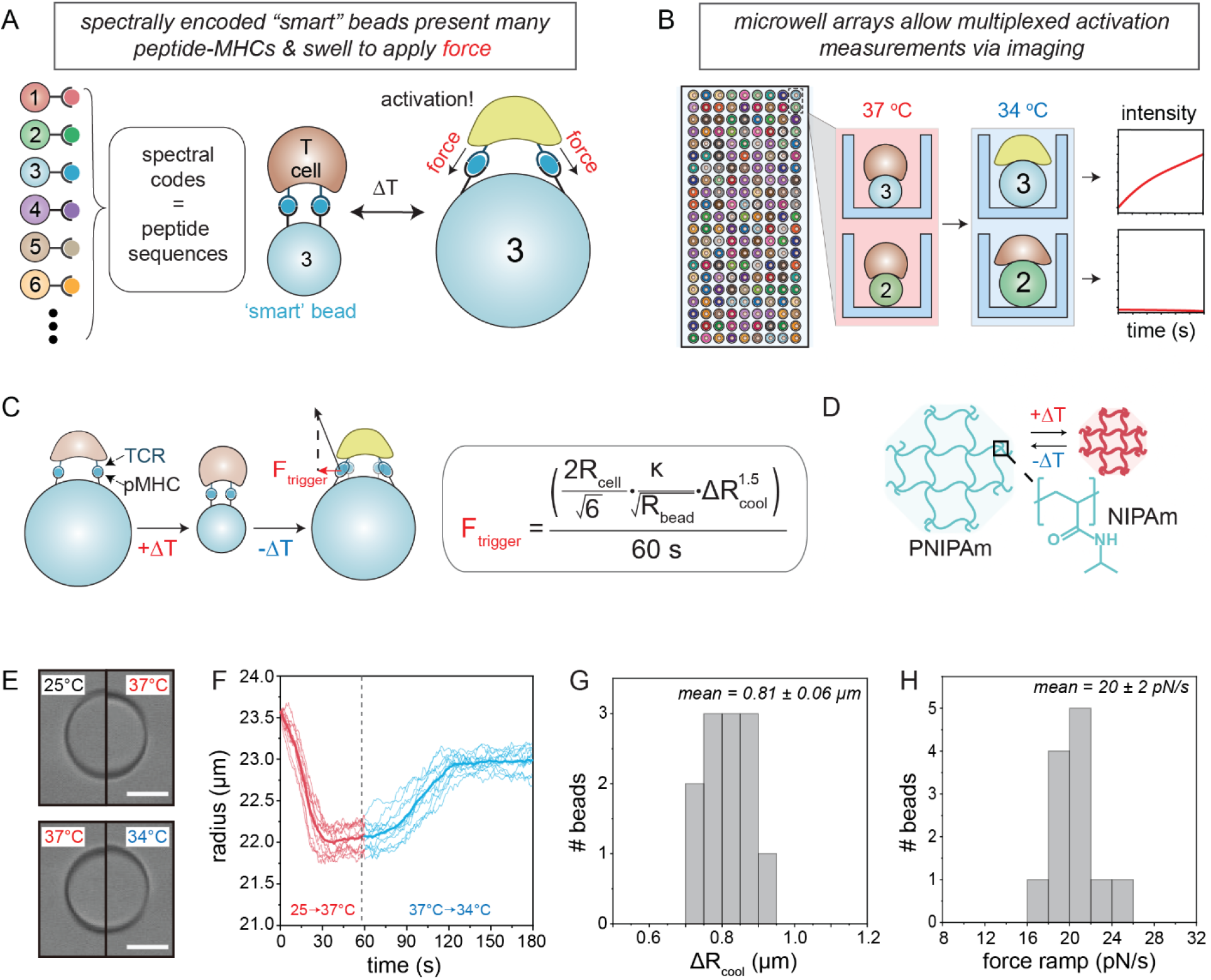
Overview of BATTLES assay and ‘smart beads’. **(A)** BATTLES applies shear forces to pMHC/TCR interactions at cell surfaces via thermally responsive ‘smart beads’ that are spectrally encoded (with a 1:1 linkage between embedded spectral code and presented pMHC sequence, presented pMHC density, or the magnitude of the applied load). **(B)** By loading T cells onto the surfaces of ‘smart beads’ loaded into microwell arrays, BATTLES allows for high-throughput single-cell measurement of T cell responses (cellular Ca^2+^ flux) via microscopy after the application of force. (**C**) Schematic and formula detailing how thermal bead expansion applies force to pMHC/TCR interactions at cell surface; *R_cell_* is the T cell radius (~ 4 μm), *K* is the modulus of rigidity of the ‘smart bead’, *R_bead_* is the radius of the bead prior to heating, and Δ*R_bead_* is the change in bead radius upon cooling from 37°C to 34°C. Force ramps are calculated assuming beads swell over 60 s after cooling. (**D**) Cartoon schematic of polymeric ‘smart bead’ matrix. (**E**) Representative bright field images showing the change in radius for a ‘smart bead’ upon heating (25 to 37 °C) and cooling (37 to 34 °C). Scale bar: 25 μm. (**F,G**) Measured bead radii (F) and changes in radii (G) over time for 12 beads upon heating and cooling. (**H**) Calculated expansion forces for 12 beads.

### ‘Smart’ beads can apply well-calibrated loads to single cells

Stimulus-responsive polymers have been widely used to probe mechanobiology within bulk tissues and for single cells (Kim and Hayward, 2012), including T cells (Liu et al., 2016b). However, the forces applied by these ‘smart’ materials have been difficult to quantify due to heterogeneity over large distances (Chandorkar et al., 2019). Moreover, while T cell crawling takes place over second-to-minute timescales and μm-scale distances, these materials typically apply near-instantaneous forces (within ms) at nm-scale (Liu et al., 2016b). ‘Smart beads’ that shrink and swell can apply homogeneous expansion forces (Ding et al., 2016; Lau et al., 2002) (**Figure 1C**), with the shear load applied at bead surfaces given by **Eqn. 1**:

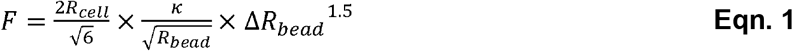

where *R*_cell_ is the T cell radius (~4 μm for the SKW3 cell line used here), κ is the modulus of rigidity, *R*_bead_ is the starting bead radius, and Δ*R*_bead_ is the radius change of the bead. ‘Smart beads’ can be generated from temperature-responsive polymers such as poly(N-isopropylacrylamide) (PNIPAm), which changes size at a critical temperature (**Figure 1D**) that can be tuned to a physiological range (e.g. 34-35 °C) by adding hydrophilic comonomers (e.g. acrylic acid (AAc) or sodium acrylate (SAc)) (Burmistrova et al., 2011). Here, we generated monodisperse ‘smart beads’ (radius = 16.10 ± 0.4 μm, mean ± SD) by using a single-layer microfluidic droplet generator (Feng et al., 2020) (**Figure S1A**; **Movie S1**) to produce NIPAm droplets containing 55 mM SAc (**Figure S1B**) and then polymerized these droplets into solid beads via batch exposure to UV light. After polymerization and swelling in aqueous buffer, this process yielded ~ 23.75 μm radius beads (CV = 6.5%) at room temperature (**Figure S1C**).

To quantify changes in radii as a function of temperature, we loaded 12 beads onto an indium tin oxide (ITO) glass slide and imaged while raising the temperature to 37°C for 60 s and then allowing the temperature to ramp down to 34°C over 120 s (**Figure 1E**; **Movie S2**). ‘Smart bead’ radii shrank by 1.49 ± 0.16 μm during the first 30 s after heating and then expanded by 0.81 ± 0.06 μm after cooling (**Figures 1F** and **1G**). After 60 s, the thermo-response reached equilibrium and the bead radius remained constant. A 37°C to 34°C temperature change has only minimal effects on early immune cell activation (Hanson, 1993; WoldeMussie et al., 1986), suggesting signaling responses should reflect the response to force alone.

Next, we measured the modulus of rigidity (κ) for a hydrogel slab comprised of identical concentrations of NIPAm monomer and SAc (**Figure S2A**). The measured modulus was ~3 kPa (**Figure S2B**), similar to rigidities for epithelial cells that support CD8+ cytotoxic T cell crawling (4.6 ± 2.2 kPa) (Wu et al., 2018) and lymph nodes (~4 kPa) (Meng et al., 2020). Combining measured radii, changes in radii, and moduli yields estimated ramping forces of 20 ± 2 pN/s (**Figure 1H**), in good agreement with physiological forces applied by T cells (estimated to be ~21.5 pN/s (Hui et al., 2015) with forces applied over minutes (Huse, 2017)).

### Spectrally encoded ‘smart’ beads allow simultaneous testing of many potential antigenic sequences

Developing a quantitative and predictive biophysical understanding of how peptide sequence dictates T cell signaling requires the ability to systematically vary presented sequences while assessing downstream responses. We previously demonstrated that poly(ethylene glycol) diacrylate (PEG-DA) hydrogel beads can be spectrally encoded via the incorporation of different ratios of lanthanide nanophosphors (Lns) (**Figure 2A**) and that bead surfaces can be functionalized for downstream chemical coupling (MRBLEs, for <u>M</u>icrospheres with <u>R</u>atiometric <u>B</u>arcode <u>L</u>anthanide <u>E</u>ncoding) (Feng et al., 2020; Gerver et al., 2012; Nguyen et al., 2017).

**Figure 2.**
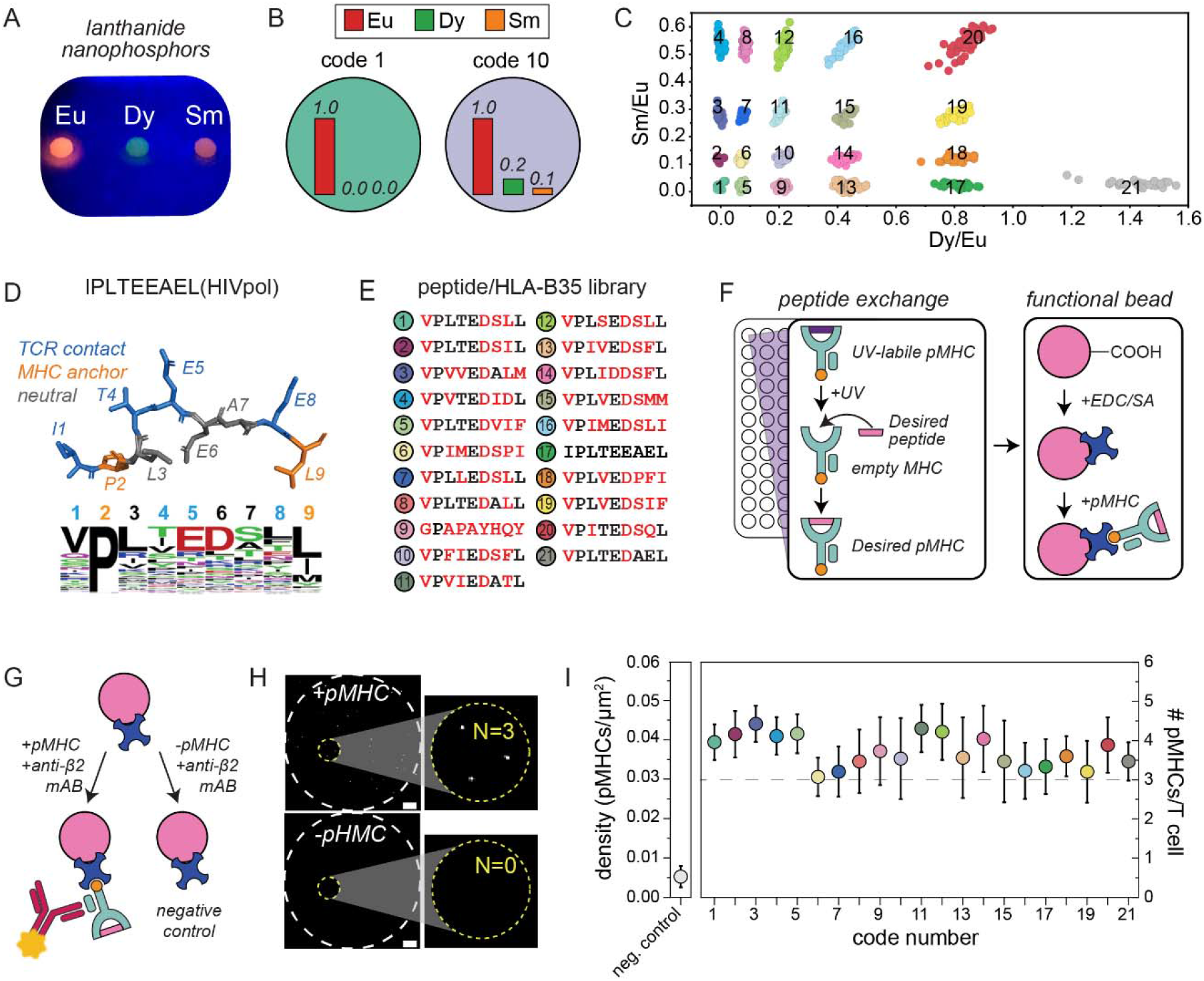
‘Smart beads’ can display 21 different peptide sequences at physiological densities. (**A**) Image of polymer droplets containing 3 different Ln species (Eu, Dy, and Sm) upon excitation with a UV LED lamp (wavelength = 285 nm). **(B)** Cartoon schematic showing 2 example ratiometric codes. **(C)** Measured median Ln ratios (colored markers) for 872 beads containing equal amounts of Eu and 21 different combinations of Sm and Dy along with the expected target ratios. (**D**) Structural schematic and peptide sequence for 21 peptides previously shown to bind TCR55 cells in yeast display experiments^20^; blue and orange residues denote TCR and MHC anchor contacts, respectively. (**E**) Sequences of HIVpol-derived peptides coupled to ‘smart beads’ containing particular spectral codes. (**F**) Schematics detailing UV-exchange method to generate biotinylated pMHCs bearing desired peptides within the MHC groove (left) and coupling of biotinylated pMHCs to ‘smart beads’ bearing streptavidin covalently attached to carboxyl groups via EDC chemistry (right). (**G**) Schematics detailing the quantification of the interfacial densities of presented pMHC molecules via TIRF microscopy using fluorescently labeled antibodies to detect peptide-loaded MHC complexes. (**H**) TIRF images for an example pMHC-coated bead (code 8, +pMHC) and an example negative control bead (−pMHC). White dashed circles denote the bead circumference settled on the slide surface (diameter ~ 50 μm); yellow circles denote the size of an average T cell; numbers indicate identified pMHCs within yellow circles. (**I**) Estimated surface density (left) and number of pMHCs/cell (right) from surface density measurements across all ‘smart bead’ codes; markers denote means and error bars denote standard deviation. The dashed line represents the physiological number of pMHCs per cell (Henrickson et al., 2008).

To test if thermally-responsive PNIPAm is compatible with spectral encoding, we generated carboxylated PNIPAm beads containing a constant amount of Europium (Eu) and varying amounts of Dysprosium (Dy), and Samarium (Sm) (**Figure 2B; Table S1**), pooled all codes prior to imaging, and analyzed images using previously developed software (Harink et al., 2019). Per-bead Sm/Eu and Dy/Eu ratios revealed 21 distinct clusters consistent with expected target ratios (**Figure 2C**). Cluster variances were similar to those for PEG-DA beads (Gerver et al., 2012) (**Figure S3A**), with a coding capacity of 35, 48, 80, or 143 codes for clusters separated by 6, 5, 4, or 3 standard deviations, respectively (**Figures S3B** and **S3C**).

### ‘Smart’ beads can display peptides at low physiological densities to mimic *in vivo* conditions

Applying shear forces to bead-interacting T cells under physiological conditions requires that pMHC complexes be conjugated to beads at low densities. To accomplish this, we covalently coupled streptavidin (SA) proteins to carboxyl groups via EDC chemistry. After SA coupling, codes remained easily distinguishable from one another (**Figure S4A**); incubation of SA-coated and uncoated beads with ATTO-647-labeled biotin further established that SA coupling was uniform and specific (**Figures S4B-S4D**).

We then synthesized a library of peptides containing systematic mutations within an HIVpol-derived peptide (Pol448–456, IPLTEEAEL) that was previously used to test the effects of amino acid substitutions at positions P1 and P3-9 on T cell binding via yeast display (**Figure 2D**). These 21 peptides had previously measured EC_50_ values from 1.2 μM to > 100 μM and contain systematic substitutions at TCR-contacting (P1, P4, P5, and P8) and neutral residues (P3, P6 and P7) (**Figure 2E**; **Table S2**).

To display peptides on ‘smart beads’, we loaded peptides of interest in the B35 MHC groove of biotinylated pMHC complexes via UV-facilitated peptide exchange in solution (Rodenko et al., 2006) and then incubated biotinylated pMHC complexes with SA-coated beads (**Figure 2F**). All UV exchanges and bead coupling reactions took place in separate volumes to uniquely link peptide sequences with embedded spectral codes. To quantify the actual density of interfacial pMHC molecules, we incubated beads bearing pMHC complexes with fluorescently-labeled anti-β2 monoclonal antibodies (which recognize the β2 microglobulin within pMHCs successfully loaded with peptides) and imaged bead surfaces via total internal reflection fluorescence (TIRF) microscopy (**Figure 2G**). As hydrogel beads are compliant, beads settled on coverslip surfaces to yield an observable spot ~50 μm in diameter (**Figure 2H**), with individual antibody-labeled interfacial pMHCs appearing as scattered fluorescent punctae. Over 10 beads (from spectral code 8), we observed 29 ± 9 pMHCs per ~840 μm^2^ of selected bead surface area, corresponding to an expected maximum of 3-4 pMHCs per 100 μm^2^ T cell surface (**Figure S5A**) (2πr^2^, assuming a T cell (r = 4 μm) spreading when interacting with a surface and then retracting (Fritzsche et al., 2017)), with significantly fewer pMHCs for negative control beads (4.4 ± 1.8 per selected bead surface area) (**Figure S5B**). Interfacial densities were uniform across codes (~0.03 to 0.045 molecules/μm^2^, corresponding to 3-4.5 molecules/T cell) (**Figure 2I**).

Attributing differences in measured activation to changes in sequence requires that applied force be constant across codes. For all 21 codes, mean bead radii and changes in radii upon cooling ranged from 23.0 to 24.6 μm and 0.78-0.9 μm, respectively (**Figures S6** and **S7**); hydrogel rigidity across different Ln ratios also remained fairly constant (κ = 2.48 − 3.15 kPa) (**Figure S8**). Together, these measurements yield estimated force ramping rates of 20-27.5 pN/s over ~ 60s (**Figure S9**), differing by < 27% across codes.

### Pairing T cells with ‘smart’ beads for high-throughput monitoring of force- and sequence-dependent activation

Early T cell activation is associated with elevations in intracellular free calcium (Freedman, 1979), making it possible to monitor TCR responses towards a given peptide using a calcium-sensitive dye (Brazin et al., 2018; Feng et al., 2017; Hu and Butte, 2016; Husson et al., 2011; Kim et al., 2009; Li et al., 2010; Liu et al., 2014; Liu et al., 2016a; Liu et al., 2016b; Ma et al., 2008). To allow high-throughput imaging of many individual T cells in parallel upon the application of shear force to pMHC-TCR interactions, we fabricated a device containing 1440 microwells, each ~50 μm in diameter and ~56 μm in depth, that allowed sequential loading of a single ‘smart bead’ within each well followed by 1 or multiple T cells on top (**Figures 3A** and **3B; Movie S3**). After an initial incubation to allow cells to interact with and attach to bead-bound pMHCs, we increased the device temperature to 37°C for 1 min followed by cooling to 34°C for 2 mins; subsequent imaging of Ca^2+^ flux via Cal-520 fluorescence for bead-associated cells over 10 min (4 s intervals) allowed direct monitoring of single T cell signaling. After each experiment, we imaged beads across lanthanide channels to identify the embedded spectral code and thus the displayed peptide sequence (**Figure S10**).

**Figure 3.**
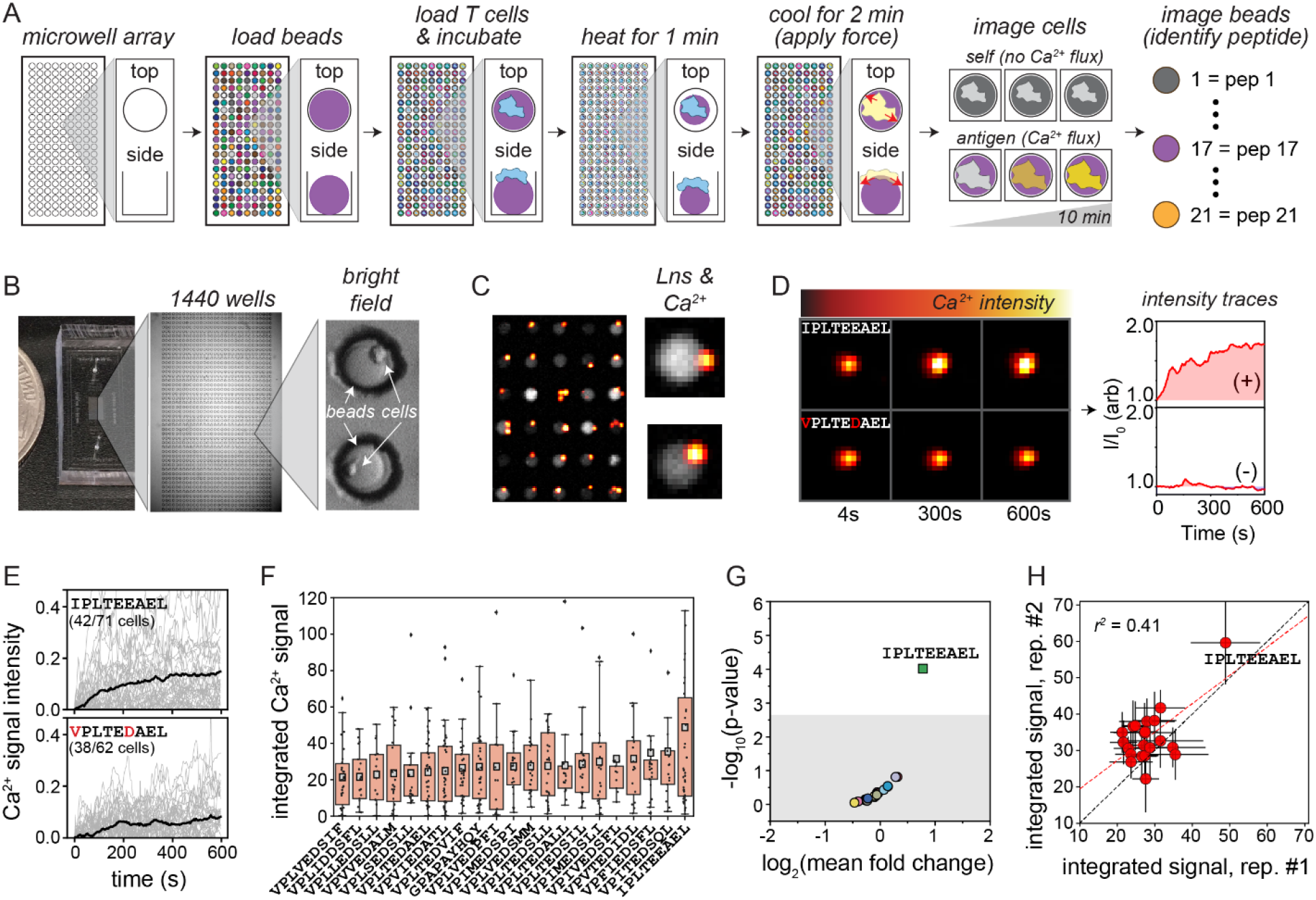
Activation responses for TCR589-transduced T cells interacting with 21-peptide library. (**A**) Cartoon schematic of BATTLES workflow and expected results. (**B**) Representative bright field images of the 1440 well microwell device containing ‘smart beads’ and loaded T cells. (**C**) Representative merged lanthanide and fluorescence images of ‘smart beads’ and associated T cells in the presence of a Ca^2+^-sensitive dye. (**D**) Representative images (left) and processed intensity traces (right) quantifying Ca^2+^ flux within single TCR589 T cells interacting with pMHC-coated ‘smart beads’ bearing the stimulatory HIVpol peptide (IPLTEEAEL) (top) and a nonstimulatory peptide (VPLTEDAEL) (bottom). (**E**) Ca^2+^ signal intensity as a function of time for all positive single cells (∫(I/I_0_-1) > 0) (light grey); mean signal intensity over time is shown in black. (**F**) Integrated Ca^2+^ signals for all positive single cells as a function of peptide sequence. Individual cell signals are shown as black markers; box lower and upper limits indicate 25^th^ and 75^th^ percentiles, respectively. Grey squares represent the mean values. (**G**) Estimated p-value (calculated via bootstrapping, see Methods) *vs*. the log2-transformed mean fold change for integrated Ca^2+^ signals for each peptide sequence. Grey box indicates Bonferroni-corrected p-value at a significance of 0.05 (p = 0.0024). (**H**) Integrated Ca^2+^signals for each peptide across 2 replicates. Markers indicate mean; error bars indicate SEM; dashed black line indicates the 1:1 line; red dashed line indicates a linear regression.

### BATTLES correctly identifies peptide agonists known to drive cytotoxic killing *in vivo*

To demonstrate BATTLES, we profiled two HLA-B35-HIV(Pol_448-456_) (HIVpol)-specific TCRs (TCR589 and TCR55) previously shown to bind HIVpol pMHCs at μM affinities (K_D_ = 4 μM and 17 μM, respectively) but with divergent downstream responses (Sibener et al., 2018). While TCR589-transduced SKW-3 T lymphoblastic leukemia cells bound HIV-pol tetramers and secreted IL-2 in a dose-dependent manner, the same cell line transduced with TCR55 showed ~4x weaker binding to HIVpol but failed to secrete detectable IL-2. If BATTLES can reliably identify stimulatory peptides, we would therefore expect to see HIVpol-induced Ca^2+^ flux for TCR589.

To test this, we quantified Ca^2+^ flux for TCR589-transduced T cells interacting with all 21 HIVpol-derived bead-bound pMHCs (**Figure 2C; Table S2**). Brightfield images demonstrated loading of ‘smart beads’ within all microwells (**Figure 3B**), and movies acquired during device heating and cooling established that cooling swelled beads as expected and that T cells remained in contact with bead surfaces throughout (**Figure 3B; Movie S4**). After cooling, fluorescence images of force-induced Ca^2+^ flux for 961 cells across all peptides (66 ± 18 beads and 46 ± 15 cells per peptide, **Figure S11**) revealed a variety of behaviors (**Figures 3C–3E**). While some T cells exhibited substantial and rapid increases in intracellular calcium (typical of a type-α calcium response indicating successful TCR triggering) (**Figure 3D**, **top**), most cells showed decreasing fluorescence intensities over time, reflecting an absence of triggering (type-β calcium response) combined with photobleaching (**Figure 3D**, **bottom; Figure S12A**). This is consistent with prior literature suggesting that only a fraction of cells are activated at low pMHC densities, even under optimal force (Ma et al., 2008).

To probe for sequence-dependent differences in the proportion of triggered cells and amplitude of type- α calcium responses, we integrated per-cell fluorescence over time and then analyzed only ‘positive’ cells that increased in fluorescence (**Eqn. 2**):

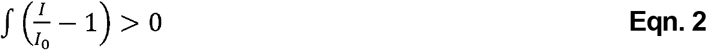

For TCR589 cells, the HIVpol peptide (IPLTEEAEL) yielded the highest percentage of ‘positive’ T cell responses (**Figure S12B; Movie S5**) and ‘positive’ cells had larger integrated intensity signals (~2-fold greater than for other peptides (**Figures 3E–3G**, **S13;** p = 10^−4^ as determined via bootstrapping analysis against all other peptide sequences, see Methods). Results were consistent across technical replicates (**Figures 3H**, **S14** and **S15;** r^2^ = 0.41), establishing that BATTLES can identify peptide epitopes capable of producing a robust cytotoxic response *in vivo*.

### Applying BATTLES to identify novel peptide agonists

Next, we tested whether BATTLES could identify novel likely potent agonists for TCR55-tranduced T cells that bind HIVpol without activation (**Figures 4A** and **S16;** 67 ± 24 beads and 51 ± 23 cells per peptide). In contrast with TCR589, the HIVpol peptide IPLTEEAEL did not induce significant Ca^2+^ responses; instead, a mutant version of this peptide (VPLTEDALL) produced the strongest functional response across 2 full biological replicates in which we shuffled the identity of the peptide associated with each spectral code (**Figures 4B–4D, S17**-**S19**; bootstrapped p = 10^−5^ and 8×10^−4^ for each replicate; r^2^ = 0.62 between replicates).

**Figure 4.**
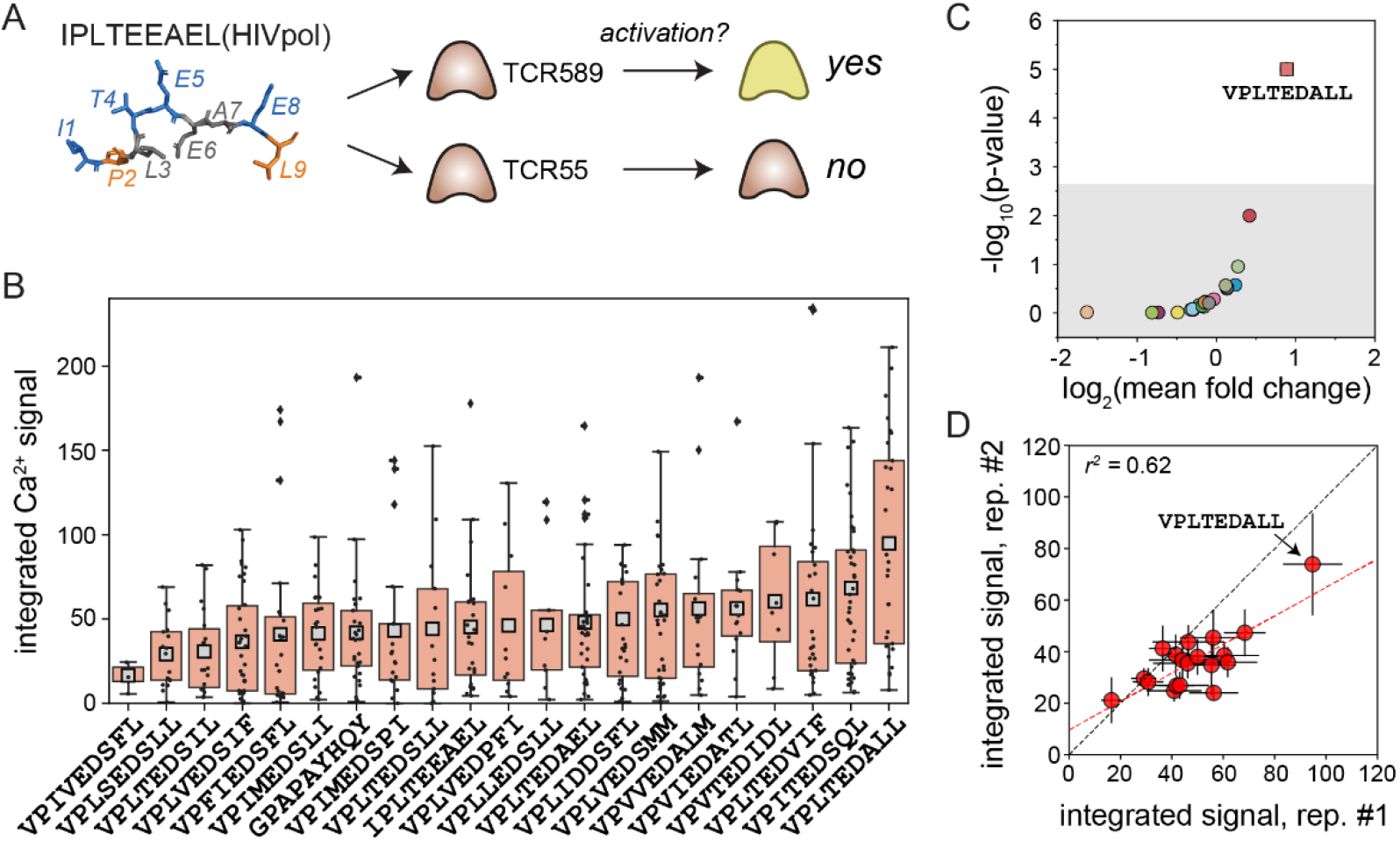
Activation responses for TCR55-transduced T cells interacting with 21 peptide library. (**A**) Schematic illustrating previously observed divergent cellular reactivity responses for TCR55 *vs*. TCR589 interacting with the HIVpol-derived peptide. (**B**) Integrated Ca^2+^ signals for positive cells across all peptide sequences tested. Integrated signals for individual cells are shown as black markers; box lower and upper limits represent 25^th^ and 75^th^ percentiles, respectively. Grey squares represent the mean values. (**C**) Estimated p-value (calculated via bootstrapping, see Methods) *vs*. the log2-transformed mean fold change for integrated Ca^2+^ signals for each peptide sequence; Grey box indicates Bonferroni-corrected p-value at a significance of 0.05 (p = 0.0024). (**D**) Integrated Ca^2+^signals for each peptide across 2 replicates. Markers indicate mean; error bars indicate SEM; dashed black line indicates the 1:1 line; red dashed line indicates a linear regression.

To confirm that observed Ca^2+^ responses required force and were not simply due to changing temperatures, we repeated the TCR55 BATTLES assay after either heating beads to 34 °C without first heating to 37 °C or heating to 37 °C without subsequently reducing temperatures to 34 °C (**Figures S20A** and **S20B**). In both cases, integrated Ca^2+^ signals across tested peptides were 2-fold lower and a lower percentage of cells yielded positive integrated Ca^2+^ signals (**Figure S20C**); in addition, no peptide produced a response significantly higher than the others. Although some shrinking force may be applied during heating to 37 °C, the directions of the applied force and cell contractions (~6 nN after attachment (Hui et al., 2015)) are aligned at interaction surfaces such that the net applied force is negligible.

### Multiplexing applied loads can elucidate ‘catch’ *vs* ‘slip’ bond behavior in a single experiment

For ‘catch’ bonds (found in TCR-agonist pMHC interactions), bond lifetimes increase with applied force up to an optimal load (typically 8-20 pN) and then decrease; for ‘slip’ bonds (found in TCR-self pMHC interactions), bond lifetimes monotonically decrease with applied load (Das et al., 2015; Liu et al., 2014; Sibener et al., 2018) (**Figure 5A**). If the VPLTEDALL peptide serves as a true potent agonist for TCR55-transduced T cells, we would expect to see the strongest triggering at an optimal force, with weaker responses at higher or lower forces. To test this, we leveraged spectrally encoded ‘smart beads’ to simultaneously vary both the peptide sequences presented and forces applied to pMHC-TCR interactions within a single experiment.

**Figure 5.**
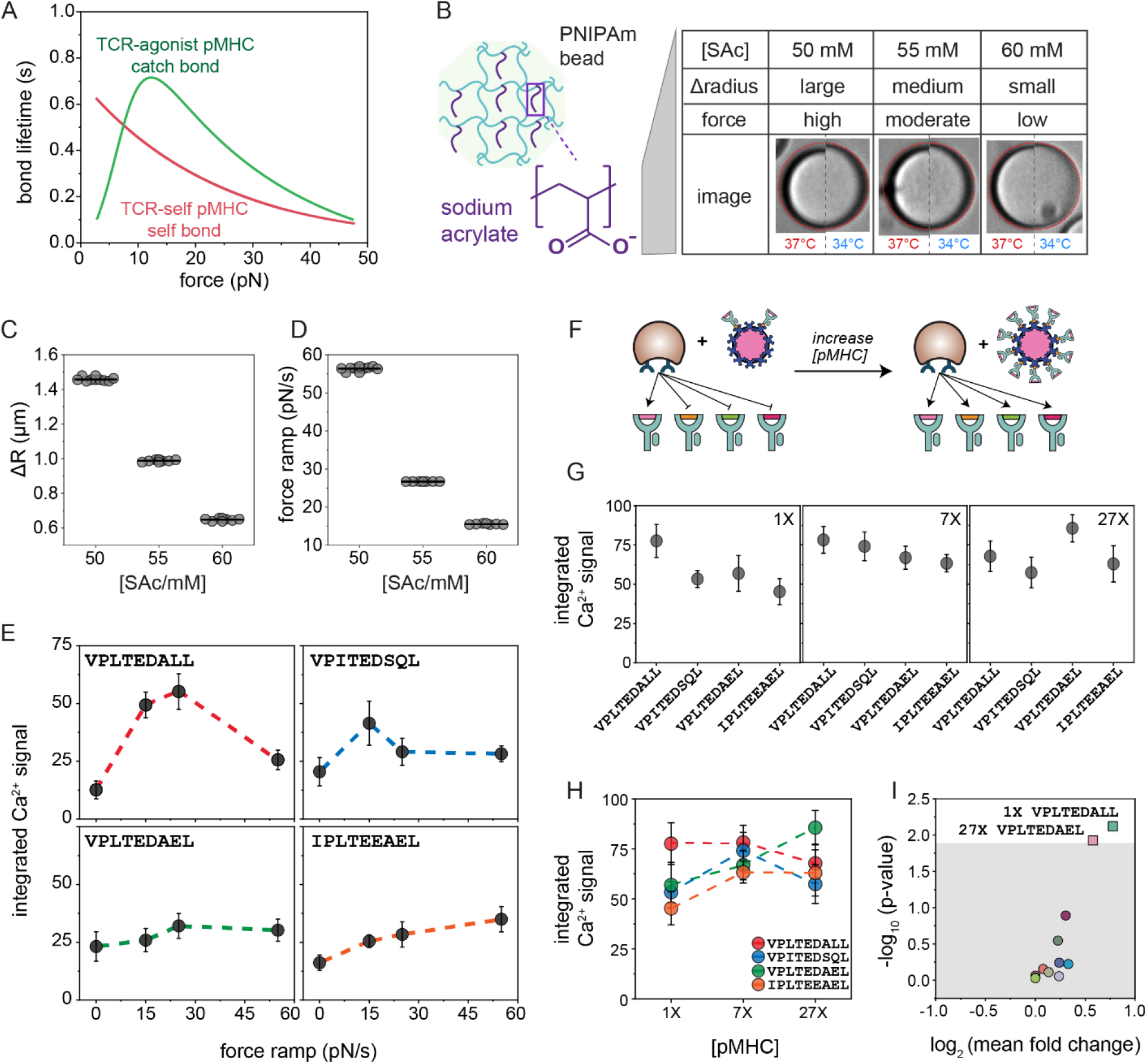
Multiplexing sequence, applied force, and displayed pMHC densities using BATTLES. (**A**) Typical catch/slip bond lifetime profiles. (**B**) Schematic and images indicating that change in bead radius depends on the amount of sodium acrylate (SAc) within the ‘smart bead’ matrix. (**C**) Measured changes in bead radius upon cooling from 37°C to 34°C for 3 codes (9, 6, and 6 beads/code) containing different SAc concentrations. (**D**) Calculated expansion forces for ‘smart beads’ in **C** based on measurements of bead radii, changes in bead radii, and the rigidity of the NIPAM slab. (**E**) Mean integrated Ca^2+^ signals for positive cells interacting with VPLTEDALL, VPITEDSQL, VPLTEDAEL and IPLTEEAEL peptides at low (~15 pN/s), moderate (~25 pN/s) and high (~55 pN/s) force ramps. Zero force data is from a TCR55 control experiment in which temperature was maintained at 34°C. (**F**) Schematic indicating loss of TCR specificity at higher pMHC concentrations. (**G**) Mean integrated Ca^2+^ signals for positive cells interacting with VPLTEDALL, VPITEDSQL, VPLTEDAEL and IPLTEEAEL peptides at 1X, 7X and 27X pMHC concentrations. (**H**) Summarized integrated Ca^2+^ signals for four selected peptides under three pMHC concentrations. (**I**) Estimated p-value (calculated via bootstrapping, see Methods) *vs*. log2-transformed mean fold-change of integrated Ca^2+^ signals for 4 peptides at 3 different pMHC concentrations; grey area represents p-value>0.013 (Bonferroni-corrected p-value at a significance of 0.05). Error bars in **E**, **G** and **H** indicate SEM.

Incorporating different amounts of SAc within the ‘smart bead’ matrix alters the relative hydrophilicity/hydrophobicity ratio and thus the change in radius upon temperature-induced swelling and shrinking, tuning the magnitude of the applied force (**Figure 5B**). To apply a range of forces, we polymerized ‘smart beads’ containing 50 mM, 55 mM, or 60 mM SAc, respectively (**Figure 5B**). After polymerization, the radius of polymerized beads was fairly constant for all 3 formulations (24.12 +/− 1.45 μm, 23.68 +/− 1.55 μm, and 24.08 +/− 1.66 μm, mean ± SD) but the magnitude of the change in radius upon cooling from 37°C to 34°C varied substantially (1.44 μm (34 beads, CV = 1.5%), 0.96 μm (24 beads, CV = 5.4%), and 0.64 μm (23 beads, CV = 2.5%)) (**Figures 5B**, **5C** and **S21; Movie S6**). As the modulus of rigidity is unchanged by small changes in SAc concentration (Burmistrova et al., 2011), these variations yielded ramping forces of 54.43 ± 1.52 pN/s, 26.96 ± 2.37 pN/s, and 15.57 ± 0.62 pN/s for the 3 formulations (**Figures 5D and S22**).

Prior work using a BFP-based bond lifetime assay established that for TCR55, VPLTEDAEL and VPITEDSQL peptides exhibit maximum lifetimes at 14 pN and 7 pN, respectively, indicating the formation of a ‘catch’ bond, while the HIVpol peptide (IPLTEEAEL) shows ‘slip’ bond behavior (Sibener et al., 2018) (**Figure S23**). Here, we profiled the force-dependent responses for these 3 peptides and the VPLTEDALL peptide shown to drive the largest integrated Ca^2+^ signal in BATTLES (**Figures 4B–4D**) via a BATTLES assay in which each spectral code corresponded to a particular sequence/force combination (4 peptide sequences × 3 forces/sequence = 12 unique codes; 116 ± 50 beads and 127 ± 54 cells per peptide) (**Figures S24; Table S4**). Integrated Ca^2+^ signals as a function of applied load for VPLTEDALL revealed a marked increase in Ca^2+^ flux at low and moderate forces followed by a decrease in Ca^2+^ at high force, consistent with ‘catch’ bond formation (**Figure 5E**). VPITEDSQL triggered the strongest response at low force, consistent with its previously measured catch bond profile, and VPLTEDAEL responses were uniformly low across all forces. Although HIVpol (IPLTEEAEL) exhibited a slight increase in signal as force increased (**Figure 5E**), there was no significant difference between tested forces compared to no force (p=0.06 via bootstrapping). Results were again consistent across an additional full technical replicate in which embedded codes and peptide sequences were shuffled (**Table S4**; **Figures S25** and **S26;** *r*^2^ = 0.80 across replicates).

### Multiplexing pMHC sequences and concentrations to reveal dose-dependent immunogenicity

These BATTLES results suggest that the VPLTEDALL peptide triggers the strongest Ca^2+^ signaling response for TCR55-transduced T cells, at odds with prior co-culture results identifying VPLTEDAEL as the most potent agonist (Sibener et al., 2018) (**Table S3**). One potential explanation for this discrepancy could be the difference in presented pMHC concentrations: peptide-pulsed APCs present peptides at densities > 100 nM (corresponding to ~10-30 pMHCs/T cell at 100 nM (Zehn et al., 2006) and ~313 pMHCs/T cell at 10 μM (Henrickson et al., 2008)) while BATTLES ‘smart beads’ present pMHCs at physiological densities of 3-4.5 pMHCs/cell. To test if signaling responses depend on presented densities, we repeated BATTLES assays with the same 4 TCR55-binding peptide sequences (VPLTEDALL, VPITEDSQL, VPLTEDAEL, and IPLTEEAEL) at 3 effective concentrations (1X (3-4.5 pMHCs/T cell); 7X (21-31.5 pMHCs/T cell); and 27X (81-121.5 pMHCs/T cell)) (4 sequences × 3 concentrations = 12 codes; 116 ± 36 beads and 211 ± 66 cells per peptide) (**Figures 5F**, **S27** and **S28; Table S5**). At 1X concentrations, VPLTEDALL again drove the strongest downstream response (**Figures 5G–5I, S29** and **S30;** bootstrapped p = 0.0075; r^2^ = 0.64 between replicates). For 7-fold higher pMHC densities, the integrated Ca^2+^ signal for VPLTEDALL remained constant but the signals associated with the other 3 peptides increased and showed no significant difference across peptides, indicating a loss of specificity (**Figures 5G–5I**, **S29** and **S30A-S30C**). At 27-fold higher pMHC densities, the integrated Ca^2+^ signal associated with the VPLTEDAEL increased even further to drive the strongest response (**Figures 5G–5I**, **S29** and **S30A-S30C;** bootstrapped p = 0.012 for two replicates), consistent with prior observations from tetramer screening and co-culture assays (Sibener et al., 2018). These results confirm that downstream signaling responses depend on presented density and underscore the importance of screening at physiological pMHC densities for accurate identification of likely potent agonists *in vivo*.

### Understanding the activation potential of TCRs capable of binding multiple distinct peptide classes

Next, we tested a third TCR with a known potential therapeutic role. The DMF5 TCR recognizes the MART-1 melanoma antigen presented by the class I MHC protein HLA-A*0201 and has been used in clinical trials of engineered T cells interacting with MART-1_26-35_ peptide-loaded dendritic cells that demonstrated robust tumor regression (Chodon et al., 2014). However, prior yeast display experiments revealed two distinct classes of peptides that bound the DMF5 TCR: (1) MART-1-like peptides containing a hydrophobic core sequence (GIG in P4-6) and (2) peptides instead containing a highly charged central core (DRG in P4-6),), with DRG class peptides binding in a unique ‘shifted register’ conformation (Riley et al., 2018). Tetramers of the MART-1 anchor-modified dodecamer (ELAGIGILTV) showed the strongest binding and the strongest downstream response in peptide-pulsed co-culture assays while IMEDVGWLNV was the most potent DRG class peptide (**Figure 6A; Table S6**). Predicting whether similar cross-reactivity is likely in therapeutic applications requires the ability to assess activation responses when presented at low densities.

**Figure 6.**
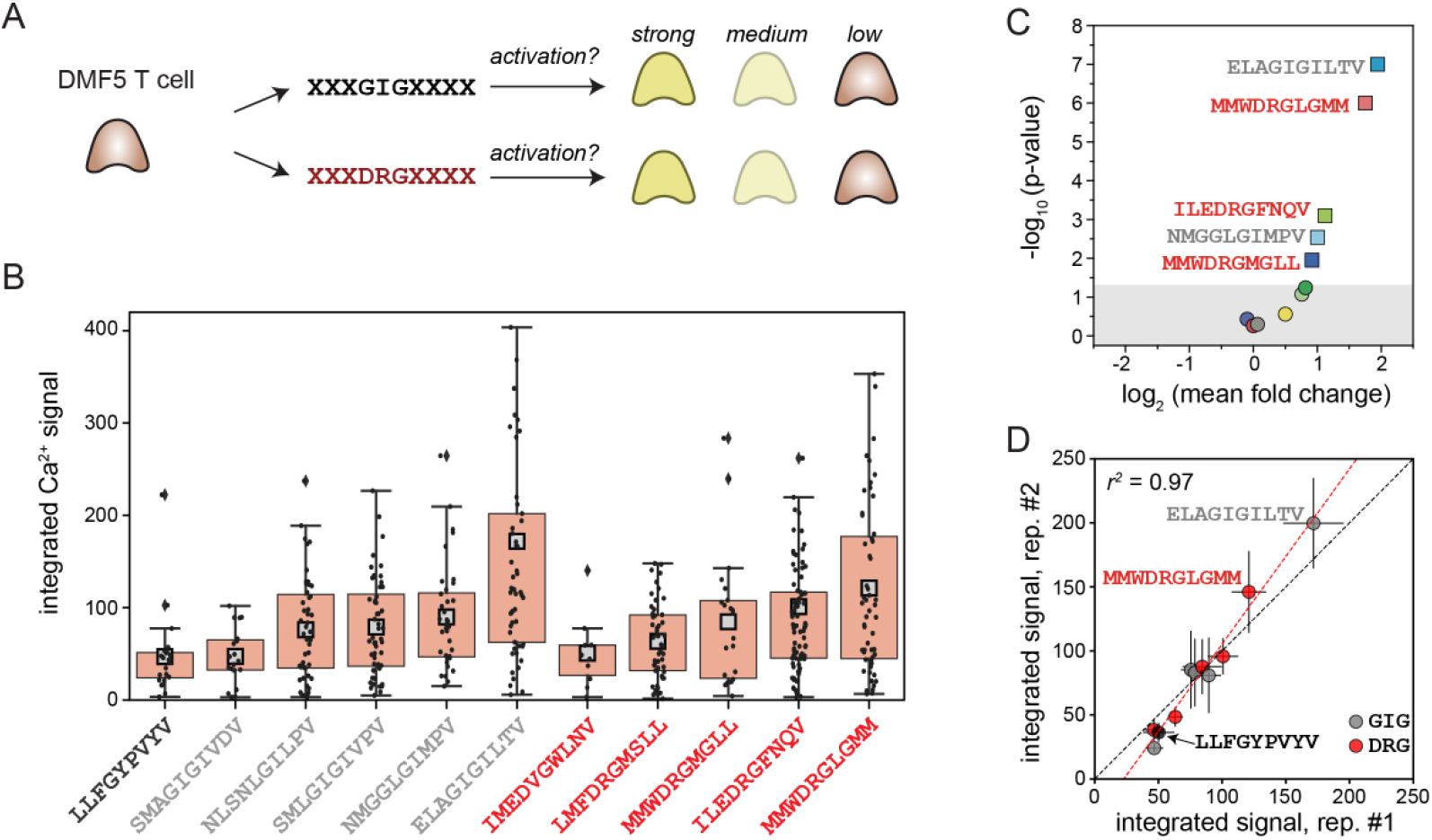
Activation responses for the therapeutically-relevant DMF5 TCR system interacting with 11 peptide sequences from 2 classes. (**A**) Schematic illustrating that DMF5 TCR shows comparable cellular reactivity to two different classes of peptide sequences (GIG and DRG, respectively) in prior co-culture assays. (**B**) Integrated Ca^2+^ signals for positive DMF5-transduced T cells interacting with Tax (LLFGYPVYV, a ‘negative’ control peptide identified from coculture experiments) and 10 additional peptides from the GIG (grey) and DRG (red) classes. Integrated signals for individual cells are shown as black markers; box lower and upper limits represent 25^th^ and 75^th^ percentiles, respectively; grey squares represent mean values. (**C**) Estimated p-value (calculated via bootstrapping, see Methods) *vs*. log2-transformed mean fold change of integrated Ca^2+^ signals; grey area represents p-value > 0.05. (**D**) Mean integrated Ca^2+^ signals for positive DMF5 cells for each peptide across 2 technical replicates. Error bars indicate SEM; dashed black line indicates the 1:1 line; red dashed line indicates linear regression.

To assess reactivities under physiological densities, we applied BATTLES to DMF5 T cells interacting with 11 peptides: 5 GIG class peptides (including the MART-1 anchor-modified dodecamer), 5 DRG class peptides, and a negative control peptide (Tax, LLFGYPVYV) (124 ± 34 beads and 75 ± 29 cells per peptide) (**Figure S31**). Consistent with its known biological role, the MART-1 anchor-modified dodecamer (ELAGIGILTV) drove by far the strongest Ca^2+^ response in DMF5 T cells across two replicates (**Figures 6B–6D, S32, S33** and **S34**; p=10^−7^ and 3×10^−4^ from bootstrapping comparisons with Tax; *r*^2^ = 0.97). From the DRG class, the MMWDRGLGMM (and not the previously identified IMEDVGWLNV) induced the strongest Ca^2+^ flux (p= 10^−6^ and p = 6.5×10^−3^ from bootstrapping comparisons with Tax). Thus, the BATTLES assay can reveal potential cross-reactivity of therapeutical TCRs towards different epitopes, even for motifs sharing no significant similarities, providing clinically relevant information predicting off-target effects.

## DISCUSSION

T cells exert shear forces during immunosurveillance (Huse, 2017) and immune synapse formation on an APC (Grakoui et al., 1999; Wülfing and Davis, 1998). Such physical loads placed on the TCR-pMHC bond can enhance sensitivity and specificity of recognition by facilitating discrimination between ‘catch’ and ‘slip’ bond-forming complexes. Recapitulating efficient T cell activation *in vitro* therefore requires the ability to: (1) display a peptide capable of forming a catch bond with a given TCR, (2) apply optimal forces to drive mechanotransduction, and (3) display pMHCs at sufficiently low densities that applied forces are transmitted to individual intramolecular interactions at the pMHC/TCR interface. While a wide variety of mechanobiology tools have been developed to identify potent peptide agonists (Lei et al., 2020) (as summarized in **Table S7**), the BATTLES platform is the first (to our knowledge) that recapitulates these physicochemical cues (*e.g.* active force and low monomeric pMHC density) and screens many peptide sequences for their potential to not just bind TCRs but also activate downstream signaling responses.

Accurately mimicking physiological conditions requires matching not just the magnitude of applied forces but also the ramp rate and overall duration over which forces are applied. *In vivo*, actin rearrangement is initiated during the first minute of T cell-APC interaction (Ritter et al., 2015), during which cytoskeleton-associated TCRs rapidly form and break bonds with agonist pMHCs. While the estimated force ramp applied by BATTLES ‘smart beads’ (20-30 pN/s, ~1 min duration) is similar to that applied during traction force microscopy (13-30 pN/s) (Hui et al., 2015), optical trapping (30 pN/s (Feng et al., 2017)), and atomic force microscopy (25 pN/s (Husson et al., 2011)), critical differences in how forces are applied may allow BATTLES to more accurately mimic *in vivo* physiology. First, while polystyrene or silica beads used in optical trapping assays distribute applied forces across all formed pMHC/TCR interactions at the interface, ‘smart beads’ apply force locally via pMHCs attached to PNIPAm filaments within the bead hydrogel matrix. This geometry, analogous to interactions between APC-presented pMHCs and protrusive T-cell microvilli prior to activation (Ghosh et al., 2020), may prevent dissipation of intramolecular forces at higher pMHC densities. Second, the duration of the force ramp applied by ‘smart beads’ over 60 s of cooling and associated swelling may allow peptides with a fast 2D on-rate to iteratively rupture and rebind, driving dynamic (Pryshchep et al., 2014) and reversible ~10 nm structural transitions within the TCR/pMHC complex (Das et al., 2016). Such energized work (10 nm transitions under 10-15 pN loads = ~100 to 150 pN·nm or hydrolysis of ~2 ATPs) may lead to a stronger and more sustainable Ca^2+^ flux (Feng et al., 2017) similar to that observed for naïve T cells (Christo et al., 2015). Moreover, this mechanical work has been proposed to exponentially amplify the reaction rate value for antigenic pMHCs (Feng et al., 2018), potentially explaining the observed physiological discrimination ratio of 10,000-100,000 for non-self *vs*. self pMHCs.

The TCR-transduced T cell lines employed in typical high-throughput screens are less sensitive than primary T cells, requiring APC pulsing with high peptide concentrations (~ 1 μM). However, several lines of evidence suggest that peptide sequences identified in screens at relatively high densities may not represent the most promising antigen vaccine candidates. The identity of synthetic peptide sequences capable of stimulating IL-2 cytokine release and CD69 expression for crawling primary T cell blasts (Wolf et al., 2003) displaying 2B4 and 5cc7 TCRs (Birnbaum et al., 2014) depends on the concentration at which they are displayed, consistent with the concentration multiplexing BATTLES results presented here. The ability to accurately identify and discriminate potent agonists presented at low pMHC concentrations even when using TCR-transduced SKW3 T cell lines suggests that BATTLES could be a bona fide proxy to screening primary T cells.

Overall, we anticipate that BATTLES can be applied to a wide range of future questions beyond obtaining a basic understanding the structure-activity relationships and physical mechanisms that drive ‘catch bond’ formation. While here we assay only 21 peptides, lanthanide-based spectral encoding has previously yielded >1,100 codes (Nguyen et al., 2017), suggesting BATTLES can be applied to much larger libraries in the future. In medicine, BATTLES can be used as a platform for investigating the force response of multiple engineered T cells bearing either mutant TCRs or chimeric antigen receptor (CARs) (Gross et al., 1989) as well as to enhance the potency of peptide vaccines to a therapeutic TCR by directly probing a peptide library generated from either pMHC tetramer binding or pMHC yeast display. Finally, BATTLES could be applied as a general mechanobiology tool for a wide range of mechanosensitive systems beyond T cells, ranging from other immune cells to adhesion cells and even neurons (Chen et al., 2017).

## ACKNOWLEDGEMENTS

This work was supported by NIH grants 1DP2GM123641, R01GM107132 and a Stanford Bio-X Interdisciplinary Initiatives seed grant. K.C.G. is an investigator of the Howard Hughes Medical Institute, and is also supported by NIH-5R01AI103867, U19AI057229, Mathers Foundation, and Ludwig Foundation support. P.M.F. is a Chan Zuckerberg Biohub Investigator and acknowledges the support of a Sloan Research Foundation Fellowship. Y.F. is a Cancer Research Institute Postdoctoral Fellow. X.Z is funded by a Stanford Bio-X seed grant. A.K.W was funded by the Natural Sciences and Engineering Research Council of Canada Postdoctoral Fellowship. Part of this work was performed at the Stanford Nano Shared Facilities (SNSF), supported by the National Science Foundation under award ECCS-1542152. The authors would also like to thank Dr. E. Appel and D. Chan for helpful discussions concerning polymers and help with shear modulus measurement, and Dr. Z. Bryant and Dr. P.V. Ruijgrok for help with TIRF microscopy.

## AUTHOR CONTRIBUTIONS

Y.F. conceptualized the platform and validation experiments; X.Z. made all the T cell lines and UV-liable pMHC folding. Y.F., X.Z., and A.K.W. analyzed data; K.C.G and P.M.F. provided funding, resources, mentorship, and project supervision. Y.F., X.Z., A.K.W., K.C.G. and P.M.F. wrote the paper.

