## Supplementary Information for "Structure-activity mapping of the peptide- and force-dependent landscape of T-cell activation"

**STAR*METHODS**

**KEY RESOURCES TABLE**

| REAGENT or RESOURCE | SOURCE | IDENTIFIER |
| --- | --- | --- |
| Antibodies |  |  |
| Anti-human TCR α/β | BioLegend | Cat#306718 |
| PE-Anti-human beta-2 micro-globin | BioLegend | Cat#A17082A |
| beta-2 Microglobulin Monoclonal Antibody (B2M-01), Alexa Fluor 647 | Thermo Fisher | Cat#MA5-18119 |
| Bacteria |  |  |
| *E.coli* DH5α | Thermo Fisher | Cat#18265017 |
| E.coli BL21 (DE3) | Novagen/Millipore | Cat#69450 |
| Chemicals, Peptides, and Recombinant proteins |  |  |
| N-Isopropylacrylamide | Sigma-Aldrich | Cat#415324 |
| Sodium acrylate | Sigma-Aldrich | Cat#408220 |
| Acrylic acid | Sigma-Aldrich | Cat#147230 |
| Poly(ethylene glycol) diacrylate | Sigma-Aldrich | Cat#455008 |
| 1X PBS | Thermo Fisher | Cat#10010023 |
| RPMI 1640 | Thermo Fisher | Cat#72400047 |
| FBS | Sigma-Aldrich | Cat#F2442 |
| Penicillin-Streptomycin | Thermo Fisher | Cat#15140148 |
| HFE-7500 | fisher scientific | Cat#NC1242157 |
| Krytox 157 FSH Oil | Grainger | Cat#35RV65 |
| 4”-Silica wafer | University Wafer | N/A |
| RTV-615 | R.S. Hughes | RTV615 CLEAR 010 |
| Bovine Serum Albumin, Biotinylated | Thermo Fisher | Cat#29130 |
| Cal-520, AM | AAT Bioquest | Cat#21130 |
| Pluronic F-127 | Sigma-Aldrich | Cat#P2443 |
| RPMI 1640, colorless | Thermo Fisher | Cat#11835030 |
| N,N-Dimethylformamide | fisher scientific | Cat#D119 |
| Dichloromethane | fisher scientific | Cat#AA39116M1 |
| Methanol | fisher scientific | Cat#A412 |
| SU-8 2025 | MicroChem | Cat#Y111069 |
| SU-8 2050 | MicroChem | Cat#Y111072 |
| SU-8 developer | MicroChem | Cat#Y020100 |
| Tween-20 | Sigma-Aldrich | Cat#P1379 |
| Atto-647N, biotin | Sigma-Aldrich | Cat#93606 |
| MES | Sigma-Aldrich | Cat#M3671 |
| Boric acid | Sigma-Aldrich | Cat#B0394 |
| N-(3-Dimethylaminopropyl)-N′-ethylcarbodiimide hydrochloride | Sigma-Aldrich | Cat#E6383 |
| Sodium hydroxide | Sigma-Aldrich | Cat#S5881 |
| Lithium phenyl-2,4,6-trimethylbenzoylphosphinate | Sigma-Aldrich | Cat#900889 |
| Flex-T™ HLA-A*02:01 Monomer UVX | BioLegend | Cat#280003 |
| Synthetic peptides | Genescript | N/A |
| Recombinant B35-UV (refold) | This paper | N/A |
| UV-cleavable peptide | Genescript | N/A |
| Commercial kits |  |  |
| FuGene HD | Promega | Cat#E2312 |
| Cell lines |  |  |
| SKW-3 | DSMZ | Cat#ACC 53 |
| KG-1 | ATCC | Cat#CLL-246 |
| LentiX | Takara | Cat#632180 |
| Recombinant DNA |  |  |
| psPAX2 | Addgene | Cat#12260 |
| pMD2.G | Addgene | Cat#12259 |
| pET28a-B35*01 | This paper | N/A |
| pET28a-β-2-microglobulin | This paper | N/A |
| pHR-TCR55α | This paper | N/A |
| pHR-TCR55β | This paper | N/A |
| pHR-TCR589α | This paper | N/A |
| pHR-TCR589β | This paper | N/A |
| pHR-DMF5α-P2A-DMF5β | This paper | N/A |
| Other |  |  |
| Superdex 200 Column | GE Healthcare | Cat#17517501 |
| MonoQ Ion Exchange Column | GE Healthcare | Cat#17-5166-01 |
| PP-Reactor 2 mL with PTFE frit | Biotage | V020TF051 |
| 3K spin column | ThermoFisher | Cat#88512 |
| SONY SH800 Sorter | Sony | N/A |
| Spectral/por dialysis membrane MWCO:6-8 kD | Spectrum Labs | Cat#132675 |
| DEAE-Cellulose | Santa Cruz Biotechnology | Cat#sc-211213 |
| Photo masks | Fineline Imaging | N/A |
| THINKY mixer | THINKY | AR-250 |
| Spin coater | Laurell | WS-650 |
| Catheter punch | SYNEO | CR0350255N20R4 |
| Indium tin oxide (ITO) glass slide | Bioscience Tools | TC-1-100s |
| 48×65 mm Coverslip | fisher scientific | Cat#12-518-210 |
| 150W Plasma cleaner | Femto |  |
| Flood UV | IntelliRay | UV0338 |
| UV-LED | Thorlabs | M285L5 |
| UVP compact and handheld UV Lamp | fisher scientific | UVP95000505 |
| Tygon tubing (0.125” ID ×0.25” OD) | McMaster-Carr | Cat#6516T14 |
| PEEK tubing (0.010” ID × 0.020” OD) | ZEUS | N/A |

**RESOURCE AVAILABILITY**

**Lead contact**

**Material Availability**

This study did not generate new unique reagents.

**Data and code availability**

AutoCAD designs (parallel flow focuser and microwell device) and T cell Ca^2+^ flux data has been deposited to OSF repository (<https://osf.io/xs7zf/>).

**METHOD DETAILS**

**Production of lentivirus**

LentiX cells were seeded in 6-well plates with 0.6 x10^5^ cells per well in 2 mL complete DMEM (ThermoFisher). The day after seeding, for each well, 750 ng lentiviral vector, 500 ng psPAX and 260 ng pMD2G were mixed with 4.5 μL Fugene (Promega) in 100 μL Opti-MEM and incubated at room temperature for 20 minutes. After incubation, this DNA/Fugene mixture was added to the LentiX cells. The lentivirus supernatant was collected 2 or 3 days after the transfection and filtered to remove dead cells.

**Cell line generation**

Lentivirus encoding transfection of TCRα and TCRβ chain genes was generated as described above (Sibener et al., 2018). Briefly, we cloned either the full-length TCRα or TCRβ chain gene into the pHR lentiviral vector. After lentivirus production, 2 mL of the supernatant containing TCR α and β chains with 1:1 expression was used to infect 1 million SKW3 cells. Following infection, we added 1.5 mL fresh completed RPMI to the mixture. After 2 days, we selected for SKW3 cells with surface-expressed TCRαβ using anti-TCR (Biolegend clone IP26) staining and flow cytometry (Sony SH800 sorter).

**Cell culture**

SKW3 cells transduced with TCR clones 55, 589 and DMF5 were cultured in RPMI 1640 GlutaMAX (Thermo Scientific) supplemented with 10% (v/v) fetal bovine serum (FBS) (Sigma-Aldrich), 100 U/ml penicillin, and 100 U/ml streptomycin (Life technology). Prior to performing all experiments, we confirmed that cells had reached the log phase of growth (1.5 million to 2 million cells/mL), as cells in the lag phase may generate insufficient Ca^2+^ flux (<https://osf.io/xs7zf/>).

**Microfluidic device design**

Molding masters for both microfluidic devices used in the paper (the parallel flow focuser for high-throughput generation of the spectrally encoded ‘smart beads’ and the microwell array for on-chip loading of beads and T cells) were designed in AutoCAD (Autodesk). The microfluidic droplet generator was largely similar to our previously published design(Feng et al., 2020) except that the width of orifice channels and all channel heights were changed to 25 μm to yield polymerized beads ~24 µm in diameter. All transparency masks used for fabrication of molding masters were printed at 50K dpi (Fineline Imaging). Designs of all devices are provided as Supplemental Files and in an associated OSF repository (<https://osf.io/xs7zf/>).

**Microfluidic molding master fabrication**

*‘Smart bead’ generator*

We fabricated the molding master for the parallel flow focuser device with a target height of 25 μm by:

(1) Spin-coating SU-8 2025 negative photoresist (MicroChem) (500 rpm for 10 s with acceleration of 133 rpm/s; 3500 rpm for 30 s with acceleration of 266 rpm/s) on a 4-inch test-grade silicon wafer (University Wafer, South Boston, MA);

(2) Soft baking the coated wafer (1 min at 65°C; 5 min at 95°C; 1 min at 65°C);

(3) Exposing the baked wafer using a UV mask aligner (Karl Suss MA6) for 26.8 s at 5.6 mW/cm^2^;

(4) Post-exposure baking (1 min at 65°C; 5 min at 95°C; 1 min at 65°C);

(5) Developing using SU-8 developer (Microchem Corp, Newton, MA) for ~4 min.

*Microwell device for pairing beads and T cells*

For the microwell device, we fabricated the microwell layer and roof layer separately on two 4-inch test-grade silicon wafers. Microwell features were fabricated using SU-8 2050 with a target height of 56 μm by:

(1) Spin-coating SU-8 2050 (MicroChem) (500 rpm for 10 s with acceleration of 133 rpm/s; 2750 rpm for 30 s with acceleration of 266 rpm/s) on a 4-inch test-grade silicon wafer;

(2) Soft baking the coated wafer (2 min at 65°C; 7 min at 95°C; 2 min at 65°C);

(3) Exposing the baked wafer using a UV mask aligner for 31.7 s at 5.6 mW/cm^2^;

(4) Post-exposure baking (2 min at 65°C; 6 min at 95°C; 2 min at 65°C);

(5) Developing using SU-8 developer for ~6 min.

The roof features were fabricated on another wafer using SU-8 2050 with a target height of 100 μm by:

(1) Spin-coating SU-8 2050 (500 rpm for 10 s with acceleration of 133 rpm/s; 1650 rpm for 30 s with acceleration of 266 rpm/s) on a 4-inch test-grade silicon wafer;

(2) Soft baking the coated wafer (5 min at 65°C; 16 min at 95°C; 5 min at 65°C);

(3) Exposing the baked wafer using a UV mask aligner for 41.1 s at 5.6 mW/cm^2^;

(4) Post-exposure baking (4 min at 65°C; 9 min at 95°C; 4 min at 65°C);

(5) Developing using SU-8 developer for ~9 min.

*Silane vapor deposition*

To prevent sticking of PDMS during device fabrication, we treated all wafers with trichloro(1H,1H,2H,2H-perfluorooctylsilane) (PFOS, Sigma-Aldrich) by placing wafers in a vacuum chamber along with an uncapped bottle of PDMS with a few PFOS droplet in the cap, pulling vacuum for 1 min, and then maintaining vacuum (without active vacuum) for an additional 9 minutes.

**Microfluidic molding master fabrication**

All microfluidic devices were fabricated using standard soft lithography protocols(Xia and Whitesides, 1998).

*‘Smart bead’ generator*

Molding masters were used to cast single-layer parallel flow-focuser devices composed of a 1:5 ratio of poly(dimethylsiloxane) crosslinker: base (PDMS, RTV 615, R.S. Hughes). After baking, devices were assembled using the ‘jumper cable’ strategy described previously (Feng et al., 2020).

*Microwell pairing device*

The microwell layer of the pairing device was cast using a 1:5 ratio of PDMS crosslinker (20 g crosslinker, 100 g base). After mixing in a THINKY mixer (3 min mixing followed by 3 min degassing), the PDMS mixture was poured on top of the microwell mold and this mold/PDMS assembly was placed in a vacuum chamber and subjected to vacuum for ~45 minutes to remove any air bubbles trapped inside the features. After vacuum, the PDMS-coated wafer was spun for 2 min at 200 rpm with 133 rpm/s acceleration using a spin coater (Laurell WS-650) to yield a final PDMS slab height of ~400 μm. The mold with PDMS was then baked at 80°C for 25 min in a convection oven to partially polymerize this layer. Then microwell layer was then peeled off from the mold, flipped over, and adhered to a cover glass (48×65 mm, GOLD SEAL No. 1). The roof layer was casted with a 1:10 ratio of PDMS crosslinker (10 g crosslinker, 100 g base), subjected to vacuum for 45 min, and then baked at 80°C for 30 min. The roof layer was then peeled from the wafer, inlet holes were punched using a catheter punch (SYNEO, 0.025” ID × 0.035” OD, Part No: CR0350255N20R4) to fit the outer diameters of PEEK tubing (ZEUS, 0.010” ID × 0.020” OD) and steel blunt pins (New England Small Tube, Part No: NE-1310-02), and aligned on top of the microwell layer. This two-layer device assembly was then baked at 80°C for 14 hours to fully polymerize and bond both layers. Longer baking time is not recommended as that may enhance the hydrophobicity and shrink the microwells.

**Production of lanthanide-encoded ‘smart beads’**

*Aqueous lanthanide mixtures*

For ‘smart bead’ synthesis, N-isopropylacrylamide (NIPAM), poly(ethylene glycol) diacrylate (PEG-DA, Mn=700), sodium acrylate (SAc), acrylic acid (AAc), lithium phenyl-2,4,6-trimethylbenzoylphosphinate (LAP) were purchased from Sigma-Aldrich and used directly without further purification. SAc solution (pH=7, c[Ac]=1 M) was obtained by adding 5M NaOH into 1M AAc solution until the pH reached 7.0. The poly-acrylic acid wrapped lanthanide nanophosphors (Lns) were synthesized as described previously (Feng et al., 2020; Nguyen et al., 2017). Pre-mixed Ln/polymer mixtures containing NIPAM monomer, Lns, PEG-DA, SAc and LAP were generated by varying ratios of three monomer master mixtures each containing different Lns (**Table S1**). The “Eu” “Dy” and “Sm” master mixtures also contained 16.3% v/v YVO4:Tm (50 mg/mL), 16.3% v/v YVO4:Dy (50 mg/mL), 16.3% v/v YVO4:Sm (50 mg/mL), respectively. A total amount of 21.3% v/v Lns maintain an equal amount of hydrophilic content across all the formulas. All master aqueous mixtures in **Table S1** contained purified water with 9.2% w/v NIPAM and 5% v/v YVO4:Eu (50 mg/mL), 2.8437% v/v PEG-DA, and 5.5% v/v SAc solution. For the force multiplex experiment, 5% v/v and 6% v/v SAc solution were added into the master aqueous mixture for high and low force ramps, respectively.

*Droplet generation*

2.5% v/v LAP (39.2 mg/mL in DI water) was added to each solution right before its injection into the droplet generator. Pre-mixed Ln/polymer mixtures and HFE7500 with Ionic Krytox (IK) and 0.05% v/v AAc were flowed into the aqueous inlet and oil inlets of the droplet generator device, respectively, yielding high-throughput production of pre-gel droplets at the flow-focusing nozzle. Droplets (radius=16.1 ± 0.4 μm) were generated with aqueous and oil flow rates of 500 μL/h and 3200 μL/h and then collected through Tygon tubing into a 24-well plate (Thermo Fisher Scientific). ~80 μL running oil was added into the each well prior to the collection of the droplets to prevent evaporation of HFE7500 and resultant droplet breakage.

*Bead polymerization and functionalization*

During droplet generation, the AAc in the oil phase gradually diffuses into the aqueous phase to form a carboxy shell to allow subsequent covalent coupling of streptavidin to the bead surface. For 2 formulas at a time, we applied flood UV light (IntelliRay, UV0338) at 100% amplitude (7” away from the lamp, power= ~50-60 mW/cm^2^) for 2 minutes to induce polymerization and crosslinking of carboxyl groups at interfacial surfaces. After polymerization, beads were transferred into a 2 mL fritz column with ~20 μm pore size (Biotage) and then washed with: 2 mL dimethylformamide (DMF, Thermo Fisher Scientific) for 20 s; 2 mL dichloromethane (DCM, Thermo Fisher Scientific) for 10 s; and 2 mL methanol (Thermo Fisher Scientific) for 20 s. After washing, beads were resuspended in 1 mL PBST buffer for the aqueous EDC chemistry.

**EDC chemistry for streptavidin surface conjugation**

To functionalize ‘smart’ beads with streptavidin, 150 μL of carboxy ‘smart’ beads (~200,000 beads) were washed with 200 μL 0.1 M MES buffer (pH = 4.5) supplemented with 0.01% (v/v) Tween-20 (activating buffer) three times prior to resuspension in 200 μL of activating buffer. Next, 200 μL of a freshly made 2%w/v 1-Ethyl-3-(3-dimethylaminopropyl)carbodiimide (EDC, Sigma-Aldrich) in activating buffer was added to the bead solution. The entire reaction was then incubated for 3.5 hours at room temperature on a rotator with end-over-end mixing (10 rpm). The bead slurry was then washed with 1 mL 0.1 M borate buffer (pH = 8.5) supplemented with 0.01% (v/v) Tween-20 (conjugating buffer) and subsequently resuspended in 400 μL of conjugating buffer. Conjugation of streptavidin (Sigma-Aldrich) was carried out by adding 16 μL streptavidin solution (dissolved in 1X PBS at 1 mg/mL) into the mixture and rotated the whole slurry overnight at 4°C. The reaction was quenched by adding 10 μL of 0.25 M ethanolamine in conjugating buffer to the mixture and rotating for 30 minutes at 4°C. The final product was washed 3 times with PBST buffer and resuspended in 200 μL of the same buffer for subsequent bright field and fluorescence imaging. The efficiency and consistency of streptavidin coupling was checked by incubating 0.5 µL 1 mg/mL Biotin-Atto647 and ~5 µL bead slurry (~500 beads, either streptavidin-functionalized or without streptavidin as a negative control) for 1 h at room temperature and imaging in the Cy5 channel.

**Production of inclusion bodies of B35 MHC heavy chain and human β-2-microglobulin**

The codon-optimized B35 MHC heavy chain gene or human β-2-microglobulin gene was cloned into a pET28a vector. The construct was transformed into BL21 (DE3) *E. coli* and grown over night on LB agar plates supplemented with kanamycin. Single colonies were then picked, used to inoculate 10 mL LB broth containing kanamycin, and then grown overnight at 37°C with shaking at 250 rpm. The 10 mL overnight culture was added into 1 L of LB broth containing kanamycin and grown until OD= 0.5. IPTG was added into the same culture at a final concentration of 1 mM and shaken for another 3 hours. The bacteria were pelleted down by centrifuge at 6,000g for 30 min and then resuspended in 50 mL Buffer 1 (50 mM Tris-HCl, pH=8, 100 mM NaCl, 1mM DTT, 5% Triton-X-100, 1 mM EDTA). The bacteria were then sonicated with 2-second sonication and 2-second rest for 2 min and rested for another 2 min on ice. The sonication was repeated for 4 times. The lysed bacteria were then pelleted down at 6,000 g for 15 min, washed in 50 mL Buffer 1, and subjected to the same sonication procedure two more times. The pellet was then washed in 50 mL Buffer 2 (50 mM Tris-HCl, pH=8, 100 mM NaCl, 1 mM EDTA) and subjected to the same sonication procedure 2 more times. Finally, the pellet was solubilized in 25 mL Buffer 3 (8 M urea, 50 mM Tris-HCl pH=8, 10 mM EDTA, 10 mM DTT) by rotating overnight. The inclusion body was run on SDS-PAGE to check the protein molecular weight.

**Refolding of peptide-MHC**

We prepared 1 L refolding buffer containing 5 M urea, 400 mM L-Arginine, 100 mM Tris-HCl (pH=8), 2 mM EDTA, 0.5 mM oxidized glutathione, and 5 mM reduced glutathione. We then dissolved 30 mg UV-cleavable peptide in 1 mL DMSO and added it into the refolding buffer. Next, we mixed 30 mg B35 MHC heavy chain inclusion body and 30 mg human β-2-microglobulin inclusion body and added this into refolding buffer through a 23G needle by gravity. The refolding buffer was then transferred to dialysis tubing and dialyzed against 10 L 10 mM Tris-HCl (pH=8). Every 12 hours, the 10 mM Tris-HCl (pH=8) buffer was changed to fresh buffer; the dialysis was performed for a total of 60 hours. After dialysis, the refolding buffer was purified by flowing through diethylaminoethyl (DEAE-) cellulose which was equilibrated using 10 mM Tris-HCl (pH=8). The refolded pMHC was eluted from the DEAE-cellulose using 0.5 mM NaCl in 10 mM Tris-HCl (pH=8), buffer exchanged to 10 mM Tris-HCl (pH=8), and concentrated by Amicon concentrator. The protein was then biotinylated overnight at 4 °C. The biotinylated pMHC protein was then purified via size-exclusion chromatography (GE Superdex 200 increase 10/300 GL) and fractions were analyzed by running on SDS-PAGE. Fractions with refolded pMHC protein were further purified by running on a Mono-Q column. Fractions were again analyzed by SDS-PAGE and fractions with refolded pMHC protein were then flash frozen in liquid nitrogen and stored in -80 °C.

**Peptide production**

21 peptide ligands (length = 9 aa) were produced by Fmoc-based solid-phase peptide synthesis (GenScript). The UV-labile peptide used here was KPIVVLJGY, where "J" is Fmoc-(S)-3-Amino-3-(2-Nitro-Phenly)-Propionic Acid (also produced by GenScript). All peptides were dissolved in dimethyl sulfoxide (20 mM) and stored at -20 °C until further use.

**Peptide exchange reaction**

UV-facilitated peptide exchange was performed as described previously (Rodenko et al., 2006). Briefly, we first prepared peptide exchange buffer containing 20 mM Tris-HCl (pH 7.0) and 150 mM NaCl. Next, for each exchange reaction, we combined 40.4 μL exchange buffer, 5.6 μL of the peptide to exchange (500 μM, diluted from 20 mM stocks in DMSO in exchange buffer) and 10 μL UV-labile pMHC (2.8 μM in PBST) in one well of a 96-well plate. The plate was placed on a thermo-mixer whose temperature was pre-equilibrized to 4°C. A UVP compact and handheld UV Lamp (365 nm, UVP95000505, 6W) was placed over the sample for 2 h, with the distance between the lamp and sample set to ~3 cm. After that, the sample was withdrawn from the wells and purified and concentrated using 3K spin columns (ThermoFisher) three times at 4°C. The final concentration was estimated using a NanoDrop 2000 spectrophotometer (Thermo Scientific).

**pMHC-functionalized ‘smart’ bead**

The streptavidin (SA)-coupled ‘smart’ beads were washed with PBST buffer three times. To create sample beads with pMHC at physiological density (for normal BATTLES and force multiplex experiments), we mixed 0.5 µL of 10 nM pMHCs with ~20,000 streptavidin ‘smart’ beads in 50 µL PBST buffer. The mixture was gently rotated for 1 h at room temperature, washed with 200 µL PBST four times, and then resuspended in 200 µL PBST buffer. Bead surfaces were blocked by adding 5 µL of 20 mg/mL biotin-BSA (Thermo Scientific) to the pMHC coated bead slurry for 1 h at room temperature, washed three times in PBST, and then resuspended in 50 µL PBST for further use. The BSA-coated surface eliminates potential non-specific binding of the bead to the T cell(Feng et al., 2017). For the concentration multiplex experiment, we mixed 0.5 µL of 100 nM (7X) or 500 nM (27 X) pMHCs with ~20,000 streptavidin ‘smart’ beads in 50 µL PBST buffer and followed identical bead-coating procedures.

**pMHC quantification by single molecule TIRF microscopy**

To determine the actual density of pMHC molecules on ‘smart bead’ surfaces, we imaged beads via single-molecule total internal reflection fluorescence (smTIRF) microscopy (evanescent wave penetration depth of ~400 nm and a ~50 µm× 50 µm field of view). To detect bead-bound pMHCs, we added 1 µL of PE anti-human β2-microglobulin antibody (BioLegend, clone: A17082A, 0.2 mg/ml) to 5 µL beads solution containing either ~2,000 pMHC- or 2,000 SA-coated beads (for control experiment) and gently rotated for 1 h. After staining, we washed beads with PBST five times, resuspended in 50 µL 1X PBS buffer, and placed samples on ice prior to imaging. Stained beads were imaged by smTIRF microscopy using a simple flow cell created using Scotch double-sided tape, a cover glass (VMR, 22 × 50 mm, #1.5), and a glass slide (Corning, 75 × 25 mm). After loading ~200 beads into the flow cell, the channel was washed with 1X PBS five times and then sealed with nail polish to prevent evaporation during the imaging. For one bead in the field of view, we recorded 10 sequential images at 100 ms intervals. To quantify individual pHMC molecules, we counted the number of bright spots using “Find maxima” in ImageJ with an intensity prominence of 40 and spots with different coordinates were summed across all 10 images.

**Shear modulus measurement**

To access the modulus of rigidity/shear modulus of each code, we made NIPAm slabs containing identical ingredients as the 21 codes and then added 50 µL of each code to the ~15 mm diameter wells in the lid of 24-well plate. After 2-min UV polymerization, we washed the slab three times and rehydrated in colorless RPMI. We then measured the shear modulus (modulus of rigidity) using the TA rheometer (HR 10, Discovery Hybrid Rheometer) with a 20 mm parallel plate associated with Peltier plate Steel plus Solvent Well. The strain and angular frequency were set to 0.015% and 0.1 rad/s to 100 rad/s with 5 points per decade, respectively. The shear modulus is the root sum square of the storage modulus and loss modulus.

**Quantifying bead shrinking and swelling with changes in temperature**

All bead imaging used a small dissection scope (AmScope) equipped with a 20X objective with either a 0.5X relay lens or a 1X relay lens (for the force multiplexing beads) and a Thorlabs camera (DCC3240M). Prior to imaging, we washed SA-coated ‘smart’ beads with PBST three times, resuspended in colorless RPMI buffer, and then loaded them into a flow cell. After bead loading, we sealed the flow cell with nail polish (to prevent evaporation) and placed the flow cell on an indium tin oxide (ITO) glass slide (Bioscience Tools, TC-1-100s). While imaging continuously, we: (1) heated the sample from room temperature (25°C) to 37°C for 1 min (the heating ramp took ~8 sec), (2) cooled the sample to 34°C (the cooling ramp took ~30 sec) and then maintained the sample at 34°C for 2 min. Finally, we quantified bead edges from bright field images by tracing bead edges for every frame using ImageJ and plotting the Feret diameter over time.

***‘*Smart’ bead loading**

Prior to loading ‘smart beads’, we treated microwell devices with air plasma for 4 min at 150 W plasma (Femto, Diener). Using a syringe pump, we then immediately filled the device with PBST 100 µL/h to expel the air trapped inside each well (for ~10 min) and subsequently flowed PBSST at 20 µL/h for 1 h to block the surface. To load beads, we pooled ~10 µL of resuspended solution from each code, washed with PBST one time, and then resuspended in 20 µL PBST. This bead mixture was loaded into the microwell device at 50 µL/h to fill the chamber and then at 10 µL/h for ~ 2h in order to fill most of the wells with beads. We then rinsed the device with PBST at 100 µL/h to flush away extra beads before introducing cells.

**T cell staining**

To monitor TCR signaling, Cal-520 AM (AAT Bioquest, Inc.) was used to stain T cells and observe intracellular calcium flux associated with early T cell activation. Briefly, cells were washed, resuspended at 2 million cells/mL in 600 µL of colorless RPMI 1640 (Life technology), 3% (v/v) FBS, 0.02% (w/v) Pluronic F-127 (Sigma-Aldrich, 3% w/v in PBS buffer) and 2 µM Cal-520 (AAT Bioquest, 2 mM in DMSO), and incubated for 40 min at 37°C. During this incubation, cells were resuspended by gently pipetting the solution every 10 min. T cells were then washed with 1 mL colorless RPMI 1640 at RT and resuspended in 60 µL colorless RPMI 1640 with 3% (v/v) FBS at RT prior to loading into the device.

**Cell loading**

During cell staining, the microwell device was washed with colorless RPMI 1640 at 50 µL/h for 40 min at RT. The stained cells were then loaded into the microwell chip at 30 µL/h for 5 min and then incubated for 10 min to allow stained T cells to settle on top of the beads within the microwells. To ensure even cell loading, we then removed the syringe containing the stained cells from the device input, resuspended cells again via repetitive cycles of withdrawing and injection, and then repeated cell loading from the opposite inlet via an identical procedure. Repeating this injection twice allowed most of the beads to be associated with at least one cell. After loading, we washed away unbound cells using colorless RPMI 1640 supplemented with 3% (v/v) FBS at a flow rate of 20 μL/h for 30 min and then incubated at RT for another 30 min. Based on device microwell dimensions, flow rates <50 μL/h only generated drag forces <4 pN, preventing potential force-induced T cell activation during the loading and washing steps; cell loading was also carried out at room temperature to avoid generating any thermo-responsive forces during cell loading.

**Ca^2+^ flux imaging**

Time-lapse imaging experiments were performed on an automated inverted microscope (Nikon Eclipse Ti, Nikon) with a motorized filter turret. For 10-min Ca^2+^ imaging, exposure times were kept 300 ms during all experiments to prevent pixel intensity saturation. Images were acquired at × 4 magnification (S Plan Fluor ELWD 20x Ph1 ADM; Nikon) with 2×2 binning on a sCMOS Andor camera (Zyla 4.2, Andor Technology plc., Belfast, Northern Ireland) using µManager Software. To heat and cool beads, the entire microwell chip assembly was placed on an ITO glass slide mounted on the ASI stage. To exert force on bead-associated T cells, the chip was heated to and maintained at 37°C for 1 min and then cooled to and kept at 34°C for 2 min. Immediately after cooling, we acquired a total of 150 Ca^2+^ fluorescence images at 4 s intervals.

**‘Smart bead’ imaging**

To identify cells within each well, we first imaged the device via bright-field imaging with 2x2 binning using a 4x objective. To identify embedded spectral codes within each bead, we then illuminated the device from above using 292 nm excitation via a Xenon arc lamp (Lambda LS, Sutter Instruments, Novato, CA) equipped with an automated filter wheel (Lambda 10-2, Sutter Instruments, Novato, CA) containing a 292/27 bandpass excitation filter (Semrock, Rochester, NY) paired with UG11 absorptive glass (Newport, Irvine, CA). Emitted light was passed through an additional UV blocking filter mounted within a custom 3D printed holder mounted over the objective and then collected within Ln-channels using nine emission filters (435/40, 474/10, 536/40, 546/6, 572/15, 620/14, 630/92, 650/13, and 780/20 nm). For each image, we then identified all beads and determined the Ln ratios most likely to have produced the observed spectra associated with each pixel via linear unmixing relative to a series of Ln reference spectra as described previously (Harink et al., 2019). Finally, we created a matrix associating each microwell (indexed by row and column) with the spectral code of the bead within it.

**Image analyses**

Ca^2+^ fluxes for individual cells were analyzed in ImageJ. First, we duplicated the full stack of the fluorescence images. For one replicate, we segmented individual cells by finding all local maxima using “Find maxima” in ImageJ with an intensity prominence of 350. For the other replicate, we generated thresholded images using the triangle method and converted to binary images. We then combined the segmented images with the thresholded images using the “AND” operator under image calculator to identify individual cells. For Ca^2+^ measurements, we: (1) selected a region of interest associated with each cell using the particle analysis tool in ImageJ with a size from 4-pixel units to infinity, and then (2) recorded fluorescence signals and centroids across 150 images. We verified that cells do not move out of ROIs during the course of the experiment for Ca^2+^ analyses by tracking the centroid of each individual cell using the MultiTracker plugin. We then calculated time-lapse fluorescent intensities for selected ROIs and assigned each to a cell number with unique *x* and *y* coordinates. Signals were analyzed in Matlab (MathWorks) by custom written scripts and can be provided upon request. Briefly, we: (1) extracted the spatial coordinates of centers of individual microwells from the bright-field image, (2) converted spatial coordinates to microwell row and column numbers, (3) assigned each cell to a specific microwell by comparing cell and microwell centroids and assuming a 7.5 µm well diameter, and then (4) associated traces for each cell with the embedded code (and thus the peptide sequence) of the bead present within that microwell.

Ca^2+^ traces were normalized for each cell by plotting the fluorescence ratio at each time point divided by the measured intensity at time zero. Integrated Ca^2+^ signals were calculated by subtracting 1 from each timepoint and integrating Ca^2+^ traces over 10 min; all cells with an integrated Ca^2+^ signal > 0 were considered ‘positive’ cells.

**Bootstrap Hypothesis Testing**

To calculate bootstrapped p-values for each peptide tested using BATTLES, we iteratively: (1) pooled integrated Ca^2+^ signals of all peptides, (2) calculated the number of measurements associated with the peptide of interest (n) and for this entire pool (m), (3) sampled n and m observations with replacement from the merged pool, and (4) calculated the difference (t^*^) between the mean of the first n observations and mean of the second m observations. In each case, each observation was the fold-change of a particular integrated Ca^2+^ signal relative to the mean Ca^2+^ signal across all measurements. We repeated this procedure 100,000 times and then estimated the probability that an observed distribution was statistically significantly different from the pooled distribution as follows:

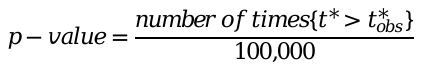

For TCR589 and TCR55 cells interacting with 21 peptides, we applied a Bonferroni correction for multiple hypothesis testing at a significance of 0.05 (0.05/21 = 0.0024). For concentration multiplex experiments, we pooled specific peptides at a given concentration with the same peptide at other concentrations for bootstrap hypothesis testing and therefore determined a Bonferroni-corrected significance threshold of p = 0.013 (0.05/4). For DMF5 T cell interacting with two different peptide classes, we pooled all 5 peptides within each class plus the Tax peptide and therefore determined a Bonferroni-corrected significance threshold of p = 0.008 (0.05/6).

**SUPPLEMENTARY FIGURES AND TABLES**

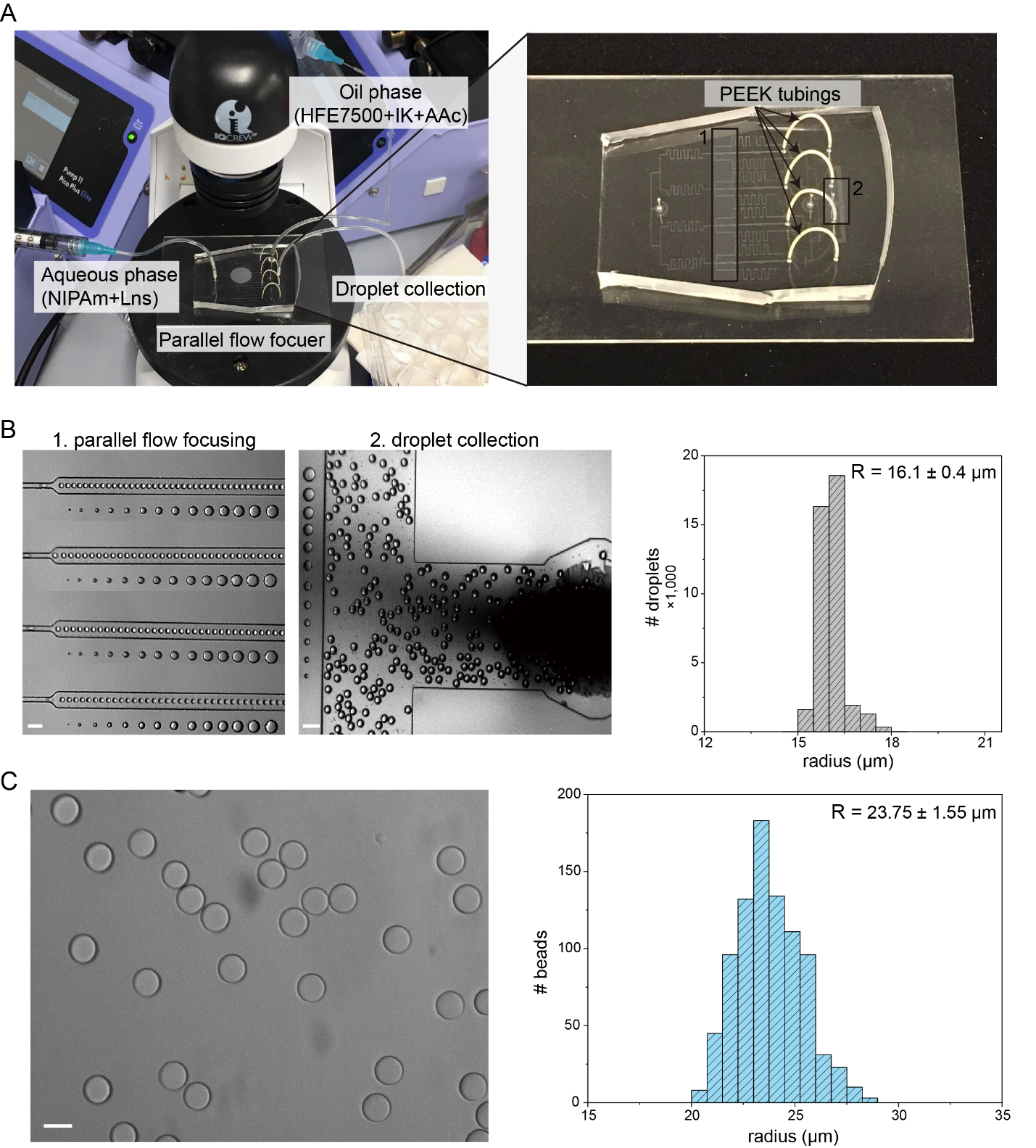

**Figure S1**. Generation of ‘smart beads’ using a microfluidic parallel flow focusing system. (**A**) Bead generation setup that includes a PDMS parallel flow focusing device, a low-cost inverted microscope, and two syringe pumps for injection of aqueous and oil phases. Generated droplets were collected into a 24-well plate. (**B**) Representative images of droplet generation and collection at position 1 and position 2 within the PDMS device (left), with an average droplet radius of 16.1 ± 0.4 μm (right). (**C**) Representative image of ‘smart beads’ after UV polymerization of droplets and rehydration in PBST buffer (left), with an average droplet radius of 23.75 ± 1.55 μm (right). Scale bars: 100 μm in **B** and 50 μm in **C**.

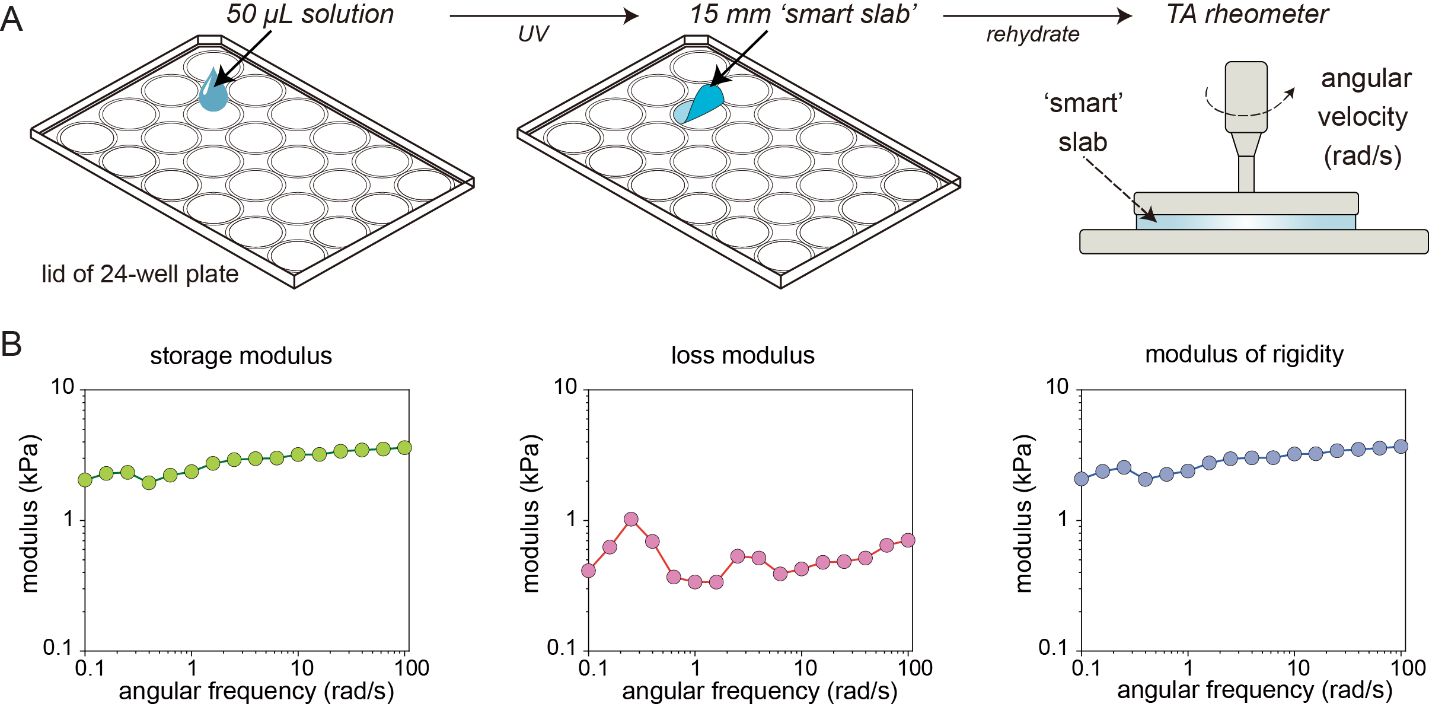

**Figure S2**. Modulus of rigidity *κ* measurement using ‘smart slabs’. (**A**) Procedure to generate ‘smart slabs’ from an identical formulation as ‘smart beads’ followed by the measurement of κ by TA rheometer. (**B**) Measured storage modulus, loss modulus, and modulus of rigidity as a function of angular frequency; *κ* was determined from the root sum square of the storage modulus and loss modulus at each angular frequency and then averaged.

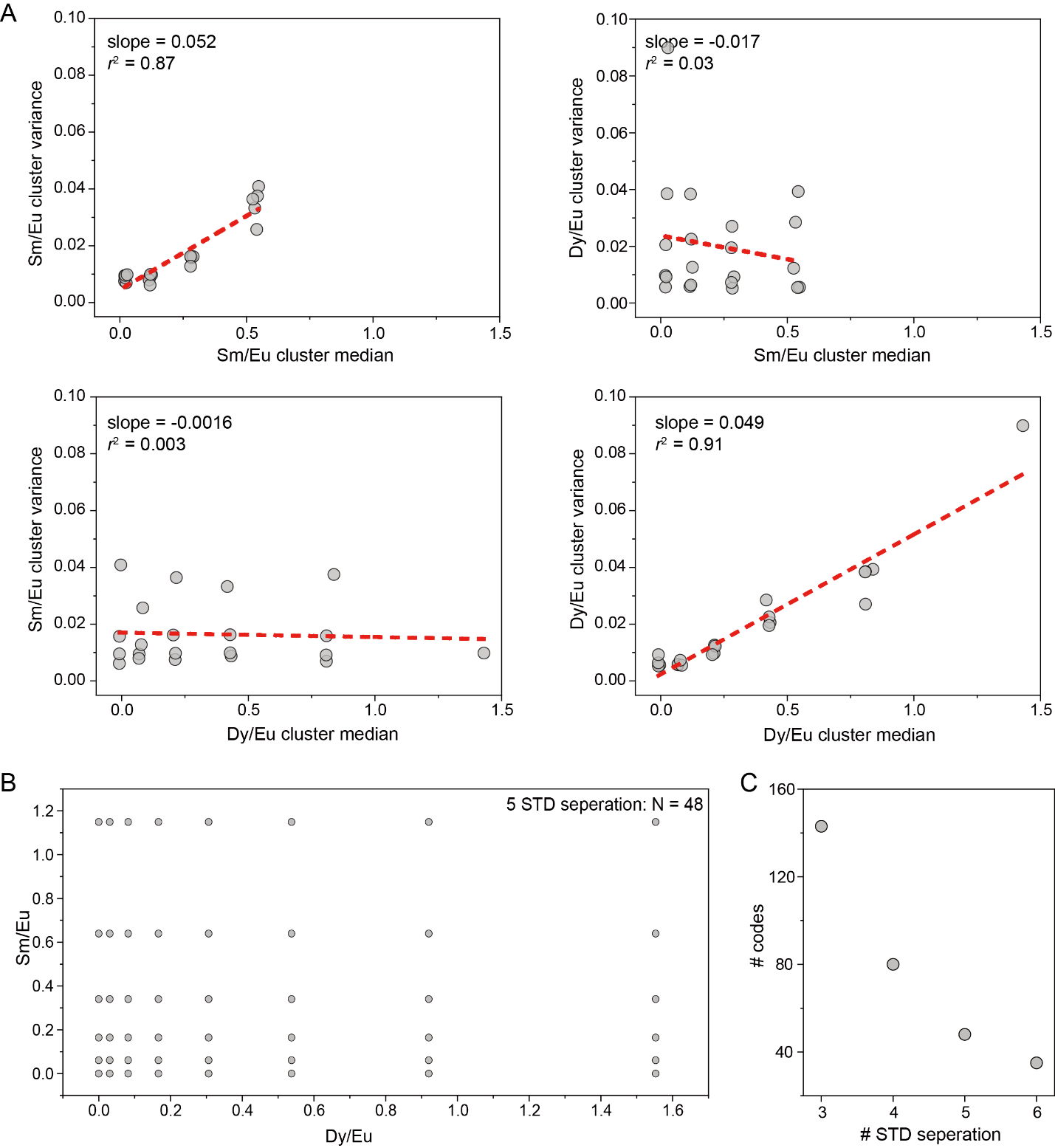

**Figure S3**. Calculated coding capacity for ‘smart beads’. (**A**) Matrix of Sm/Eu and Dy/Eu cluster variances as a function of Sm and Dy cluster medians, respectively. (**B**) Calculated target ratios for a ‘smart bead’ code set with 5-standard deviation spacing between cluster medians. (**C**) Number of calculated code clusters when requiring 3-, 4-, 5- and 6-standard deviation (STD) spacing between each code cluster.

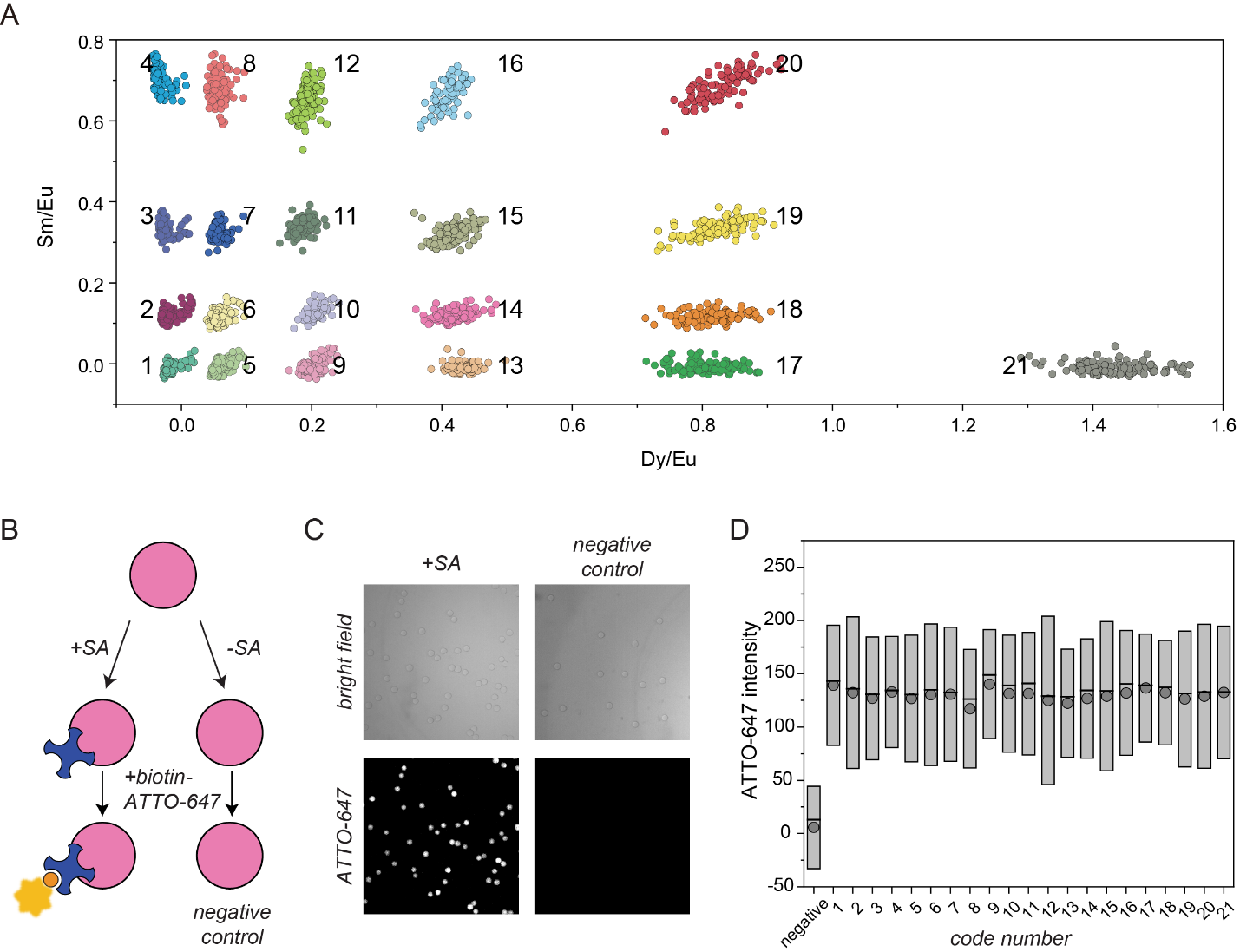

**Figure S4**. Characterization of streptavidin (SA) coated ‘smart beads’. (A) Measured Sm/Eu and Dy/Eu ratios for ‘smart beads’ containing 21 distinct codes after covalent coupling of SA to bead surfaces. (**B**) Experimental pipeline used to quantify coupling efficiency of streptavidin molecules to carboxylated ‘smart beads’. (**C**) Representative bright field (top) and fluorescence (bottom) images of streptavidin-coated (+SA) or negative (-SA) ‘smart beads’ (mixed 21 codes) incubated with biotin-ATTO-647. (**D**) Mean (filled circle) and median (line) of ATTO-647 intensities for the negative control (left) and all encoded ‘smart beads’ from each condition; box represents the standard deviation and the line and grey dot inside indicate the median and mean.

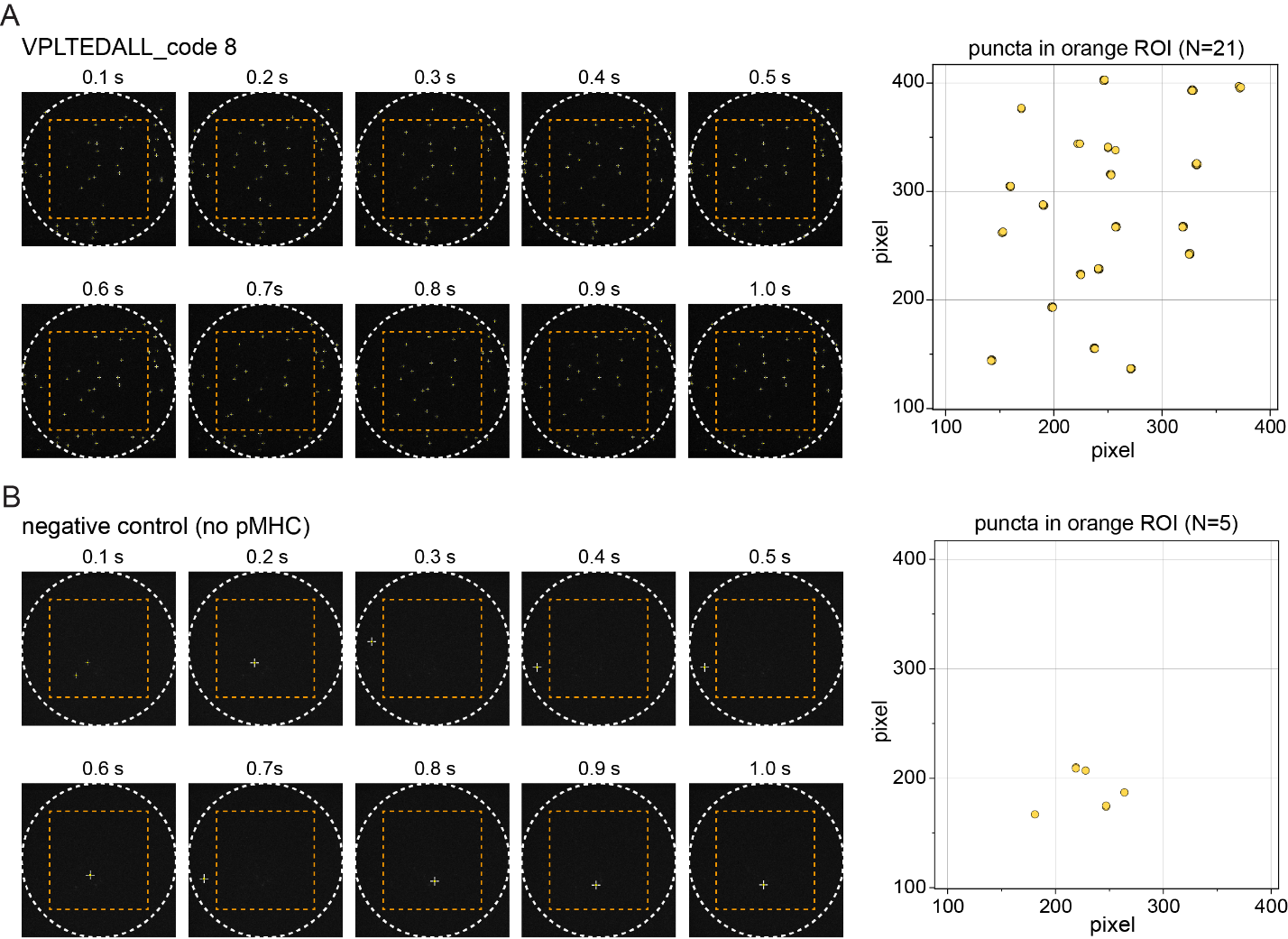

**Figure S5**. Single molecule TIRF fluorescence imaging analysis of pMHC molecules immobilized on ‘smart beads’. (**A**) Representative TIRF images (left) and integrated analysis (counts of bright punctae across all images, right) for a single bead from code 8 bearing biotinylated VPLTEDALL/B35 complexes. In each image, the white circle indicates the radius of the settled bead and the orange square indicates the region of interest used to estimate pMHC density. (**B**) Representative TIRF images (left) and integrated analysis (counts of bright punctae across all images, right) for a single negative control bead. In each experiment, biotinylated pMHC molecules were detected by incubation with PE-labeled anti-β2 mAb (clone: A17082A) immunofluorescence. Detailed bead preparation methods are found in SI Materials and Methods; the orange square ROI is ~840 μm^2^.

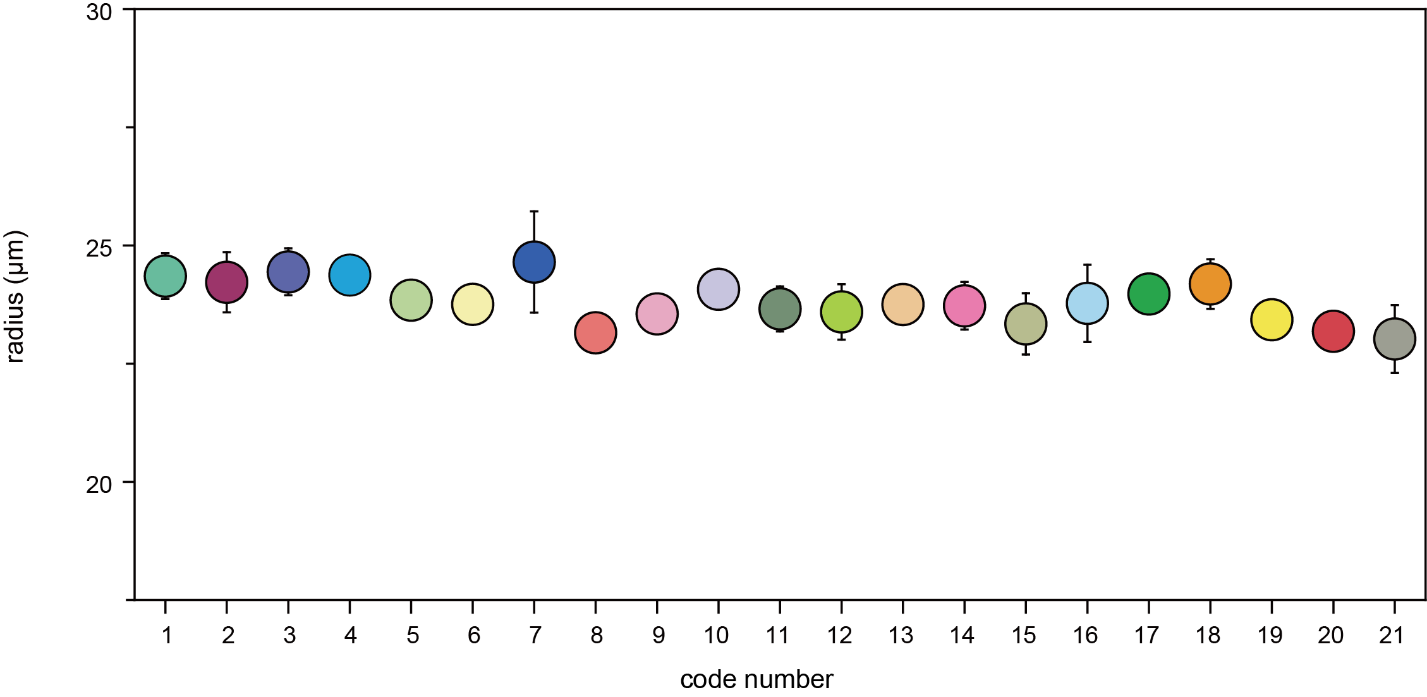

**Figure S6**. Measured radii of ‘smart beads’ across all codes (mean ± standard deviation, n=~40 beads for each code).

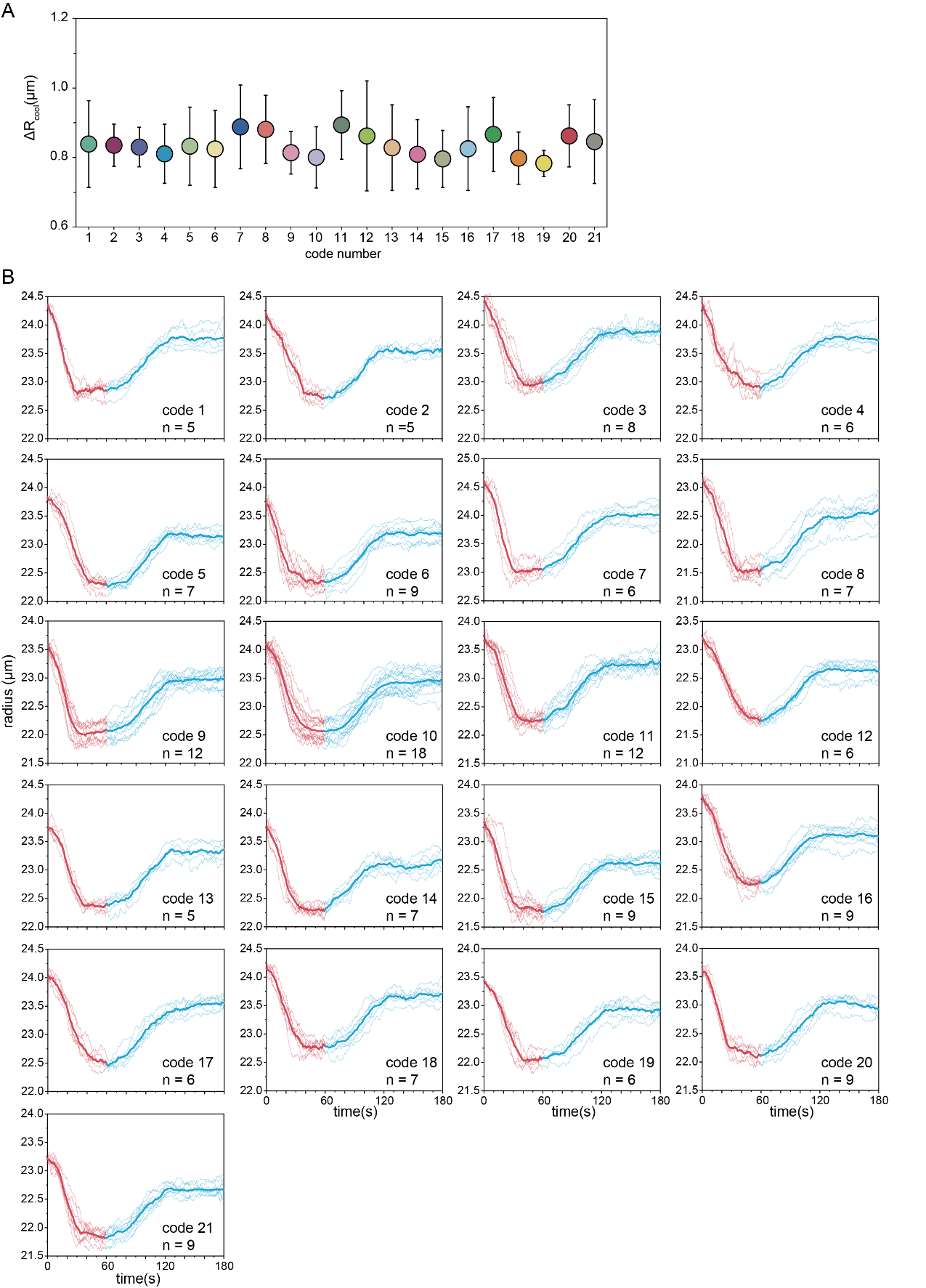

**Figure S7**. Measured changes in bead radii during cooling across all codes. (**A**) Change in bead radii per code (mean ± standard deviation). (**B**) Change in radius as a function of time after changing temperatures to 37°C (at time t=0) and 34°C (at time t=60) for beads from all 21 codes; all radii are normalized to the mean radius at time t=0 for clarity.

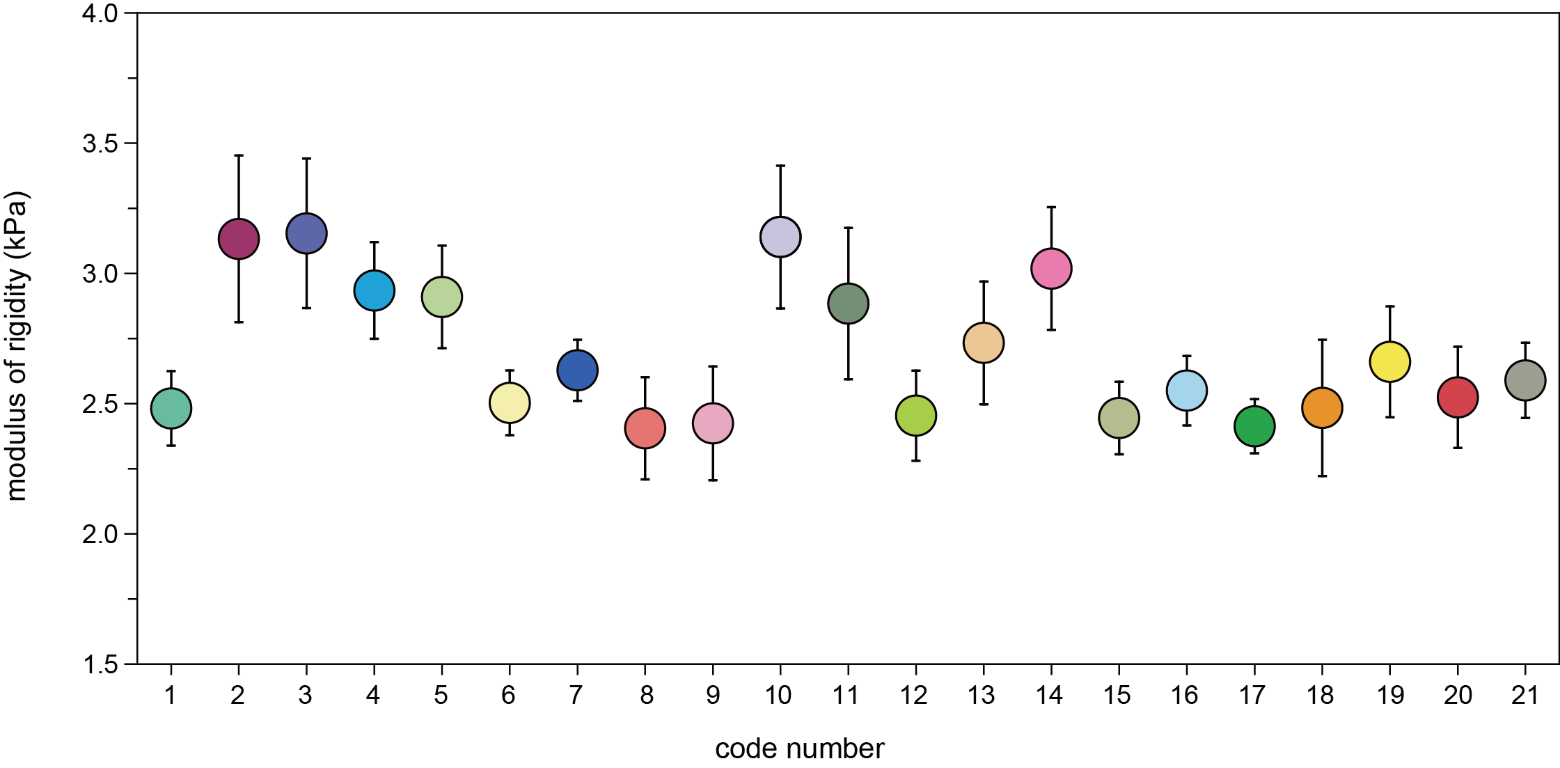

**Figure S8**. Modulus of rigidity (*κ*) for slabs with the same formulation as ‘smart beads’ for each code (mean ± standard deviation).

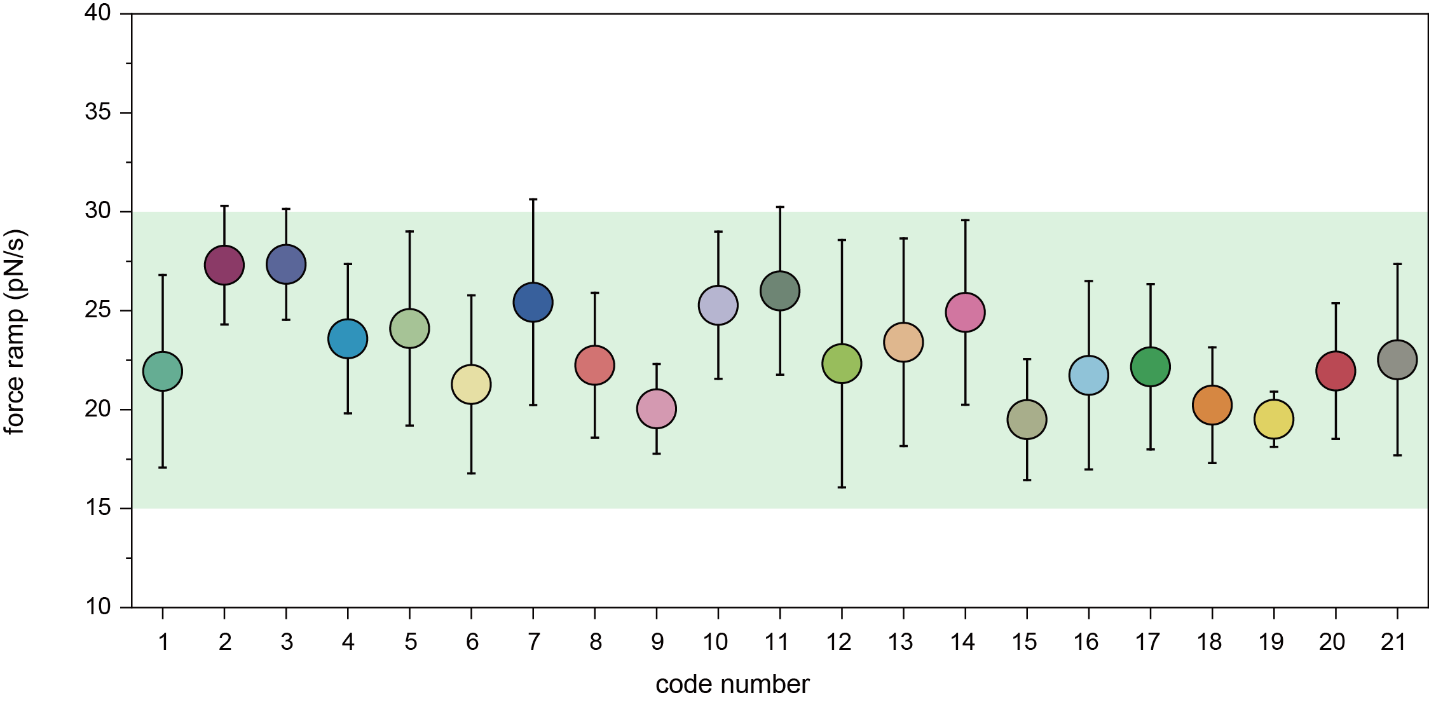

**Figure S9**. Calculated force ramp for ‘smart beads’ for all codes (mean ± standard deviation). The green box represents the physiological cellular force ramps of T cells(Hui et al., 2015).

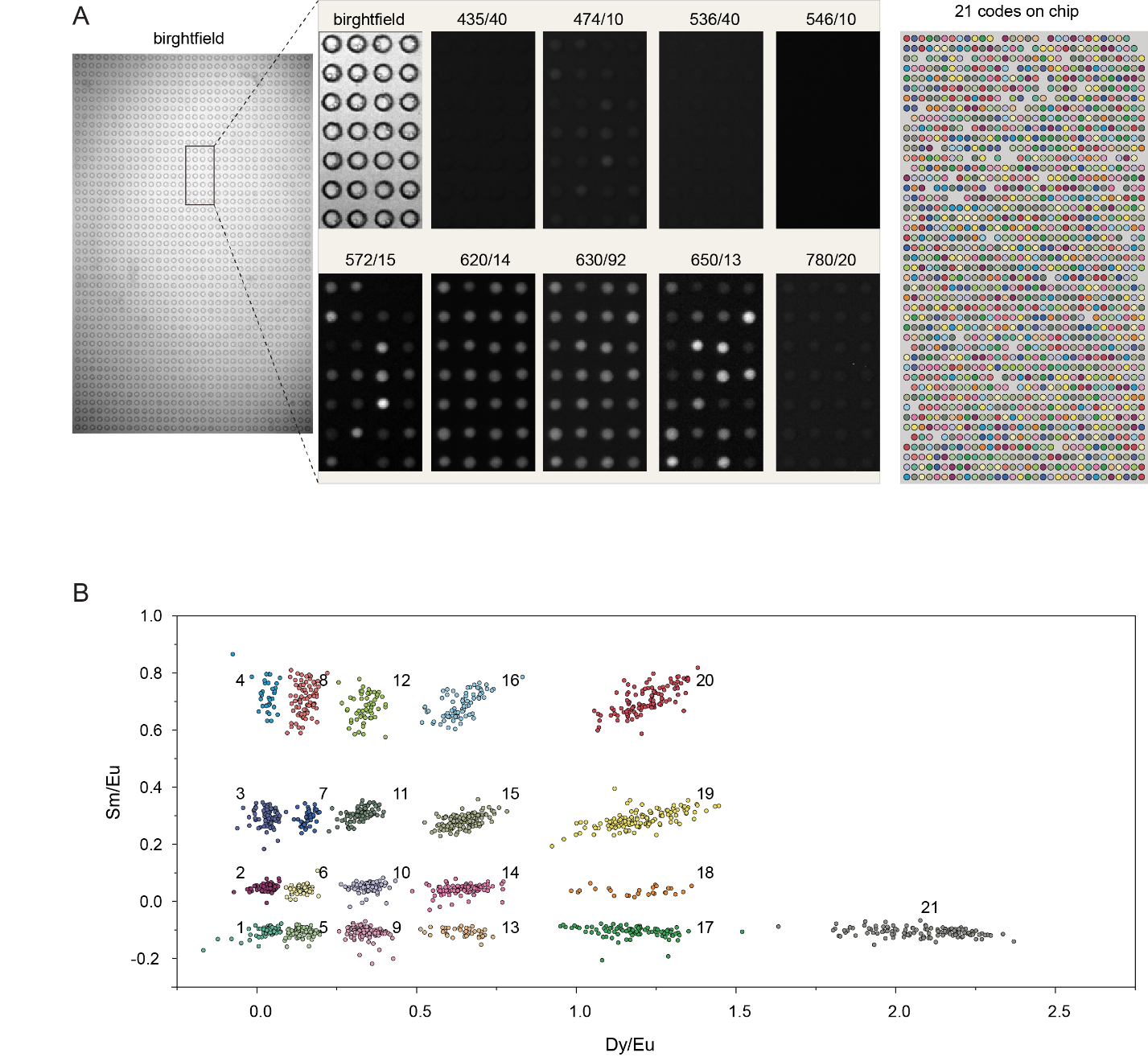

**Figure S10**. Identification of ‘smart bead’ spectral codes after loading beads within the microwell device. (**A**) Left: representative bright field and zoomed in 9 lanthanide emission images of ‘smart beads’ inside microwells. Note that bead intensities are relatively constant in the 620 and 630 nm channels (corresponding to the ‘reference’ Eu Ln added at a constant level to all beads) and vary in the 572 nm and 650 nm channels (corresponding to emission from the Dy and Sm Lns, respectively). Right: False color image illustrating spectral codes for ‘smart beads’ within the microwell device. **(B)** Measured Sm/Eu and Dy/Eu ratios for ‘smart beads’ imaged within microwells with individual code clusters colored and labeled.

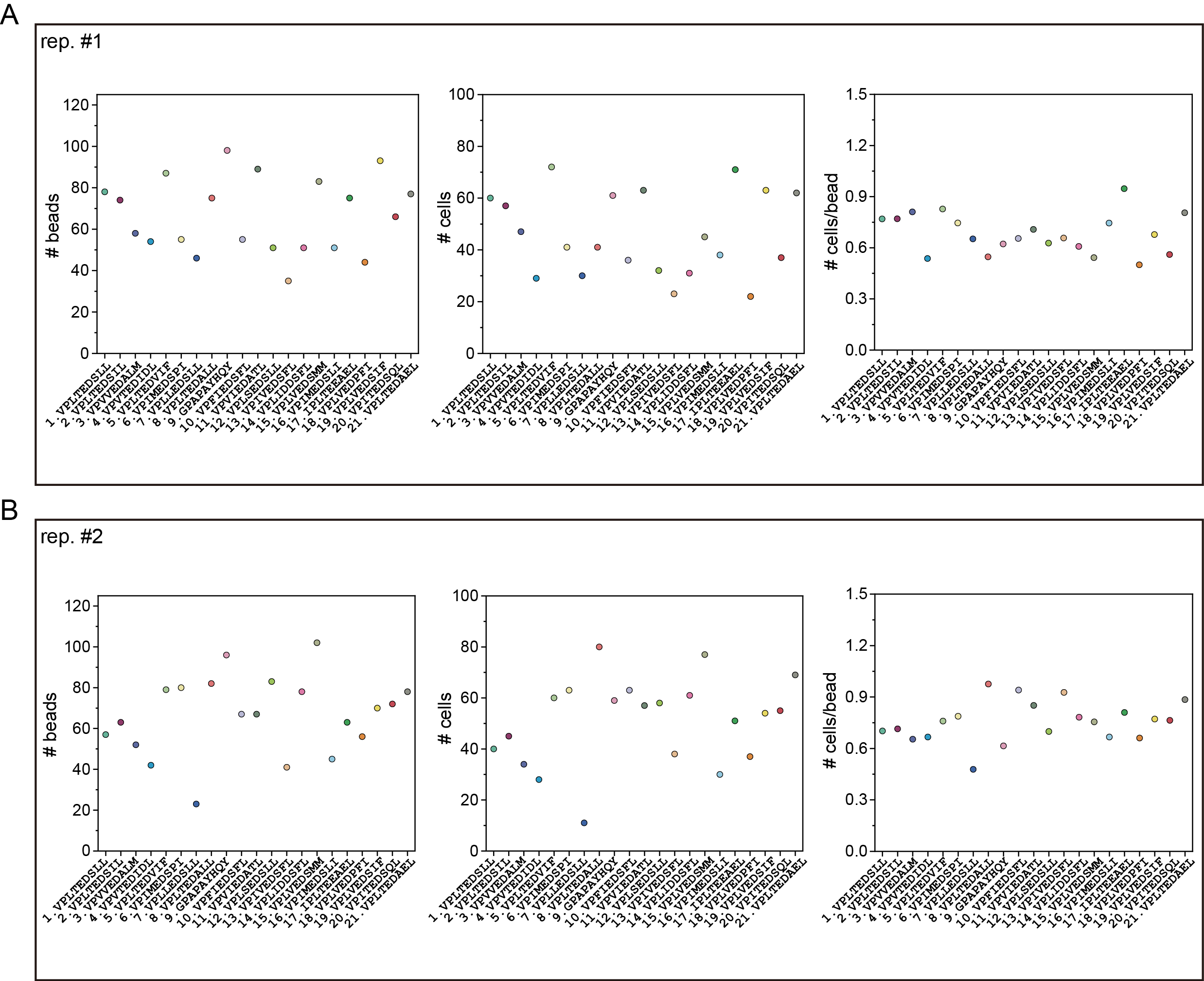

**Figure S11**. No. of loaded beads (left) and cells (middle) per code and cells per bead (right) in two TCR589 activation experiments. (**A**) Replicate 1. (**B**) Replicate 2.

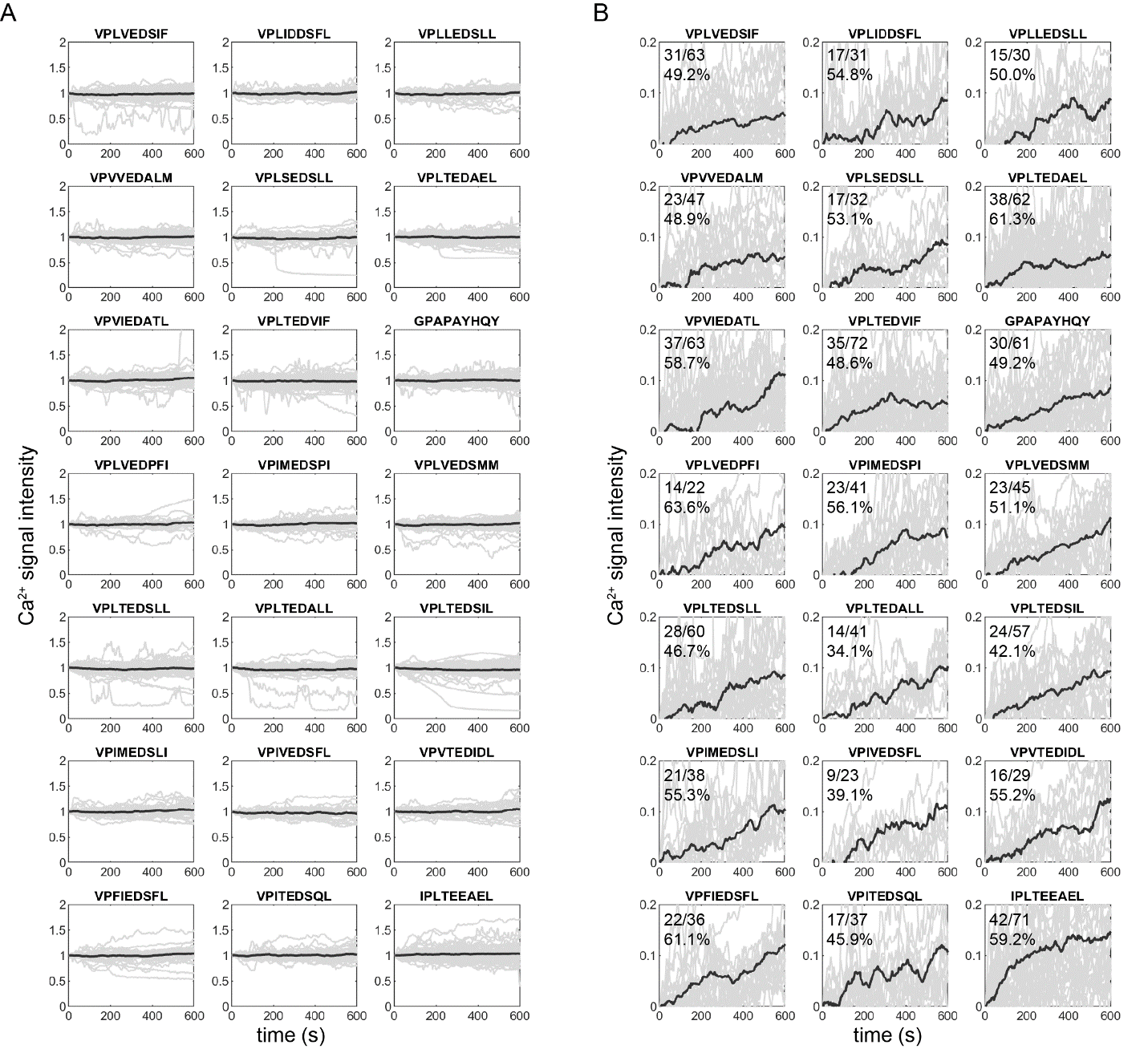

**Figure S12**. Measured Ca^2+^-induced fluorescence as a function of time for TCR589-transduced T cells interacting with 21 different peptide sequences. (**A**) All measured traces. (**B**) All positive traces with the indicated numbers of positive cells and measured cells and positive percentages. Light grey lines indicate traces for individual cells; dark grey lines indicate the mean signals.

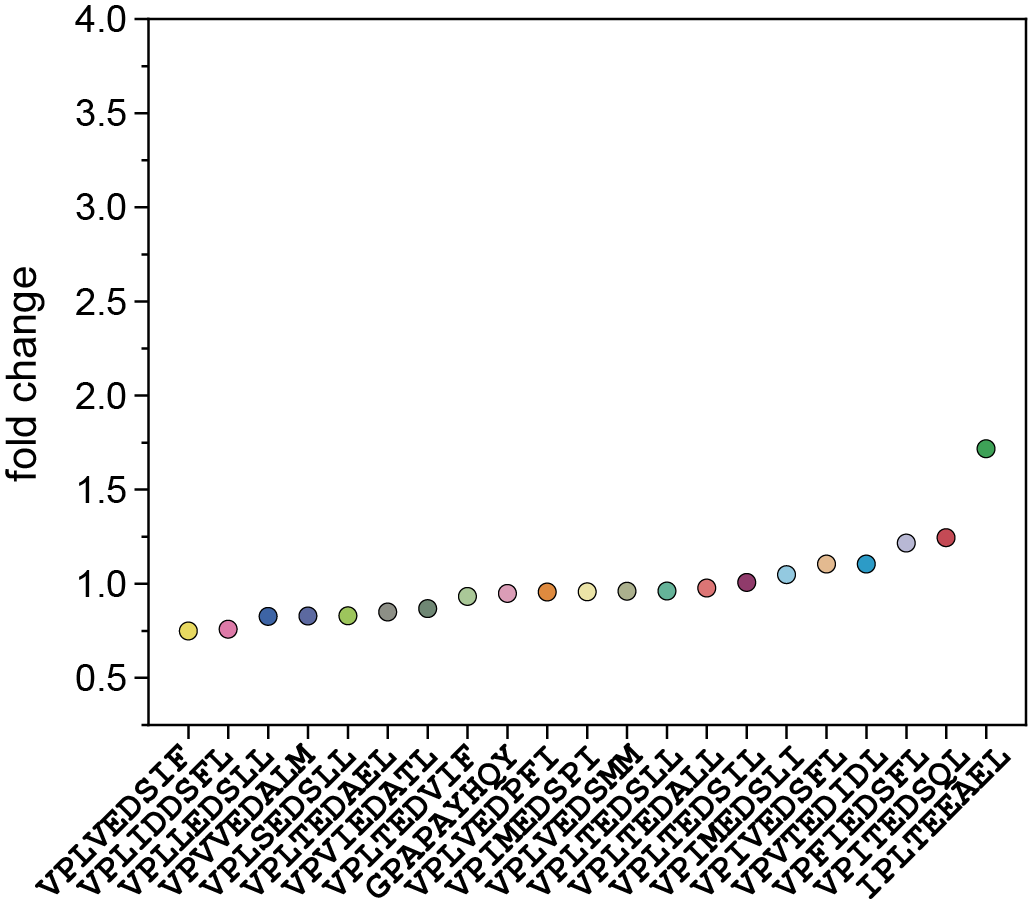

**Figure S13**. Averaged fold changes of integrated Ca^2+^ signals by peptide relative to the mean for all positive TCR589 cells in replicate 1.

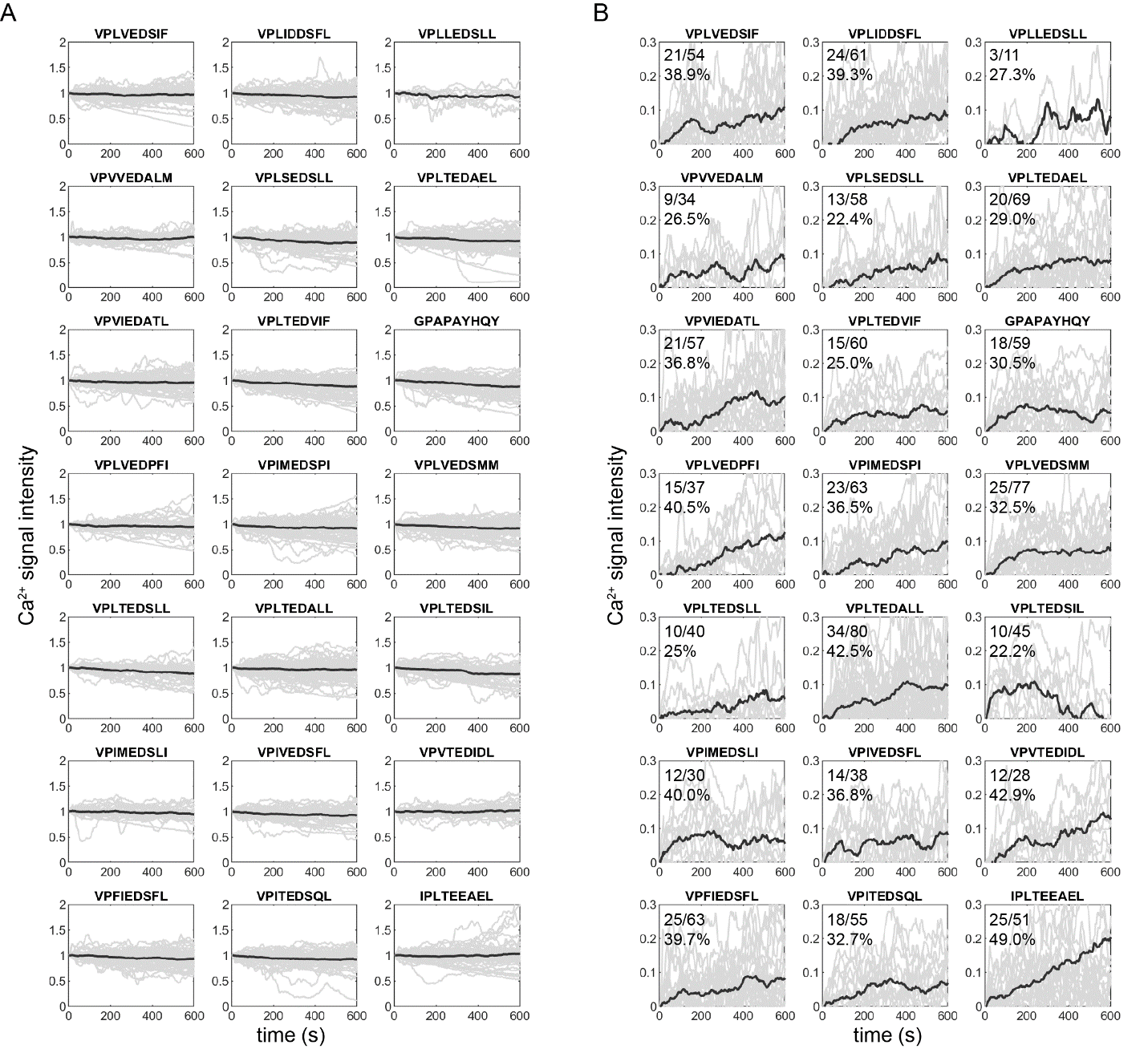

**Figure 14**. Measured Ca^2+^-induced fluorescence as a function of time for another replicate of TCR589-transduced T cells interacting with 21 different peptide sequences. (**A**) All measured traces. (**B**) All positive traces with the indicated numbers of positive cells and measured cells and positive percentages. Light grey lines indicate traces for individual cells; dark grey lines indicate the mean signals.

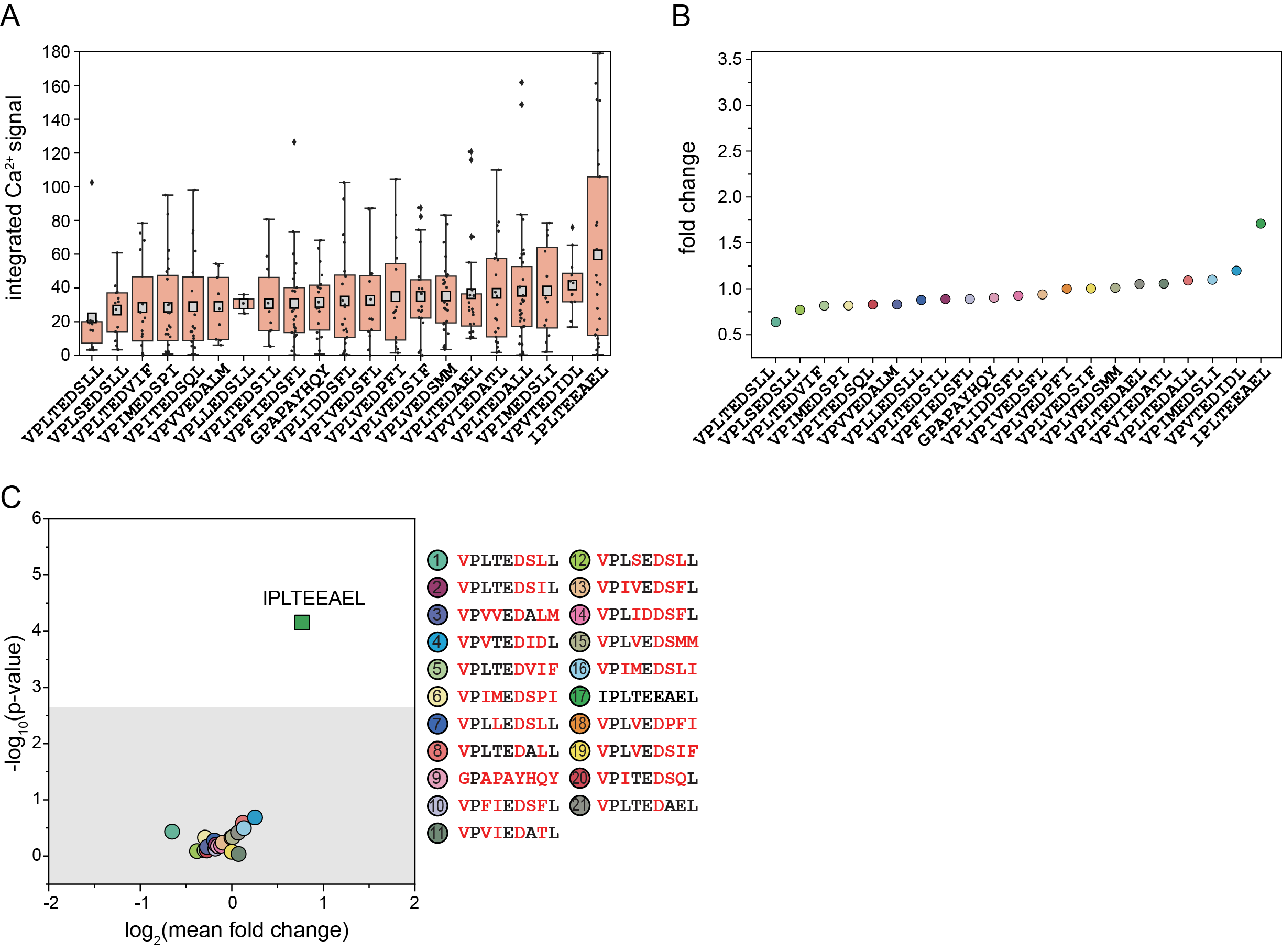

**Figure S15**. Measured integrated Ca^2+^ fluorescence signals and estimated significance for TCR589 replicate 2. (**A**) Integrated Ca^2+^ signals for each positive cell (light grey markers) and the mean integrated Ca^2+^ signal (light grey square). Box lower and upper limits indicate 25^th^ and 75^th^ percentiles, respectively. (**B**) Averaged fold changes of integrated Ca^2+^ signals by peptide relative to the mean for all positive cells. (**C**) Estimated p-value *vs.* log2-transformed mean fold change for TCR589-tranduced T cells interacting with 21 different peptide sequences. Grey box indicates Bonferroni-corrected p-value at a significance of 0.05 (p = 0.0024). Correspondence between ‘smart bead’ spectral code and displayed peptide sequence is shown at right.

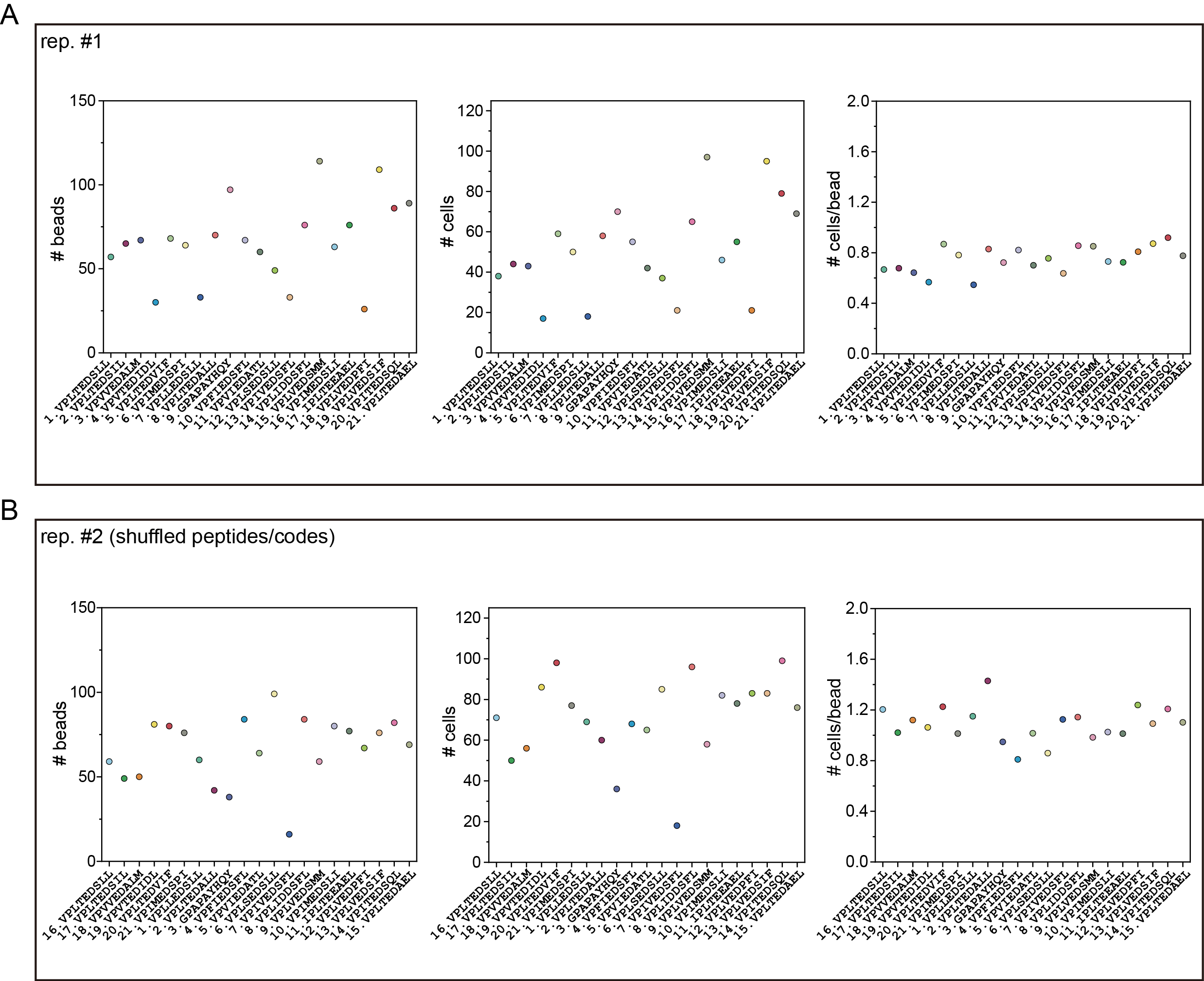

**Figure S16**. No. of loaded beads (left) and cells (middle) per code and cells per bead (right) for two TCR55 activation experiments. (**A**) Replicate 1. (**B**) Replicate 2.

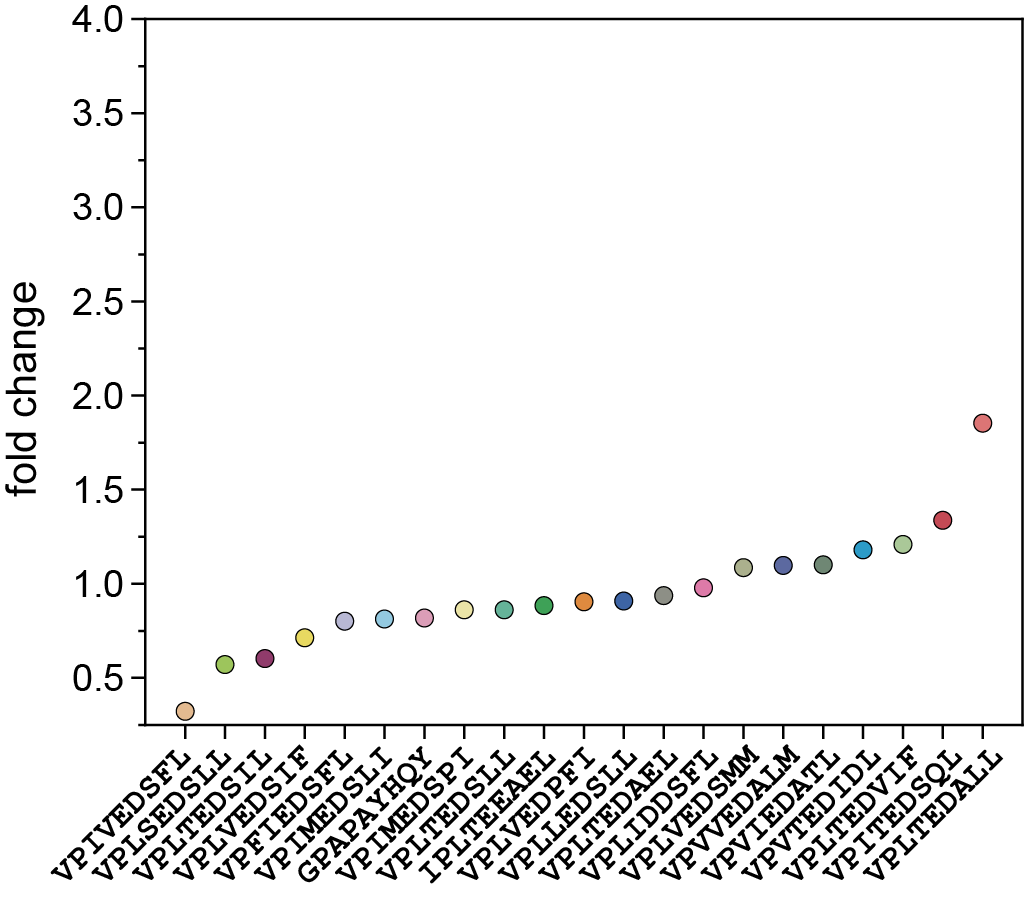

**Figure S17**. Averaged fold changes of integrated Ca^2+^ signals by peptide relative to the mean for all positive TCR55 cells in Replicate 1.

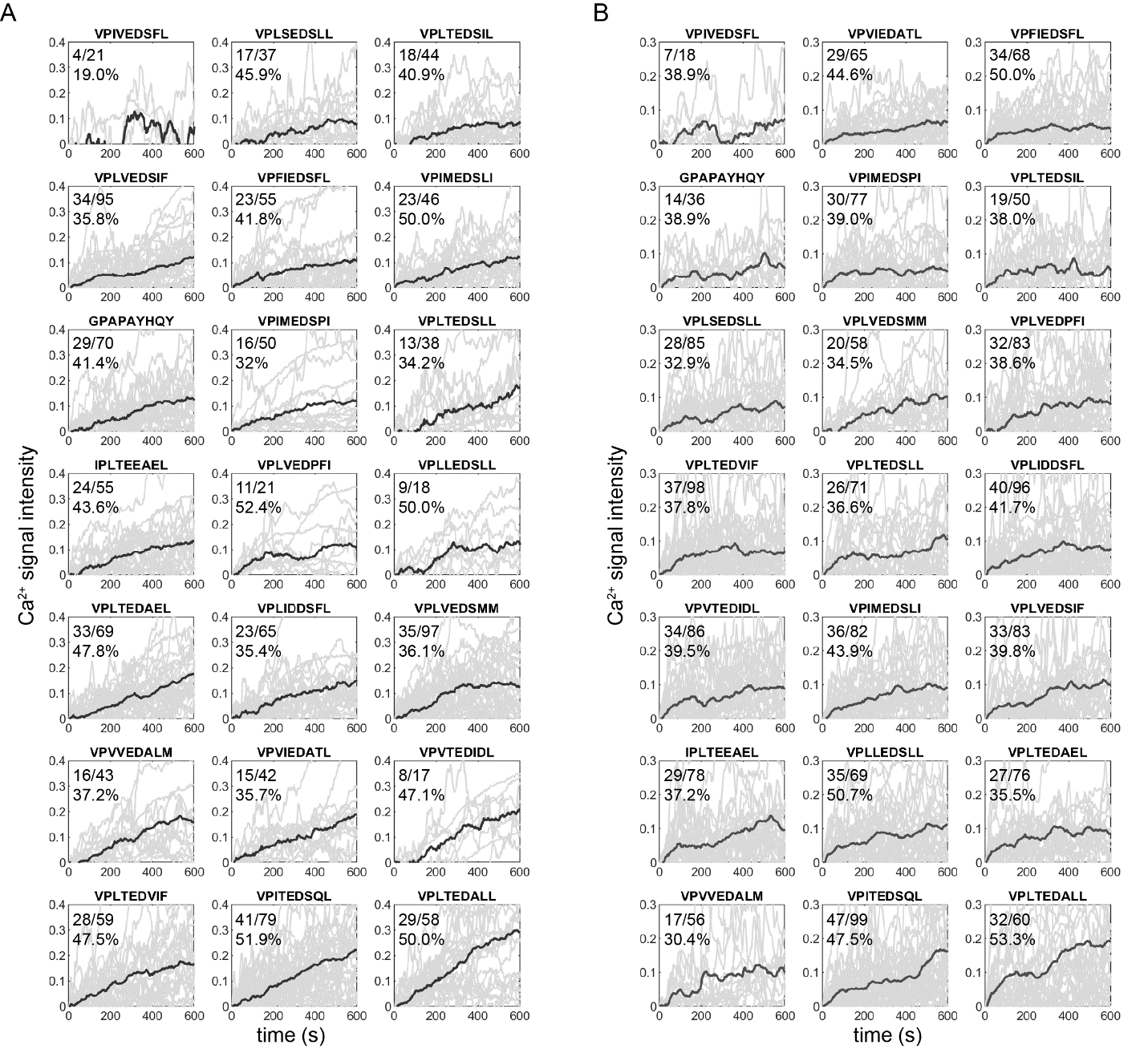

**Figure S18**. Measured positive Ca^2+^-induced fluorescence as a function of time for two replicates of TCR55-transduced T cells interacting with 21 different peptide sequences. (**A**) All positive traces with the indicated numbers of positive cells and measured cells and positive percentages for Replicate 1. (**B**) All positive traces with the indicated numbers of positive cells and measured cells and positive percentages for Replicate 2. Light grey lines indicate traces for individual cells; dark grey lines indicate the mean signals.

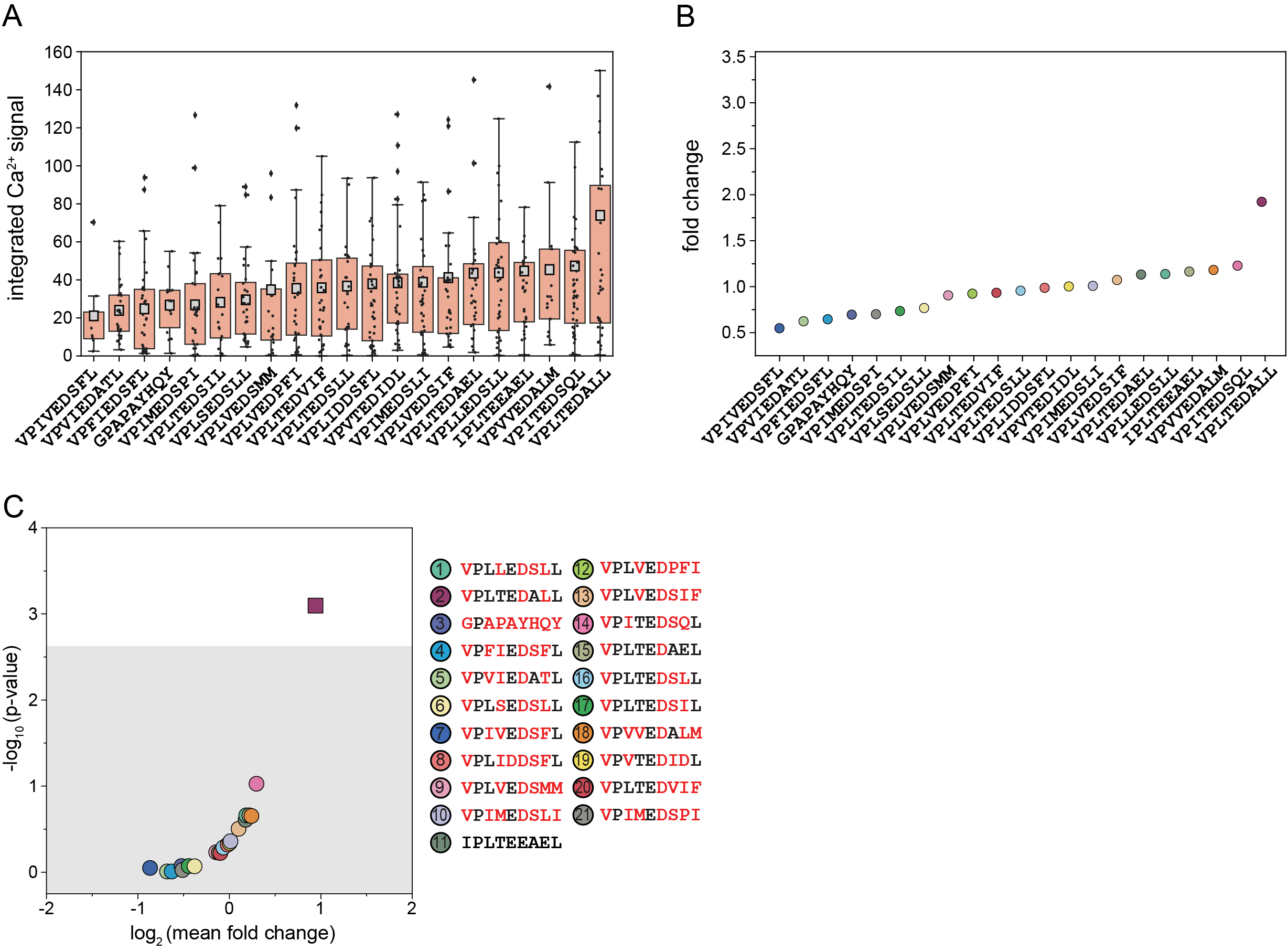

**Figure S19**. Measured integrated Ca^2+^ fluorescence signals and estimated significance for TCR55 replicate 2. (**A**) Integrated Ca^2+^ signals for each positive cell (light grey markers) and the mean integrated Ca^2+^ signal (light grey square). Box lower and upper limits indicate 25^th^ and 75^th^ percentiles, respectively. (**B**) Averaged fold changes of integrated Ca^2+^ signals by peptide relative to the mean for all positive cells. (**C**) Estimated p-value *vs.* log2-transformed mean fold change for TCR55-tranduced T cells interacting with 21 different peptide sequences. Grey box indicates Bonferroni-corrected p-value at a significance of 0.05 (p = 0.0024). Correspondence between ‘smart bead’ spectral code and displayed peptide sequence is shown at right.

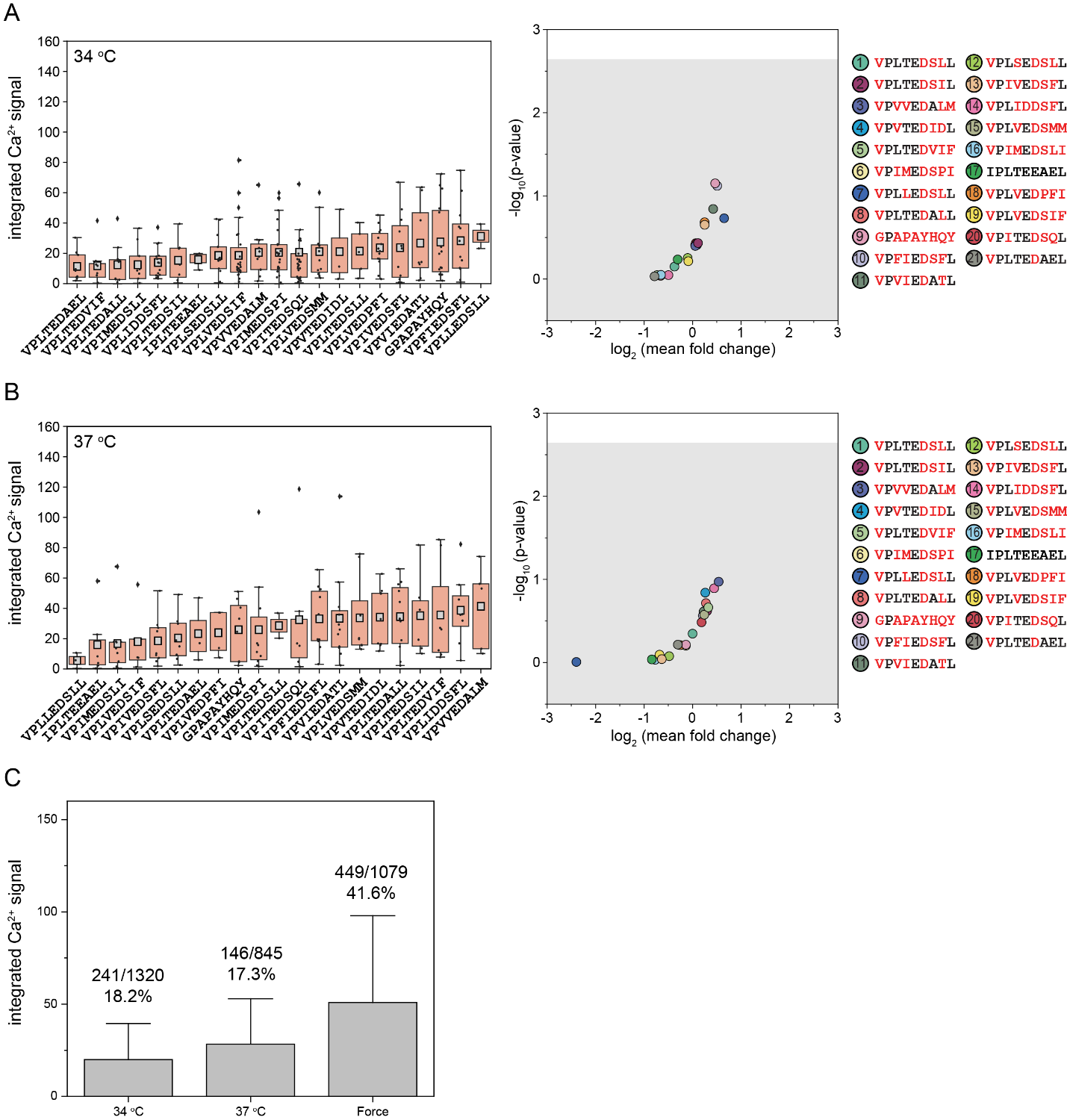

**Figure S20**. Measured integrated Ca^2+^ fluorescence signals and estimated significance for TCR55 negative controls. (**A**) Left: Integrated Ca^2+^ signals for each positive cell (light grey markers) and the mean integrated Ca^2+^ signal (light grey square) at 34 °C. Box lower and upper limits indicate 25^th^ and 75^th^ percentiles, respectively. Right: Estimated p-value *vs.* log2-transformed mean fold change for TCR55-tranduced T cells interacting with 21 different peptide sequences at 34°C. Grey box indicates Bonferroni-corrected p-value at a significance of 0.05 (p = 0.0024). Correspondence between ‘smart bead’ spectral code and displayed peptide sequence is shown at right. (**B**) Left: Integrated Ca^2+^ signals for each positive cell (light grey markers) and the mean integrated Ca^2+^ signal (light grey square) at 37 °C. Box lower and upper limits indicate 25^th^ and 75^th^ percentiles, respectively. Right: Estimated p-value *vs.* log2-transformed mean fold change for TCR55-tranduced T cells interacting with 21 different peptide sequences at 37 °C. Grey box indicates Bonferroni-corrected p-value at a significance of 0.05 (p = 0.0024). Correspondence between ‘smart bead’ spectral code and displayed peptide sequence is shown at right. (**C**) Averaged fold changes of integrated Ca^2+^ signals for all positive cells relative to the mean at 34 °C, 37 °C and with expansion force applied. The positive cell percentages are indicated above each bar.

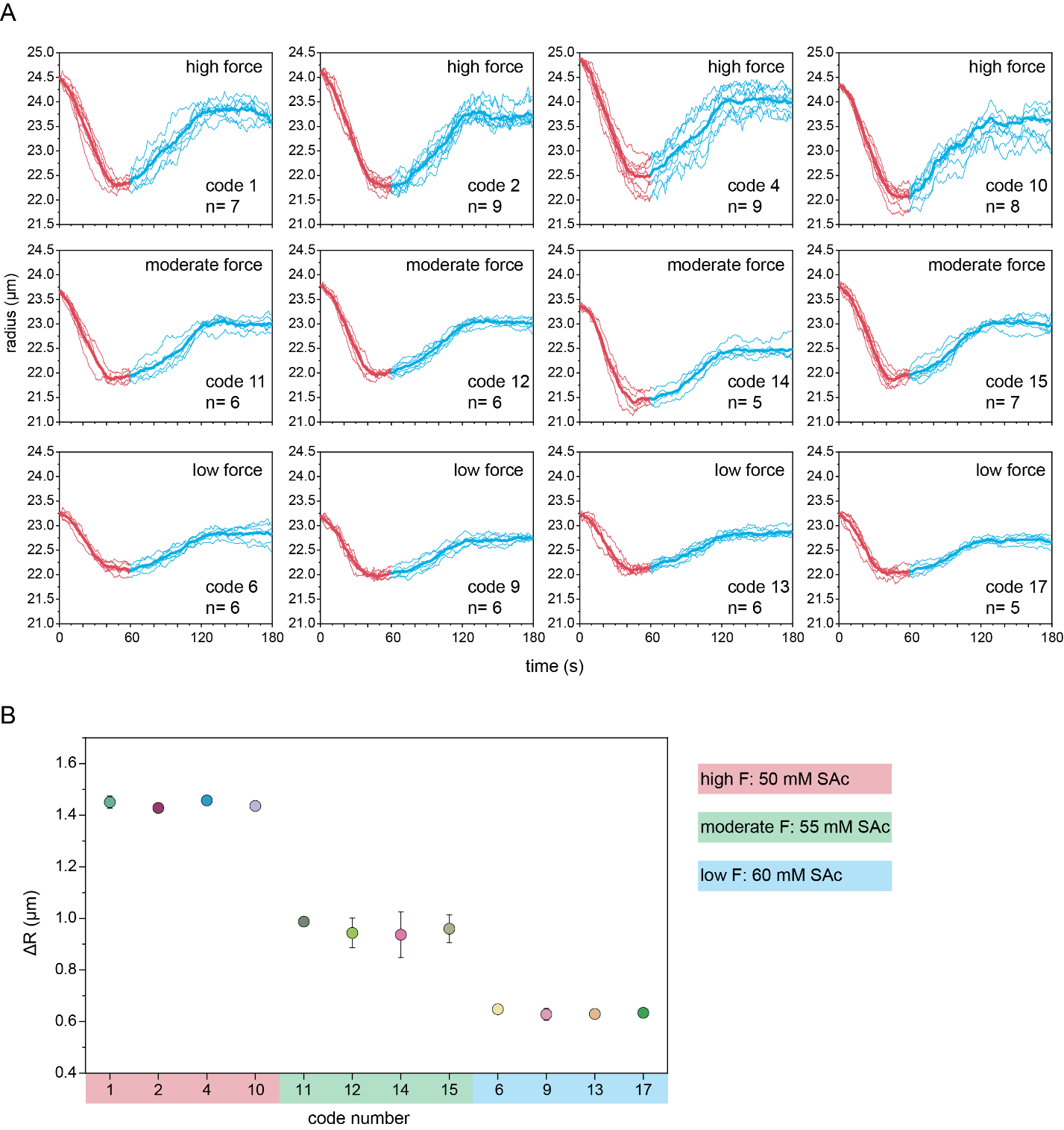

**Figure S21**. Changes in bead radii as a function of temperature for bead codes containing different SAc concentrations. (**A**) Measured radii as a function of time for ‘smart’ beads per code upon changing the temperature to 37 °C at time t = 0 s and to 34 °C at time t = 60 s; all traces are offset such that initial bead radii are set to the mean radius for each code at time t = 0 s. The red portion of each trace indicates bead heating from RT to 37°C; the blue portion of each trace indicates bead cooling from 37 °C to 34 °C. (**B**) Mean (marker) and standard deviation (error bar) for change in radius upon cooling for beads from each code.

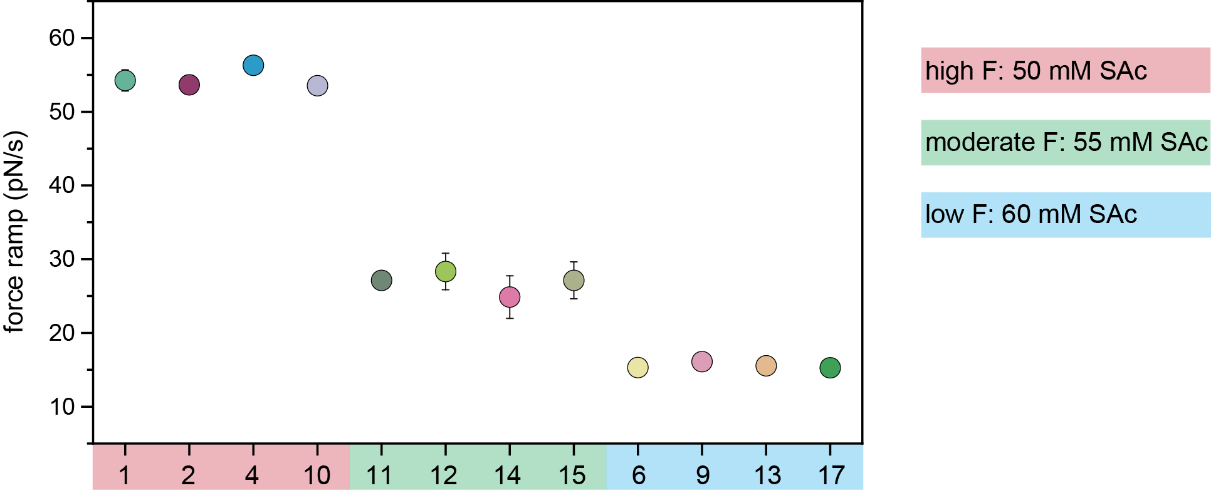

**Figure S22**. Estimated magnitudes of force ramps for 12 bead codes containing different concentrations of SA within the PNIPAm matrix.

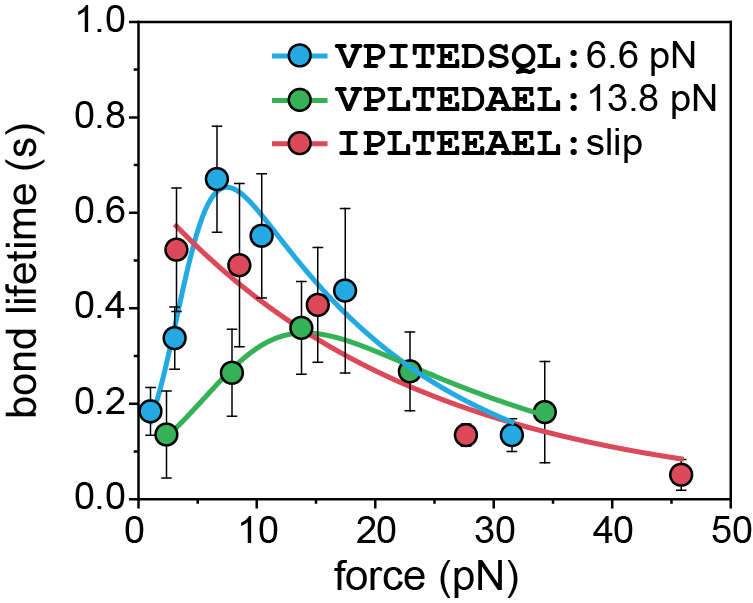

**Figure S23**. Typical catch bonds in the TCR55 T cell system. Data was recapitulated from ref. 2 and fit to the Bell’s model(Bell, 1978) with catch-slip states (for VPITEDSQL and VPLTEDAEL) or slip-only state (for IPLTEEAEL).

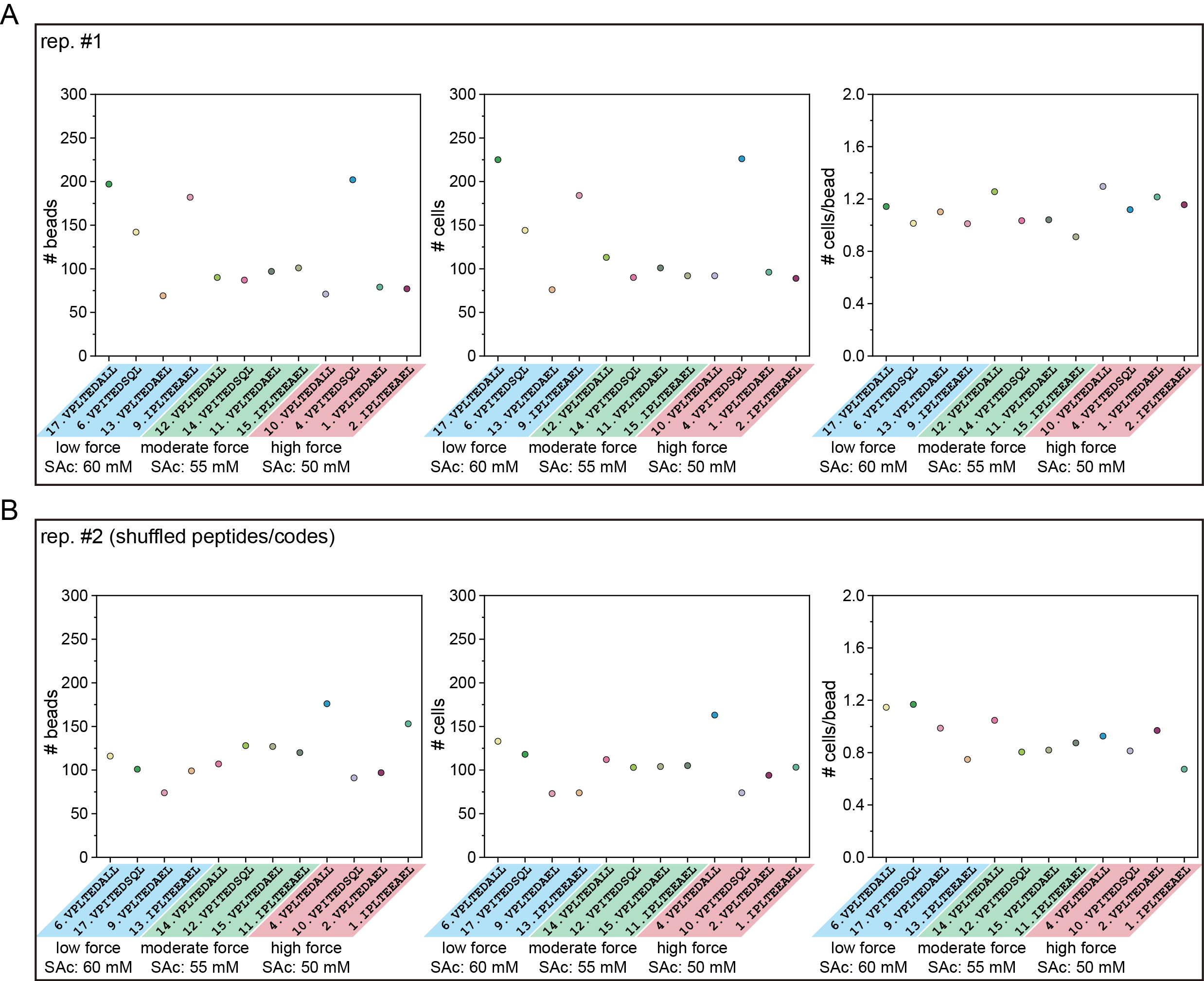

**Figure S24**. No. of loaded beads (left) and cells (middle) per code and cells per bead (right) for two TCR55 force multiplex activation experiments. (**A**) Replicate 1. (**B**) Replicate 2.

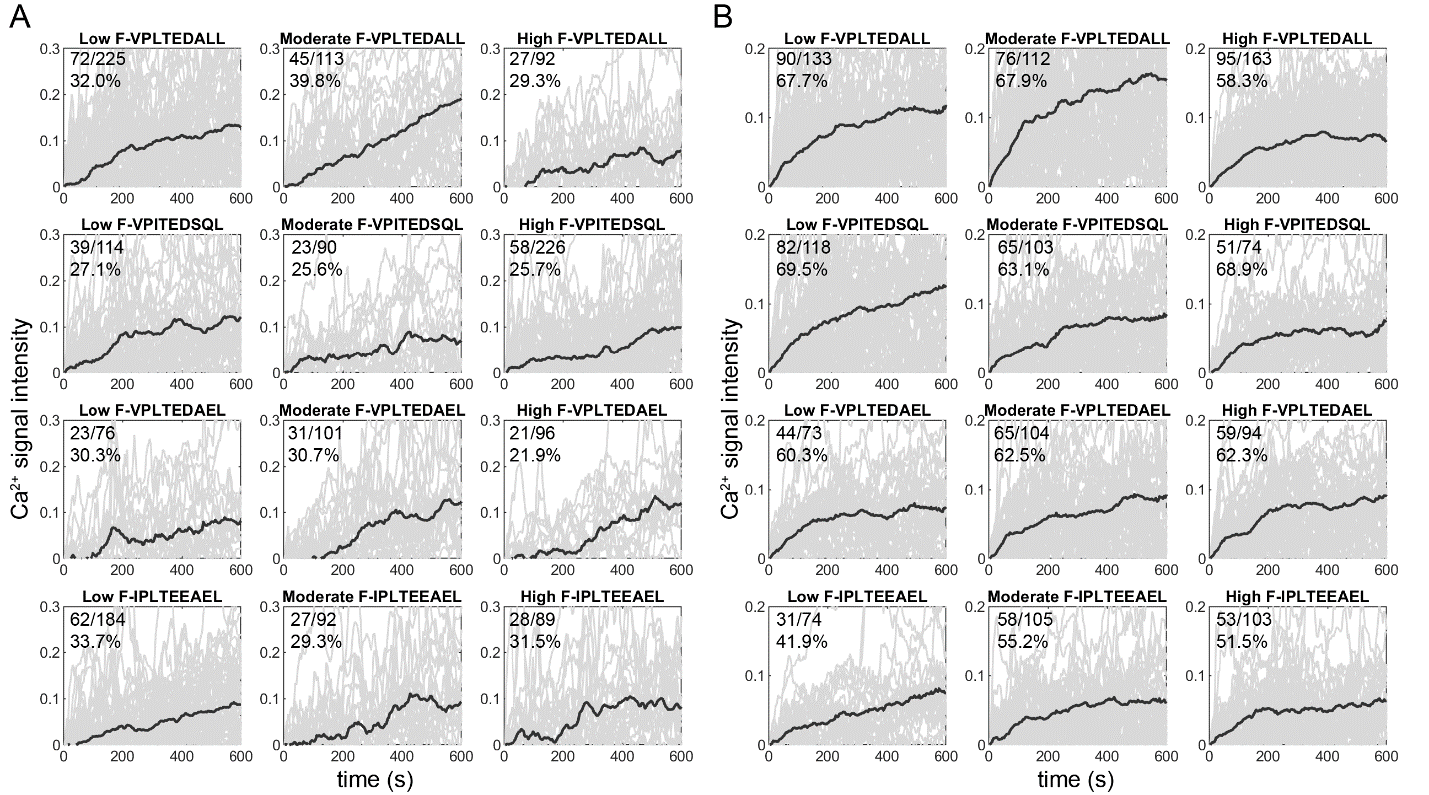

**Figure S25**. Measured positive Ca^2+^-induced fluorescence as a function of time for two replicates of TCR55 force multiplex experiments (3 different force ramps: low (15 pN/s), moderate (25 pN/s) and high (55 pN/s)) interacting with 4 different peptide sequences. (**A**) All positive traces with the indicated numbers of positive cells and measured cells and positive percentages for Replicate 1. (**B**) All positive traces with the indicated numbers of positive cells and measured cells and positive percentages for Replicate 2. Light grey lines indicate traces for individual cells; dark grey lines indicate the mean signals.

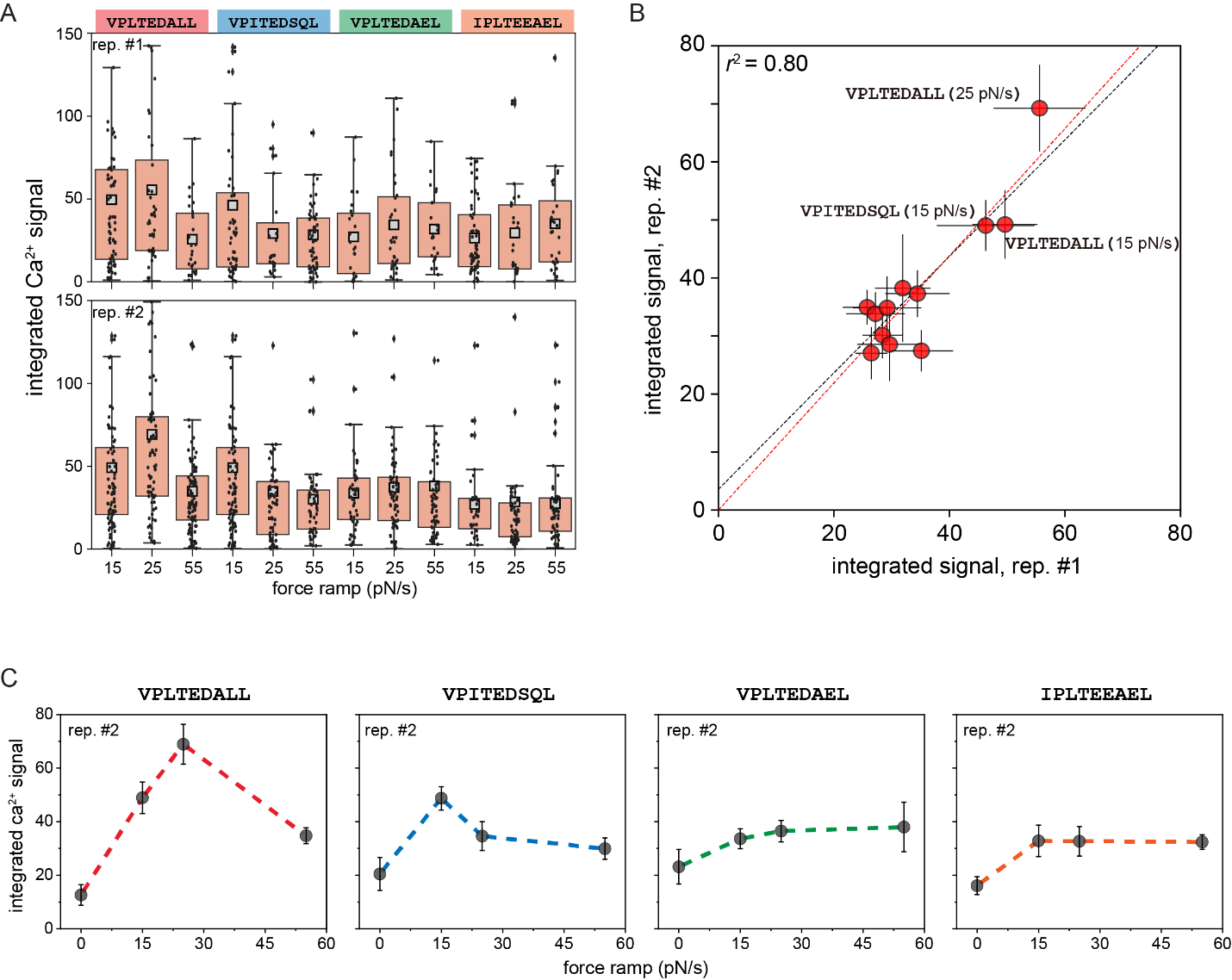

**Figure S26**. Force multiplex for two TCR55 replicates. (**A**) Measured integrated Ca^2+^ fluorescence signals for each positive cell (light grey markers) and the mean integrated Ca^2+^ signal (light grey square) for two TCR55 replicates interacting with 4 different peptides at three force ramps for Replicate 1 (top) and Replicate 2 (bottom). Box lower and upper limits indicate 25^th^ and 75^th^ percentiles, respectively. Zero force data is from a TCR55 control experiment where the temperature was maintained at 34°C; error bars indicate SEM. (**B**) Correlation plot of the means of integrated Ca^2+^ signals between two replicates. Dashed black line indicates the 1:1 line; red dashed line indicates a linear regression. (**C**) Integrated Ca^2+^ _­_signal as a function of applied force for 4 peptides for Replicate 2.

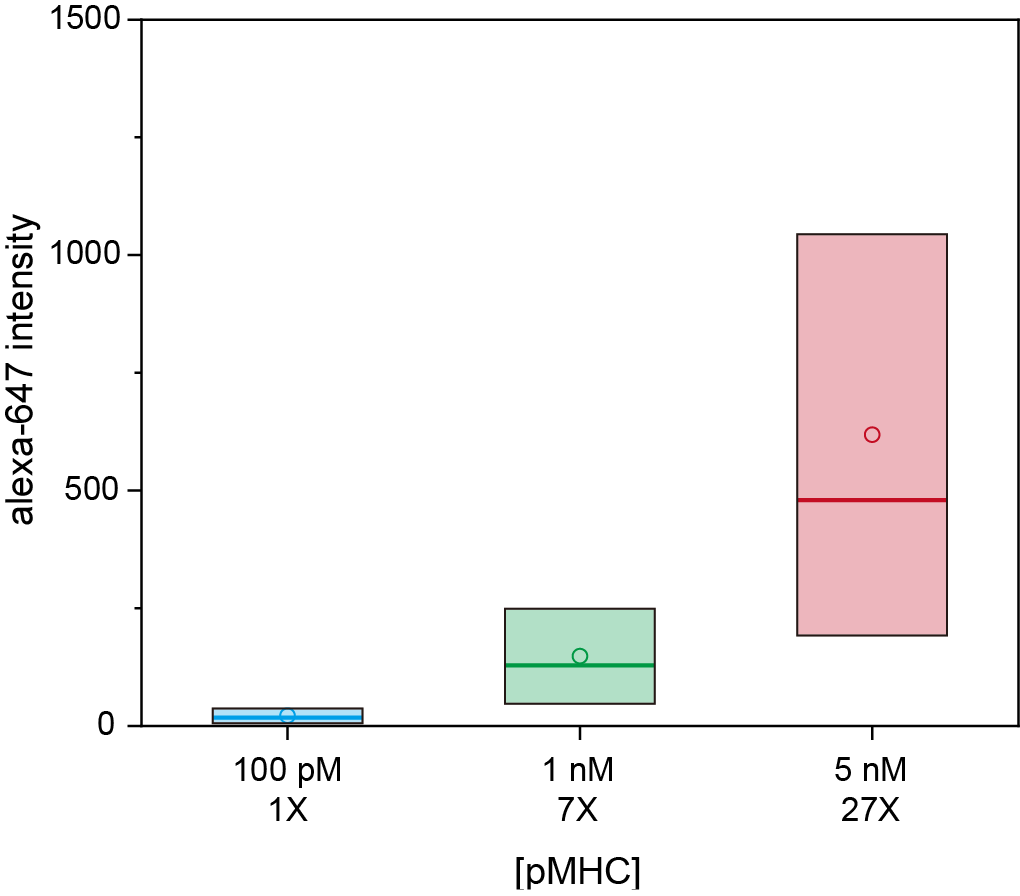

**Figure S27**. Mean (filled circle) and median (line) of Alexa-647 intensities for ‘smart’ beads coated with three different pMHC concentrations (biotin-VPLTEDALL/HLA-B35) used for concentration multiplex experiments; box represents the standard deviation and the line and dot inside indicate the median and mean, respectively. Codes 4 (coated with 1X interfacial pMHC), 14 (coated with ~7X interfacial pMHC) and 15 (coated with ~27X interfacial pMHC) were used and ~100-200 beads were analyzed for each code.

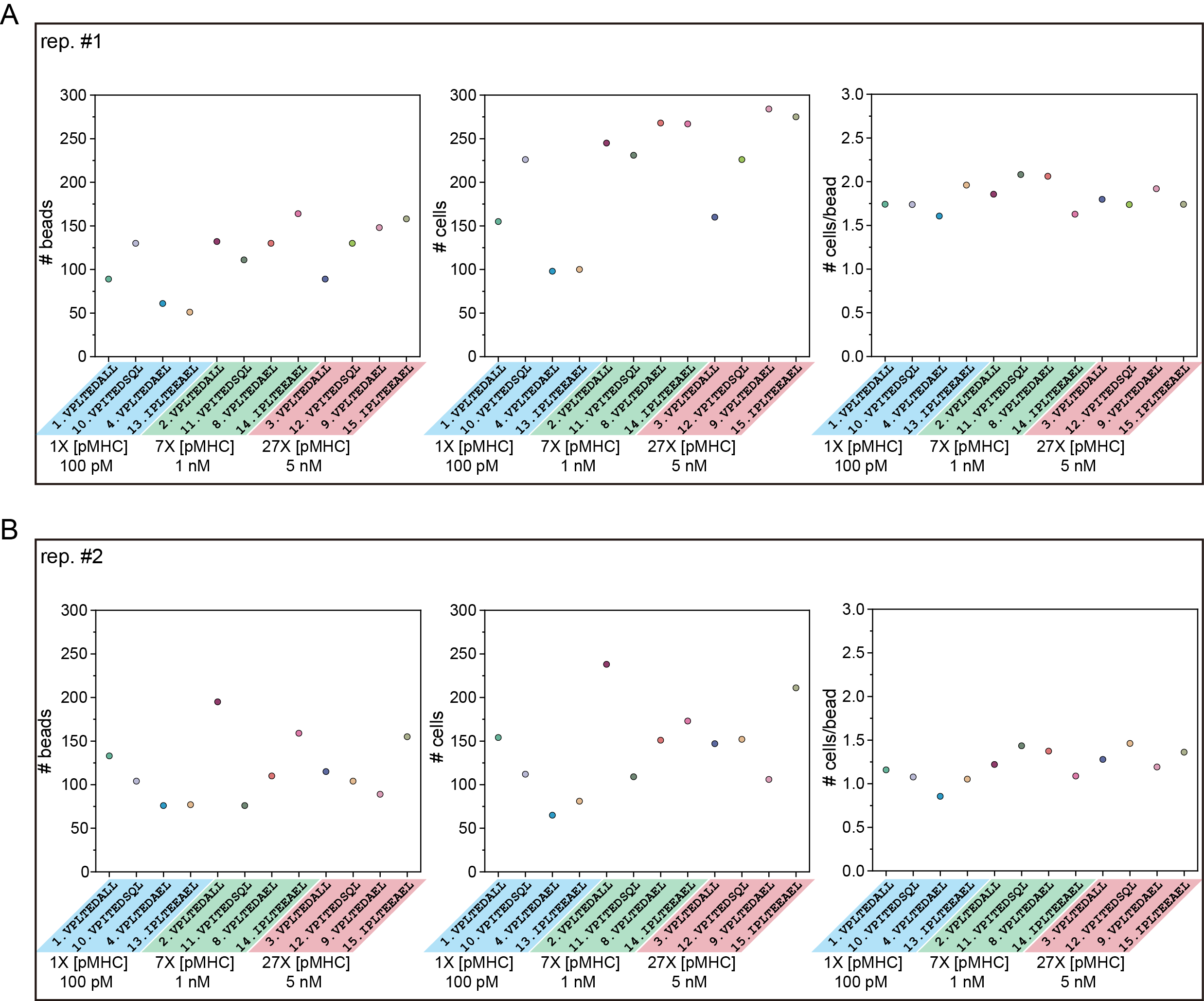

**Figure S28**. No. of loaded beads (left) and cells (middle) per code and cells per bead (right) for two TCR55 concentration multiplex activation experiments. (**A**) Replicate 1. (**B**) Replicate 2.

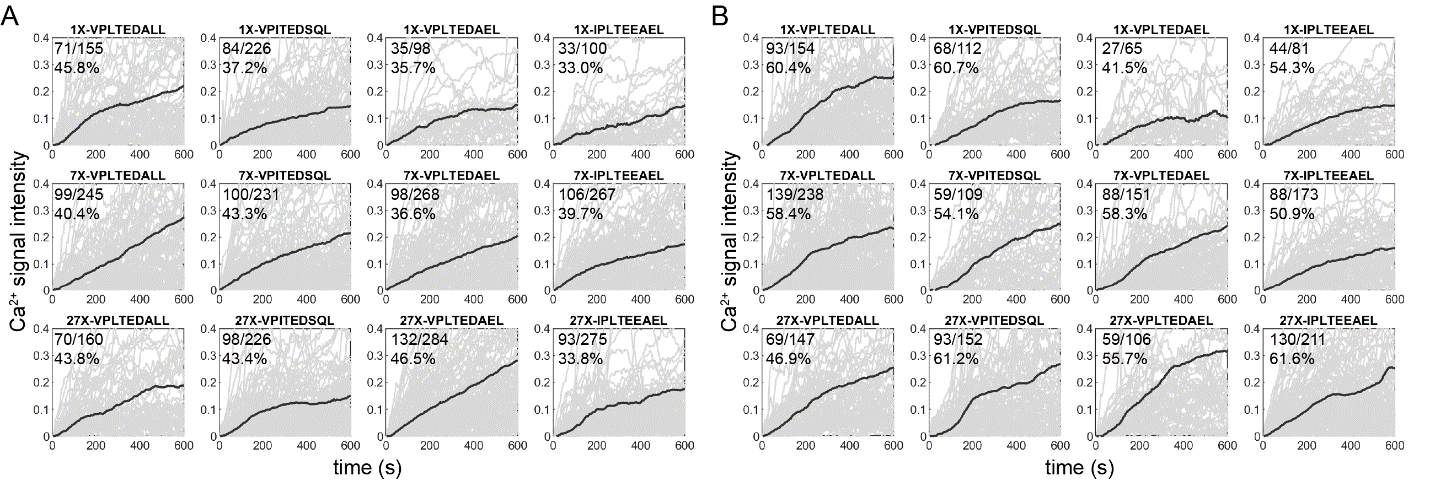

**Figure S29**. Measured positive Ca^2+^-induced fluorescence as a function of time for two replicates of TCR55 concentration multiplex experiments (3 different pMHC concentrations) interacting with 4 different peptide sequences. (**A**) All positive traces with the indicated numbers of positive cells and measured cells and positive percentages for Replicate 1. (**B**) All positive traces with the indicated numbers of positive cells and measured cells and positive percentages for Replicate 2. Light grey lines indicate traces for individual cells; dark grey lines indicate the mean signals.

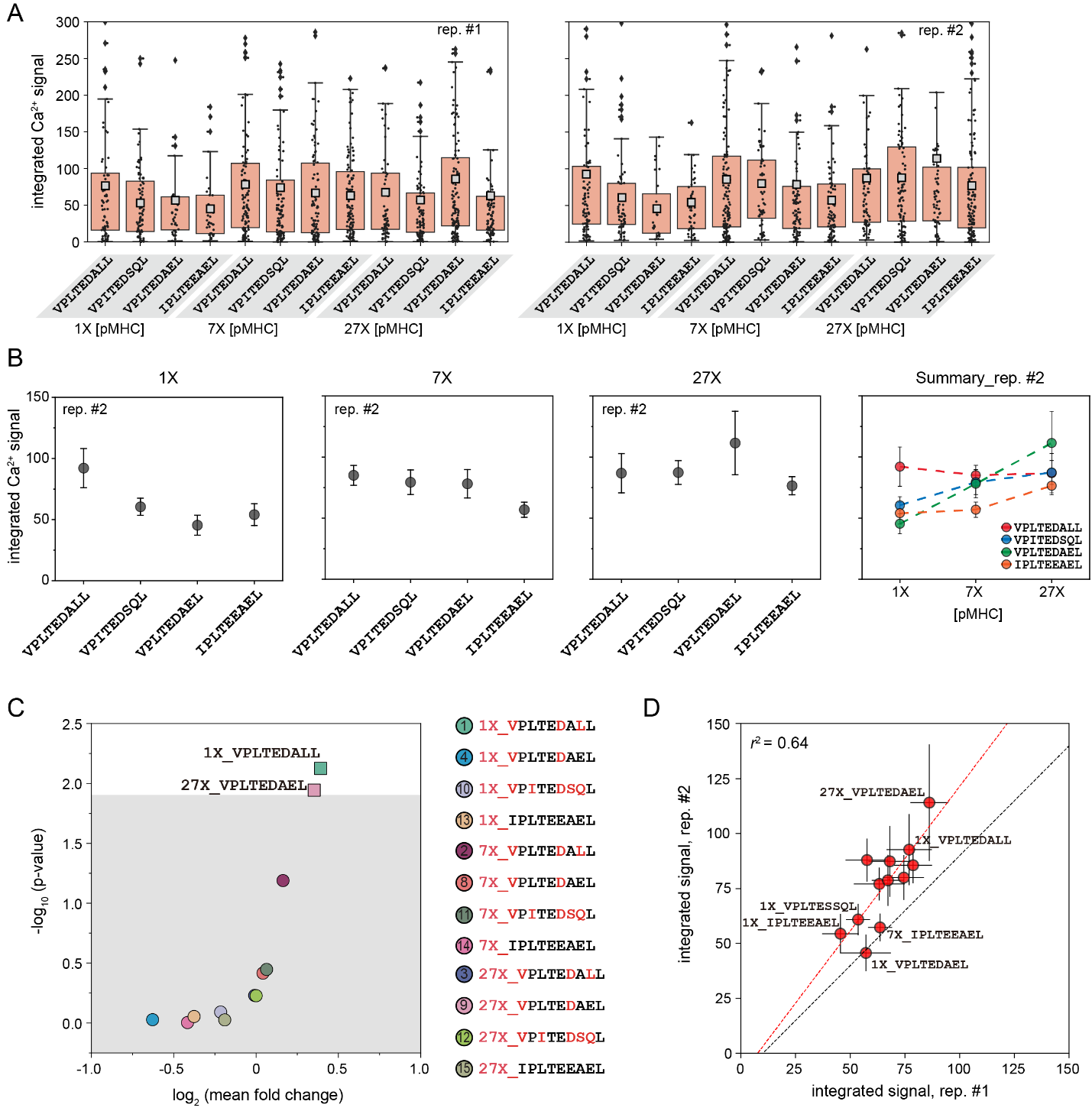

**Figure S30**. Sequence and concentration multiplexed measurements for two TCR55 replicates. (**A**) Measured integrated Ca^2+^ fluorescence signals for each positive cell (light grey markers) and the mean integrated Ca^2+^ signal (light grey square) for two TCR55 replicates interacting with 4 different peptides at three pMHC concentrations. Box lower and upper limits indicate 25^th^ and 75^th^ percentiles, respectively. (**B**) Top row: mean integrated Ca^2+^ signals for positive cells interacting with VPLTEDALL, VPITEDSQL, VPLTEDAEL and IPLTEEAEL peptides at 1X, 7X and 27X pMHC concentrations and the summarized integrated Ca^2+^ signals for four selected peptides under three pMHC concentrations for replicate 2. Error bars indicate SEM. (**C**) Estimated p-value *vs.* log2-transformed mean fold change of integrated Ca^2+^ signals for 4 peptides at 3 different pMHC concentrations; grey area represents p-value>0.013 (Bonferroni-corrected p-value at a significance of 0.05). Correspondence between ‘smart bead’ spectral code and displayed peptide sequence combined its concentration is shown at right. (**D**) Correlation plot of the means of integrated Ca^2+^ signals between two replicates. Dashed black line indicates the 1:1 line; red dashed line indicates a linear regression.

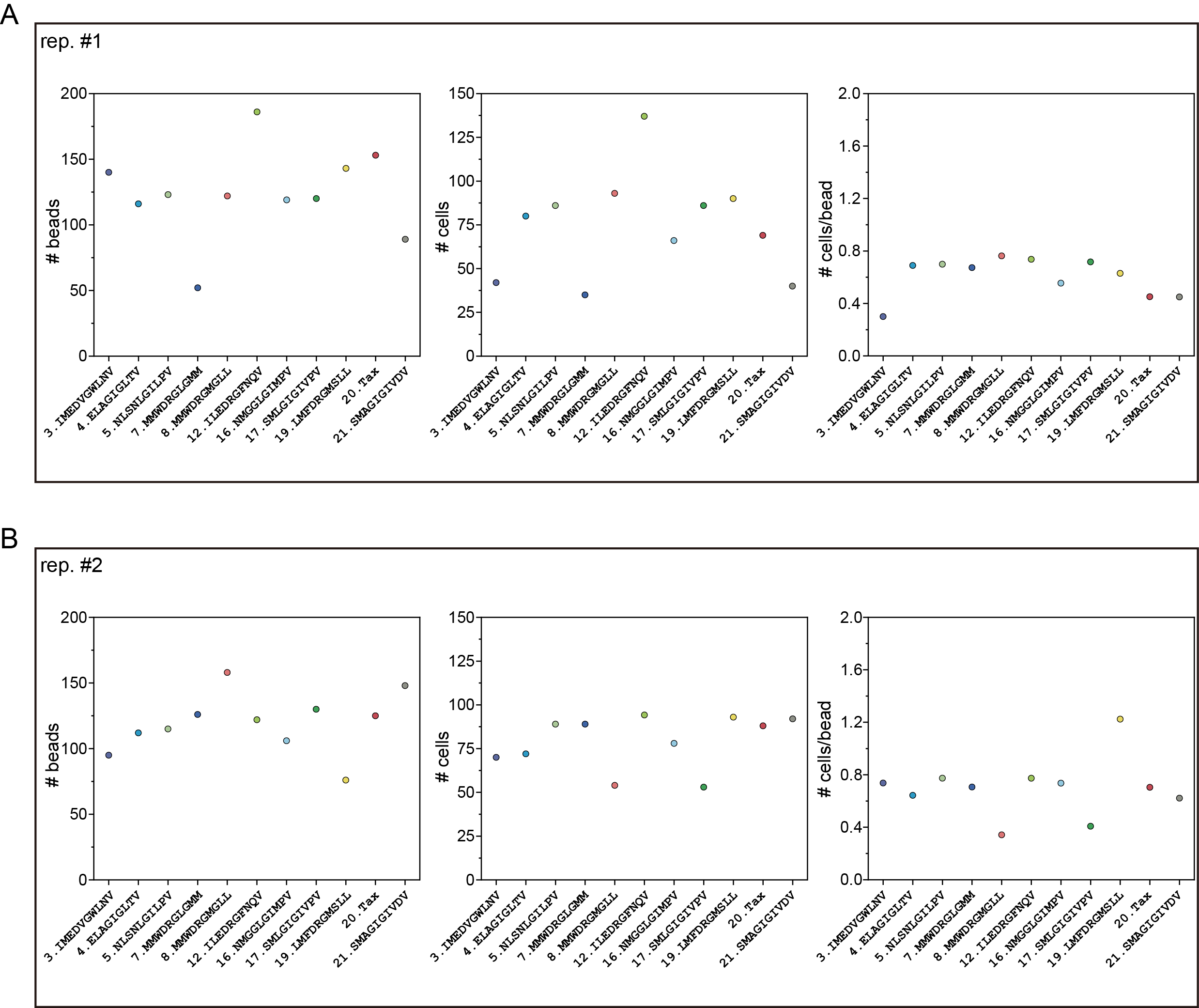

**Figure S31**. No. of loaded beads (left) and cells (middle) per code and cells per bead (right) for two DMF5 T-cell activation experiments. (**A**) Replicate 1. (**B**) Replicate 2.

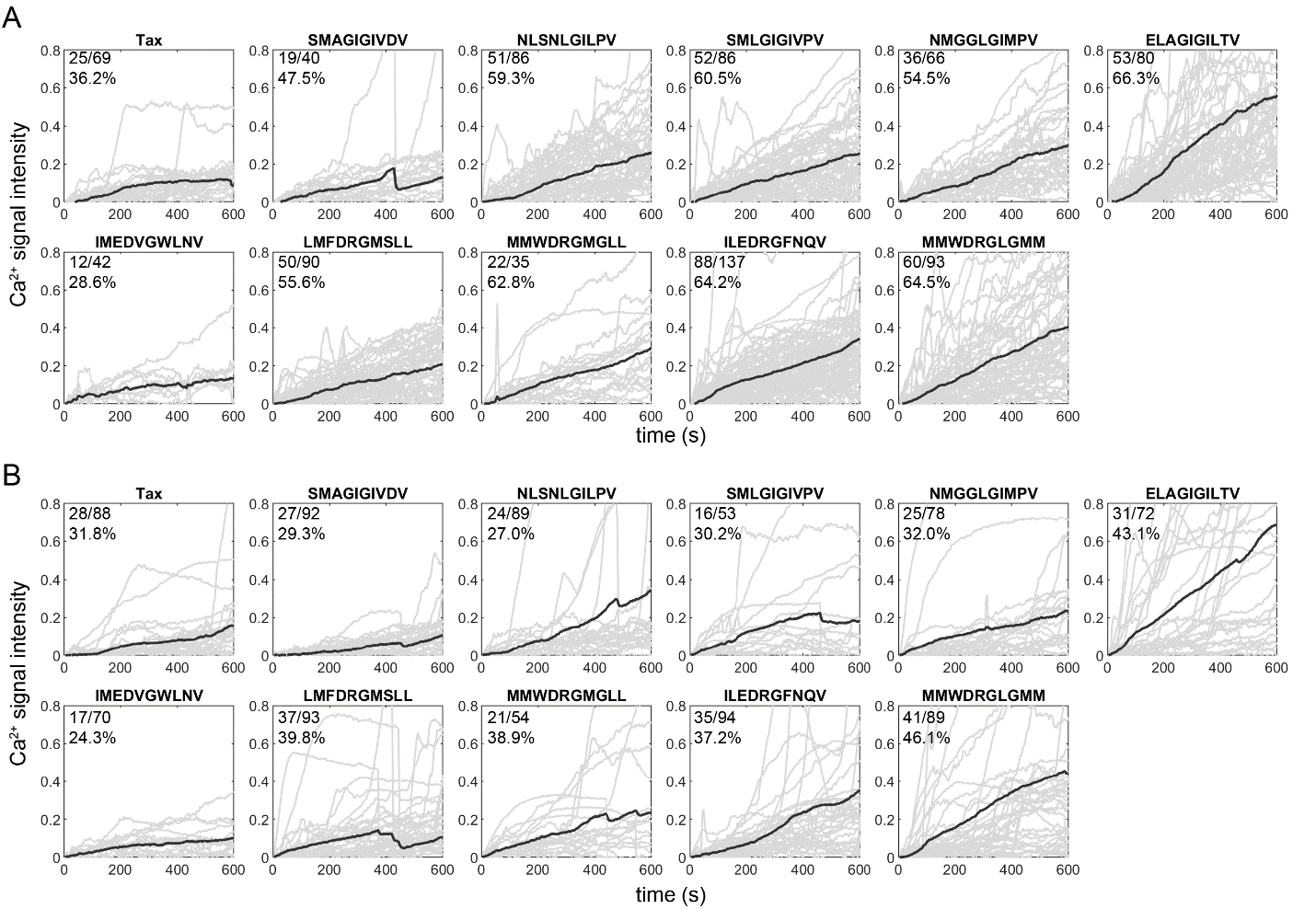

**Figure S32**. Measured positive Ca^2+^-induced fluorescence as a function of time for two replicates of DMF5-transduced T cells interacting with 11 different peptide sequences. (**A**) All positive traces with the indicated numbers of positive cells and measured cells and positive percentages for replicate 1. (**B**) All positive traces with the indicated numbers of positive cells and measured cells and positive percentages for replicate 2. Light grey lines indicate traces for individual cells; dark grey lines indicate the mean signals.

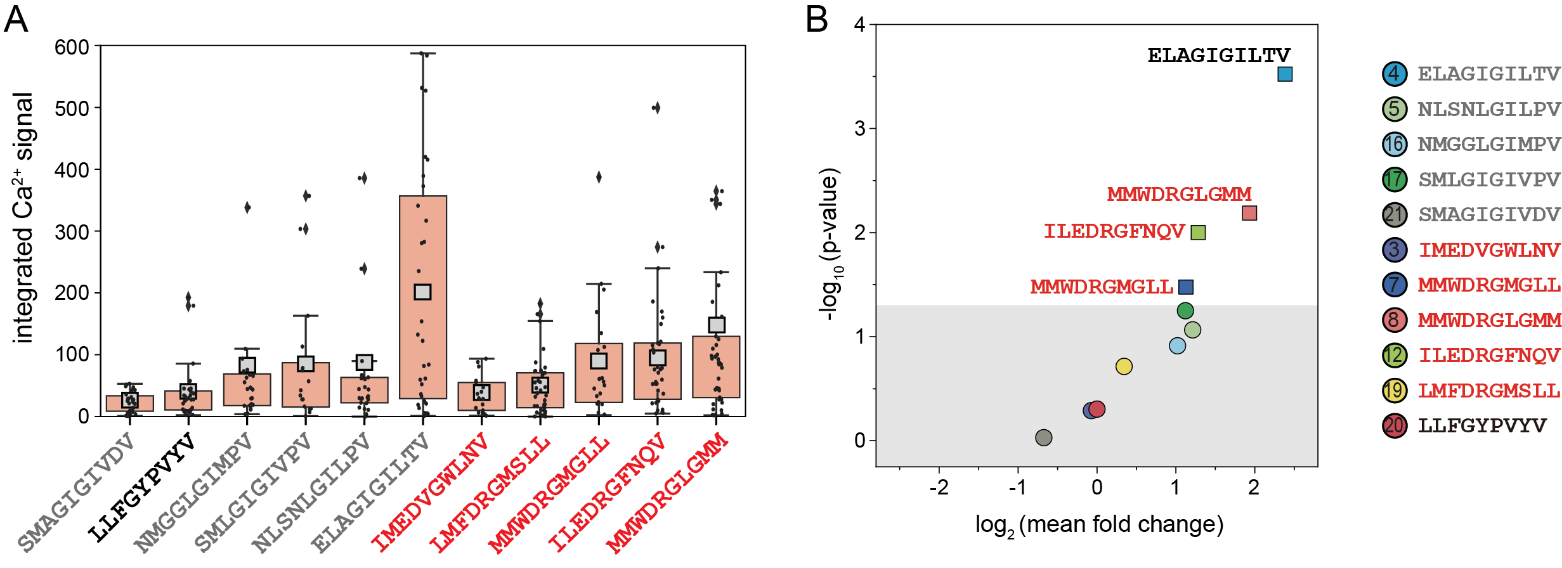

**Figure S33**. Measured integrated Ca^2+^ fluorescence signals and estimated significance for DMF5 TCR replicate 2. (**A**) Integrated Ca^2+^ signals for each positive cell (light grey markers) and the mean integrated Ca^2+^ signal (light grey square). Box lower and upper limits indicate 25^th^ and 75^th^ percentiles, respectively. (**B**) Estimated p-value *vs.* log2-transformed mean fold change for DMF5-tranduced T cells interacting with 11 different peptide sequences. Grey area represents p-value>0.05. Correspondence between ‘smart bead’ spectral code and displayed peptide sequence is shown at right.

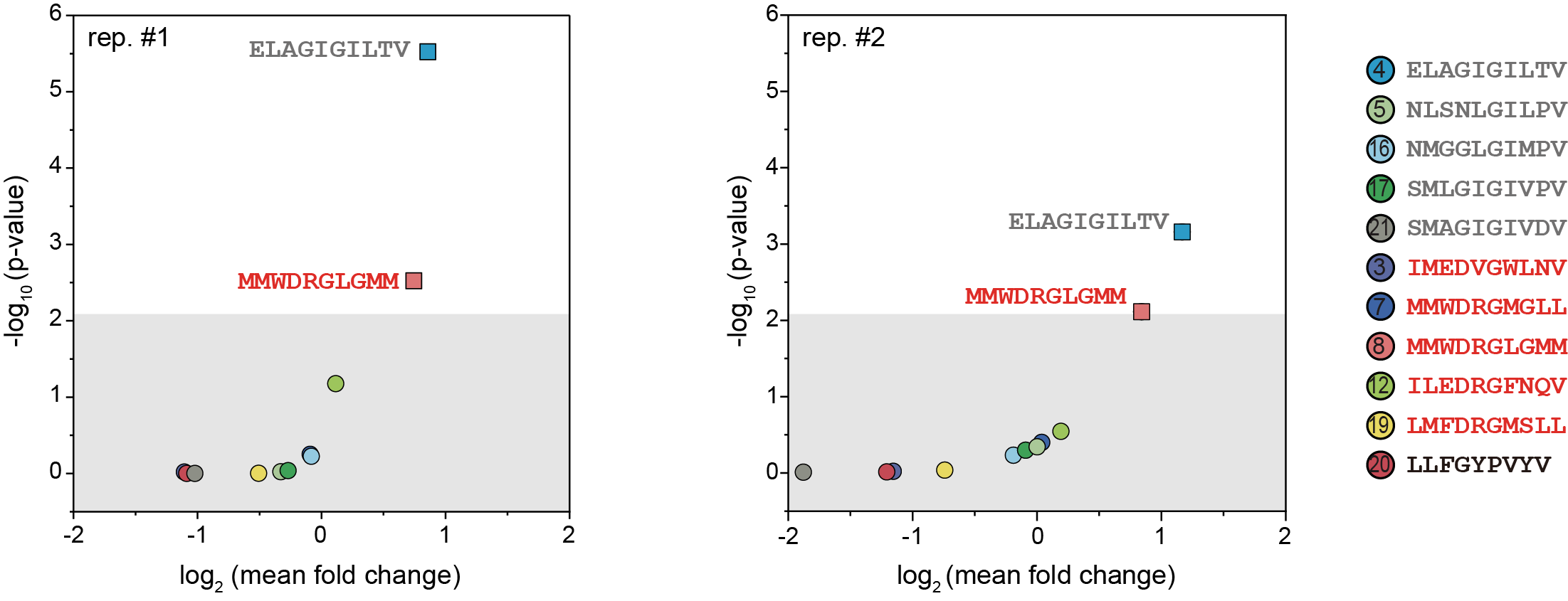

**Figure S34**. Estimated p-value *vs.* log2-transformed mean fold change for DMF5-tranduced T cells interacting with 11 different peptide sequences for replicate 1 (left) and replicate 2 (right). Grey area represents p-value>0.008 (Bonferroni-corrected p-value at a significance of 0.05). Correspondence between ‘smart bead’ spectral code and displayed peptide sequence is shown at right.

**Table S1**. Volumetric ratios of Lns master mix used in BATTLES for fabricating ‘smart beads’ with 21 codes.

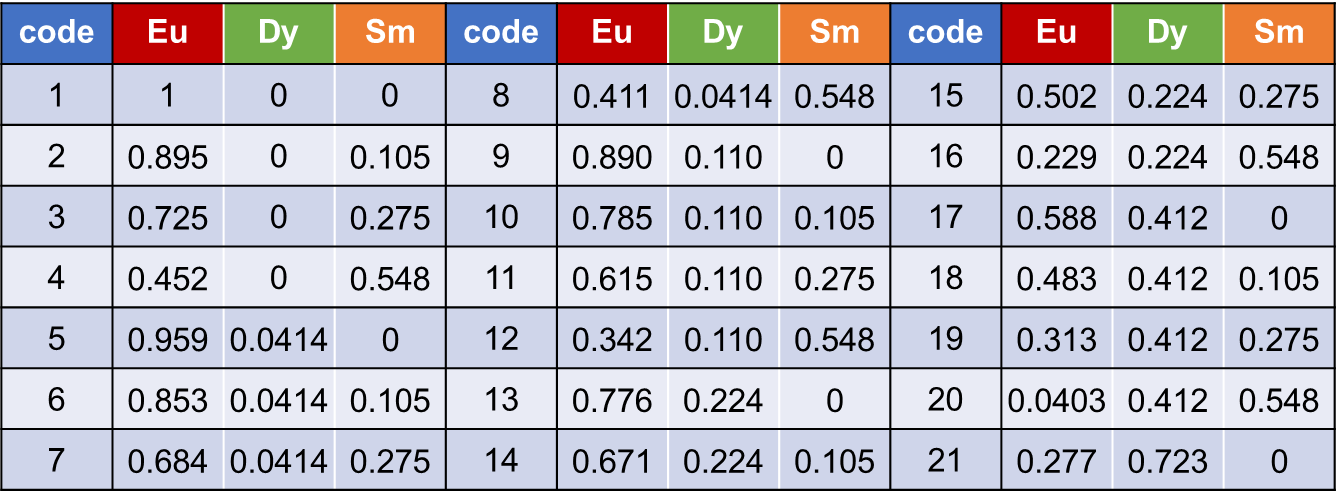

**Table S2**. Relationship between spectral codes and peptide sequences in TCR589 and TCR55 experiments.

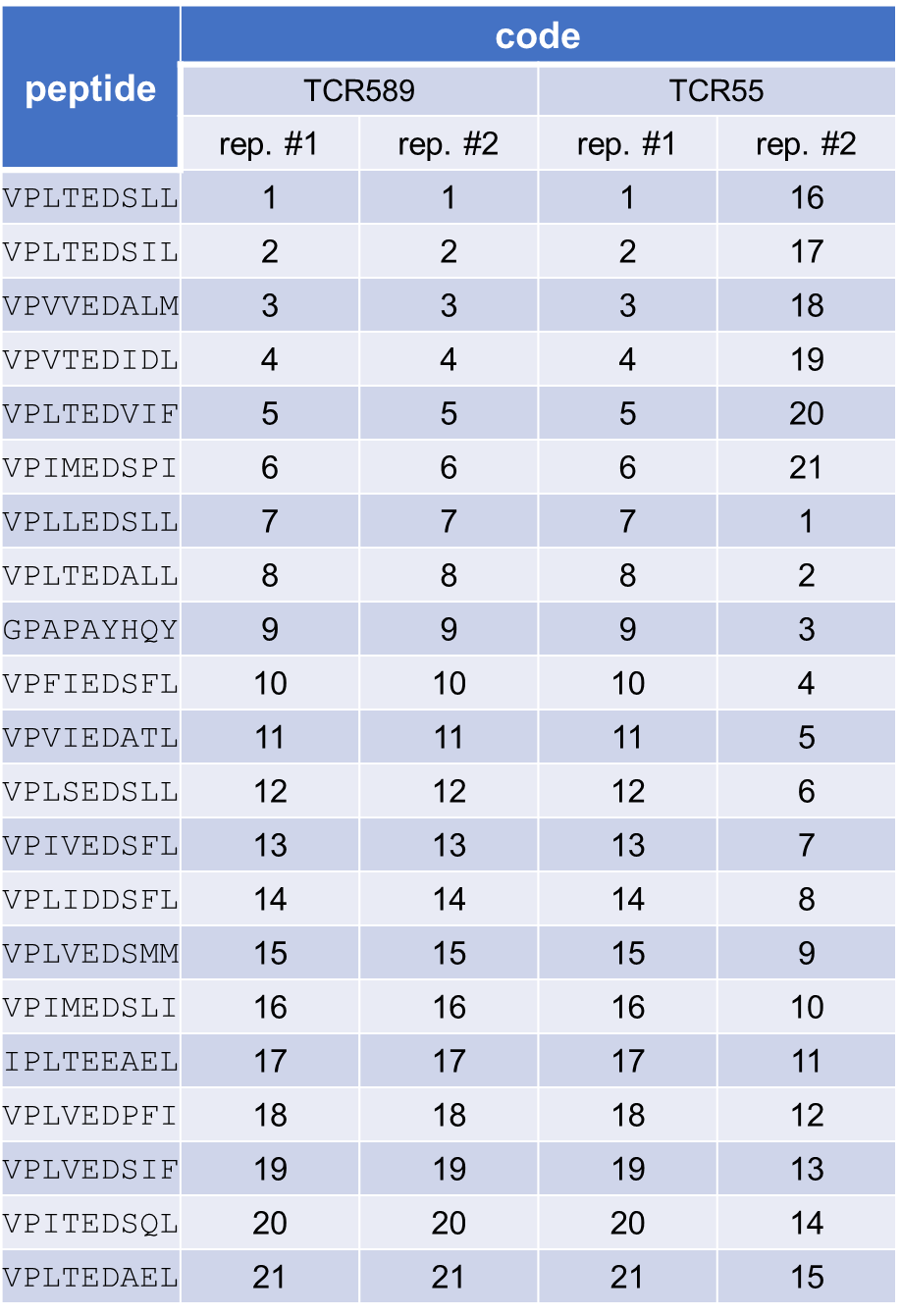

**Table S3**. Ec_50_ values for different peptides used in TCR55-APC coculture experiment. Data was fit from ref. 2.

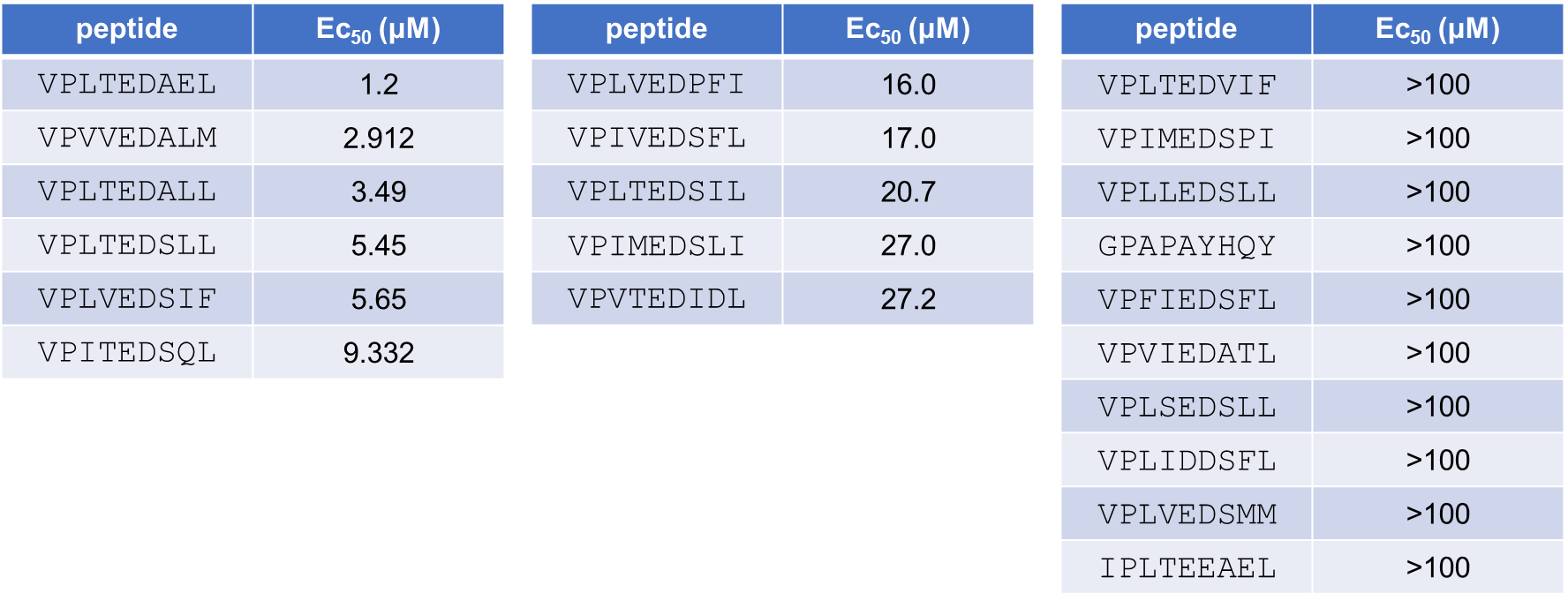

**Table S4**. Relationship between spectral codes and peptide sequences in TCR55 force multiplex experiments.

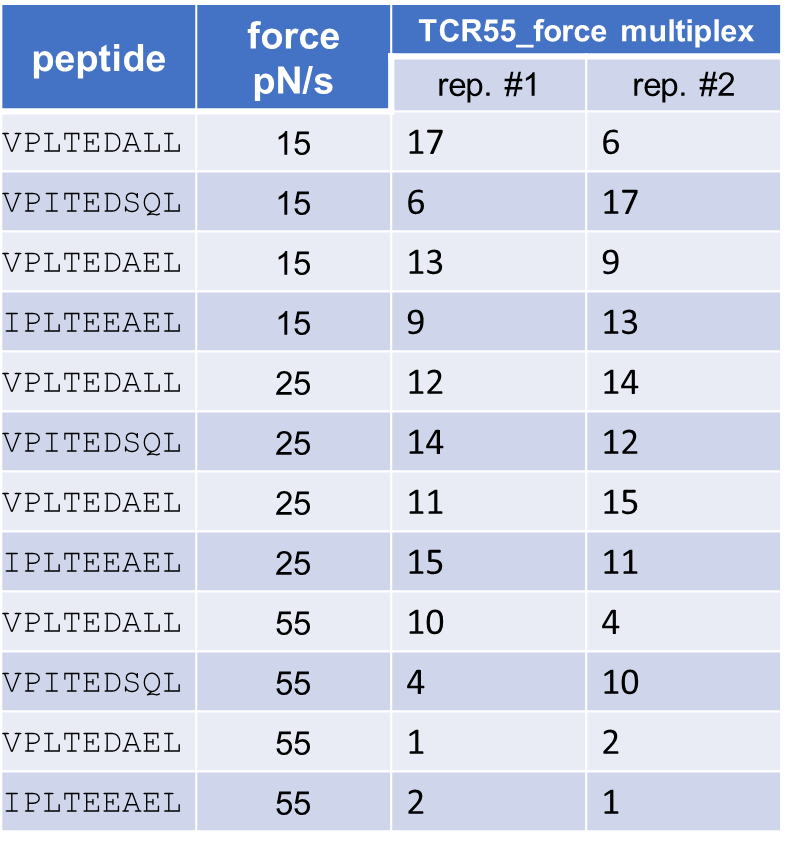

**Table S5**. Relationship between spectral codes and peptide sequences in TCR55 concentration multiplex experiments.

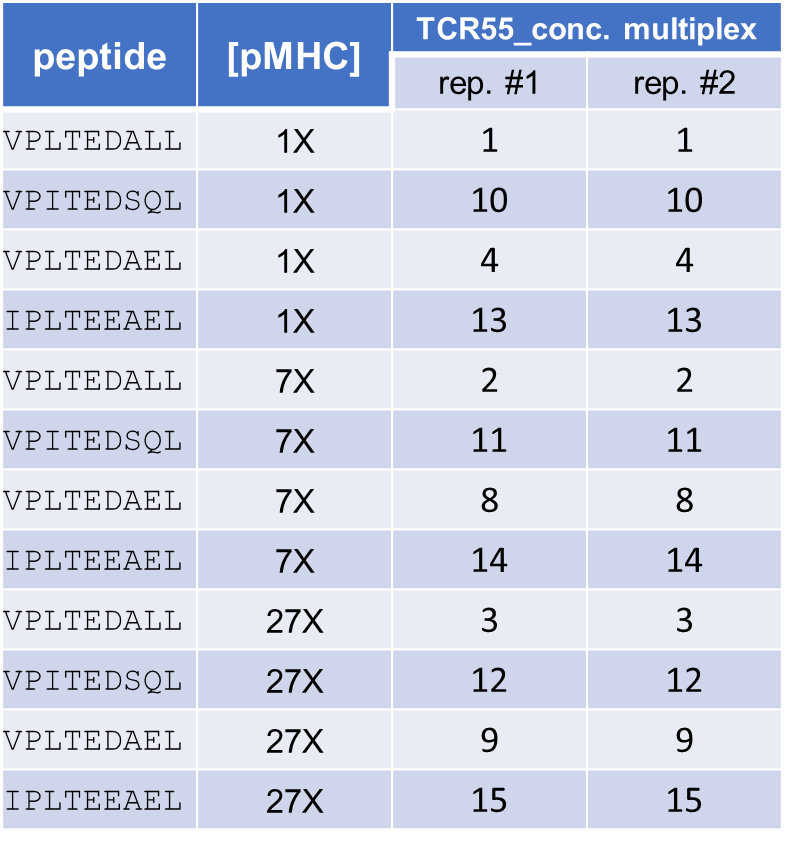

**Table S6**. Relationship between spectral codes and peptide sequences in DMF5 TCR experiments. DRG and GIG classes are in red and black. Bold sequence is Tax (LLFGYPVYV).

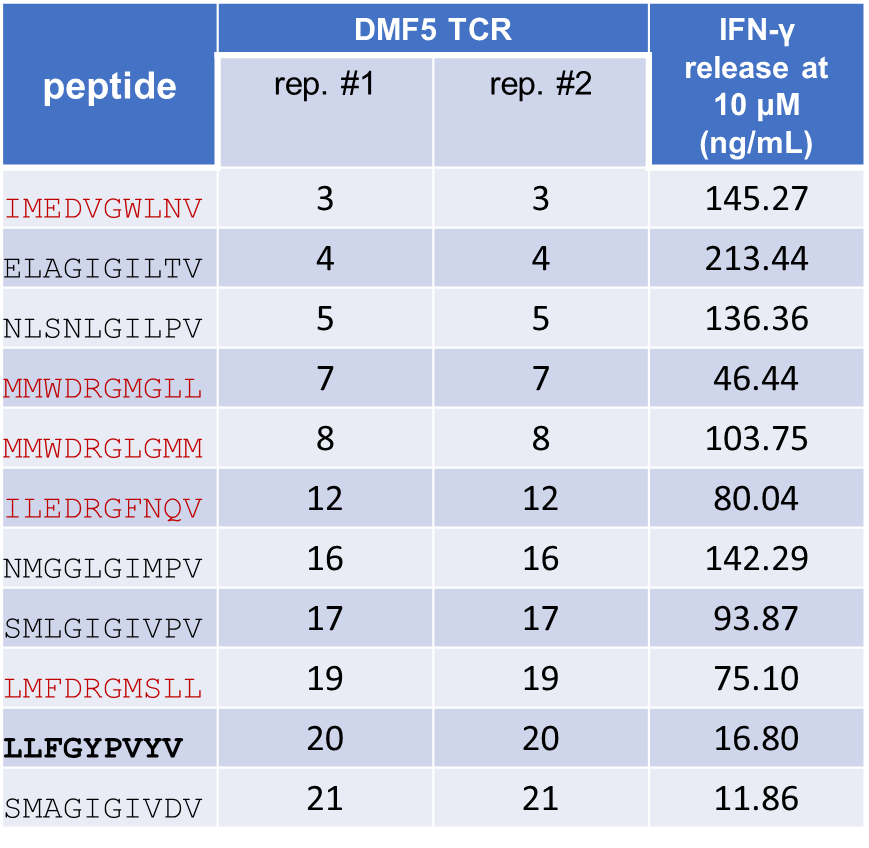

**Table S7**. Existing mechanobiology tools used for profiling T cell-pMHC interactions.

| force type | tech. | reference | binding or activation? | pMHC density | no. of test peptides | no. of test cells |
| --- | --- | --- | --- | --- | --- | --- |
| passive force | supported lipid bilayer | reviewed in (Groves and Dustin, 2003) | both | 6-175 molecules/μm^2^ | usually 1 peptide per assay but with the potential to multiple pMHCs using the arrays of nanowells (Torres et al., 2013) | 100-300 of cells per field of view |
|  | hydrogel surface | (Guasch et al., 2017), (Colin-York et al., 2019) and (Chin et al., 2020) | activation | using anti-CD3 antibodies(Chin et al., 2020; Guasch et al., 2017) or gel surface incubated with ~2 μM biotin-pMHCs (Colin-York et al., 2019) | not available | <100 cells per field of view or 1000 cells using the ELISA method |
|  | micropillar | (Hui et al., 2015), (Bashour et al., 2014), (Basu et al., 2016) and (Hu et al., 2016) | activation | not available | 1 peptide per assay (Basu et al., 2016) or using anti-CD3 antibodies (Bashour et al., 2014; Hu et al., 2016; Hui et al., 2015) | ~200 cells per surface |
|  | molecular tension probes (e.g. peptide or DNA) | reviewed in (Liu et al., 2017) | both | >30 pMHC/ μm^2^ | 1 peptide per assay | 20 cells per field of view |
| active force | optical trap | (Brazin et al., 2018; Das et al., 2015; Das et al., 2016; Feng et al., 2017) | both | as low as 2 pMHC molecules per cell for activation experiment and 1 pMHC per cell for binding assay | 1 peptide per assay | <10 cells per assay |
|  | biomembrane force probe | (Hong et al., 2018; Huang et al., 2010; Husson et al., 2011; Jiang et al., 2011; Li et al., 2010; Liu et al., 2014) | both | >40 pMHC/ μm^2^ | 1 peptide per assay | 1 cell per pulling assay |
|  | atomic force microscopy | (Hu and Butte, 2016; Puech et al., 2011) | both | Streptavidin functionalized AFM cantilever was incubated with ~1 μM biotin-pMHCs | 1 peptide per assay | 1 cell per pulling assay |
|  | optomechanical actuator (OMA) nanoparticle | (Liu et al., 2016) | activation | OMA surface incubated with ~8 μM biotin-pMHCs | 1 peptide per assay | <10 cells per field of view |
|  | microfluidics | (Stockslager et al., 2017) | binding | ∼4 × 10^4^ pMHC μm^−2^ | 1 peptide per assay | < 50 cells per assay |
|  | laminar flow chamber | (Robert et al., 2012) | binding | pMHC coating densities of 0.5 *μ*g/mL to 0.1 to 0.2 μM | 1 peptide per assay | with the potential to test multiple cells |
|  | acoustic* | (Pan et al., 2018) | activation | not available | not available | <50 cells per assay |

*This technology was applied to the CAR-T cell system with a microbubble to transduce the force to the cells engineered with the mechanosensor Piezo1 and genetic transducing modules.

**Supplementary Information References**

Bashour, K.T., Gondarenko, A., Chen, H., Shen, K., Liu, X., Huse, M., Hone, J.C., and Kam, L.C. (2014). CD28 and CD3 have complementary roles in T-cell traction forces. Proceedings of the National Academy of Sciences *111*, 2241-2246.

Basu, R., Whitlock, B.M., Husson, J., Le Floc’h, A., Jin, W., Oyler-Yaniv, A., Dotiwala, F., Giannone, G., Hivroz, C., and Biais, N. (2016). Cytotoxic T cells use mechanical force to potentiate target cell killing. Cell *165*, 100-110.

Bell, G.I. (1978). Models for the specific adhesion of cells to cells. Science *200*, 618-627.

Brazin, K.N., Mallis, R.J., Boeszoermenyi, A., Feng, Y., Yoshizawa, A., Reche, P.A., Kaur, P., Bi, K., Hussey, R.E., Duke-Cohan, J.S.*, et al.* (2018). The T Cell Antigen Receptor α Transmembrane Domain Coordinates Triggering through Regulation of Bilayer Immersion and CD3 Subunit Associations. Immunity *49*, 829-841.e826.

Chin, M.H.W., Norman, M.D.A., Gentleman, E., Coppens, M.-O., and Day, R.M. (2020). A Hydrogel-Integrated Culture Device to Interrogate T Cell Activation with Physicochemical Cues. ACS Applied Materials & Interfaces *12*, 47355-47367.

Colin-York, H., Javanmardi, Y., Skamrahl, M., Kumari, S., Chang, V.T., Khuon, S., Taylor, A., Chew, T.-L., Betzig, E., Moeendarbary, E.*, et al.* (2019). Cytoskeletal Control of Antigen-Dependent T Cell Activation. Cell Reports *26*, 3369-3379.e3365.

Das, D.K., Feng, Y., Mallis, R.J., Li, X., Keskin, D.B., Hussey, R.E., Brady, S.K., Wang, J.-H., Wagner, G., Reinherz, E.L.*, et al.* (2015). Force-dependent transition in the T-cell receptor β-subunit allosterically regulates peptide discrimination and pMHC bond lifetime. Proceedings of the National Academy of Sciences *112*, 1517-1522.

Das, D.K., Mallis, R.J., Duke-Cohan, J.S., Hussey, R.E., Tetteh, P.W., Hilton, M., Wagner, G., Lang, M.J., and Reinherz, E.L. (2016). Pre-T Cell Receptors (Pre-TCRs) Leverage Vβ Complementarity Determining Regions (CDRs) and Hydrophobic Patch in Mechanosensing Thymic Self-ligands. Journal of Biological Chemistry *291*, 25292-25305.

Feng, Y., Brazin, K.N., Kobayashi, E., Mallis, R.J., Reinherz, E.L., and Lang, M.J. (2017). Mechanosensing drives acuity of <em>αβ</em> T-cell recognition. Proceedings of the National Academy of Sciences *114*, E8204-E8213.

Groves, J.T., and Dustin, M.L. (2003). Supported planar bilayers in studies on immune cell adhesion and communication. Journal of Immunological Methods *278*, 19-32.

Guasch, J., Muth, C.A., Diemer, J., Riahinezhad, H., and Spatz, J.P. (2017). Integrin-Assisted T-Cell Activation on Nanostructured Hydrogels. Nano Letters *17*, 6110-6116.

Hong, J., Ge, C., Jothikumar, P., Yuan, Z., Liu, B., Bai, K., Li, K., Rittase, W., Shinzawa, M., Zhang, Y.*, et al.* (2018). A TCR mechanotransduction signaling loop induces negative selection in the thymus. Nature Immunology *19*, 1379-1390.

Hu, J., Gondarenko, A.A., Dang, A.P., Bashour, K.T., O’Connor, R.S., Lee, S., Liapis, A., Ghassemi, S., Milone, M.C., Sheetz, M.P.*, et al.* (2016). High-Throughput Mechanobiology Screening Platform Using Micro- and Nanotopography. Nano Letters *16*, 2198-2204.

Hu, K.H., and Butte, M.J. (2016). T cell activation requires force generation. Journal of Cell Biology *213*, 535-542.

Huang, J., Zarnitsyna, V.I., Liu, B., Edwards, L.J., Jiang, N., Evavold, B.D., and Zhu, C. (2010). The kinetics of two-dimensional TCR and pMHC interactions determine T-cell responsiveness. Nature *464*, 932-936.

Hui, K.L., Balagopalan, L., Samelson, L.E., and Upadhyaya, A. (2015). Cytoskeletal forces during signaling activation in Jurkat T-cells. Molecular biology of the cell *26*, 685-695.

Husson, J., Chemin, K., Bohineust, A., Hivroz, C., and Henry, N. (2011). Force generation upon T cell receptor engagement. PloS one *6*, e19680.

Jiang, N., Huang, J., Edwards, L.J., Liu, B., Zhang, Y., Beal, C.D., Evavold, B.D., and Zhu, C. (2011). Two-Stage Cooperative T Cell Receptor-Peptide Major Histocompatibility Complex-CD8 Trimolecular Interactions Amplify Antigen Discrimination. Immunity *34*, 13-23.

Li, Y.-C., Chen, B.-M., Wu, P.-C., Cheng, T.-L., Kao, L.-S., Tao, M.-H., Lieber, A., and Roffler, S.R. (2010). Cutting Edge: Mechanical Forces Acting on T Cells Immobilized via the TCR Complex Can Trigger TCR Signaling. The Journal of Immunology *184*, 5959-5963.

Liu, B., Chen, W., Evavold, Brian D., and Zhu, C. (2014). Accumulation of Dynamic Catch Bonds between TCR and Agonist Peptide-MHC Triggers T Cell Signaling. Cell *157*, 357-368.

Liu, Y., Galior, K., Ma, V.P.-Y., and Salaita, K. (2017). Molecular Tension Probes for Imaging Forces at the Cell Surface. Accounts of Chemical Research *50*, 2915-2924.

Liu, Z., Liu, Y., Chang, Y., Seyf, H.R., Henry, A., Mattheyses, A.L., Yehl, K., Zhang, Y., Huang, Z., and Salaita, K. (2016). Nanoscale optomechanical actuators for controlling mechanotransduction in living cells. Nature Methods *13*, 143-146.

Pan, Y., Yoon, S., Sun, J., Huang, Z., Lee, C., Allen, M., Wu, Y., Chang, Y.-J., Sadelain, M., Shung, K.K.*, et al.* (2018). Mechanogenetics for the remote and noninvasive control of cancer immunotherapy. Proceedings of the National Academy of Sciences *115*, 992-997.

Puech, P.-H., Nevoltris, D., Robert, P., Limozin, L., Boyer, C., and Bongrand, P. (2011). Force Measurements of TCR/pMHC Recognition at T Cell Surface. PLOS ONE *6*, e22344.

Robert, P., Aleksic, M., Dushek, O., Cerundolo, V., Bongrand, P., and van der Merwe, P.A. (2012). Kinetics and Mechanics of Two-Dimensional Interactions between T Cell Receptors and Different Activating Ligands. Biophysical Journal *102*, 248-257.

Stockslager, M.A., Bagnall, J.S., Hecht, V.C., Hu, K., Aranda-Michel, E., Payer, K., Kimmerling, R.J., and Manalis, S.R. (2017). Microfluidic platform for characterizing TCR-pMHC interactions. Biomicrofluidics *11*, 064103-064103.

Torres, A.J., Contento, R.L., Gordo, S., Wucherpfennig, K.W., and Love, J.C. (2013). Functional single-cell analysis of T-cell activation by supported lipid bilayer-tethered ligands on arrays of nanowells. Lab on a chip *13*, 90-99.
