## Supplementary Movie Captions for "Structure-activity mapping of the peptide- and force-dependent landscape of T-cell activation"

**Movie S1**. Parallel flow focuser for production of ‘smart beads’ with linear amplification (4 flow focusing channels).

**Movie S2**. Thermo-response of the ‘smart beads’ with 55 mM sodium acrylate during the heating and cooling process. Code 9 was used in this movie.

**Movie S3**. Initial bead loading followed by cell loading prior to on-chip imaging.

**Movie S4**. T cells remained in contact with bead surfaces throughout heating and cooling process. 4 representative wells are presented.

**Movie S5**. Representative Ca^2+^ flux within single TCR589 T cells interacting with pMHC-coated ‘smart beads’ bearing the stimulatory HIVpol peptide (IPLTEEAEL) and the nonstimulatory peptide (VPLTEDAEL).

**Movie S6**. Thermo-response of the ‘smart beads’ with 50 mM (code 1), 55 mM (code 15) and 60 mM (code 9) sodium acrylate during the heating and cooling process.
